# “A Phosphorylation-Induced Micellization switch in the low complexity domain of TDP-43”

**DOI:** 10.64898/2026.06.09.730863

**Authors:** Rodrigo F. Dillenburg, Anastasia Lopatina, Hao Ruan, Tom Scheidt, Simone Mosna, Emre Pekbilir, Julia Bieber, Frank Schäfer-Depoix, Katharina Landfester, Carla Schmidt, Martin M. Möckel, Svenja Morsbach, Friederike Schmid, Dorothee Dormann, Lukas Stelzl, Martin Girard, Edward A. Lemke

## Abstract

Phase separation (PS) of the low-complexity domain (LCD) of TDP-43 is linked to pathogenic aggregates in amyotrophic lateral sclerosis (ALS) and frontotemporal lobar degeneration (FTLD-TDP). Here, we show that extensive phosphorylation of the LCD C-terminus redirects its self-assembly. Coarse-grained Monte Carlo simulations predicted that 12 Ser phosphorylations partition the 148-residue LCD into a hydrophobic N-terminal and highly charged C-terminal block, favoring finite-sized micellization over macroscopic PS. *In vitro,* LCD phosphorylated by casein kinase 1 delta (CK1δ; mean of 12 phosphorylations by native mass spectrometry) and phosphomimetic 12D/12DD mutants formed spherical nanoparticles (≈ 20-50 nm) above a low-micromolar critical micelle concentration, whereas the unphosphorylated LCD underwent reversible PS that matured into fibrils. Increasing ionic strength shifted the mutants toward anisotropic morphologies (worm-like 12D micelles and rigid 12DD nanocylinders). Turbidity assays and confocal imaging directly visualized the absence of PS in the phosphorylated form. Negative-stain and cryo-EM confirmed the spherical micellar architecture for the phosphorylated LCD and 12D/12DD mimics. Our data identify phosphorylation as a molecular switch tuning macrophase separation and fibril formation of TDP-43 LCD, providing a framework for an aggregation-protective role through microphase separation into size-limited micelles. Whether these assemblies are stable or kinetically trapped on pathological timescales remains unclear.

## 1. Introduction

TAR DNA-binding protein 43 kDa (TDP-43) is an abundant, ubiquitously expressed nuclear RNA-binding protein.[1] Under normal conditions, it is predominantly localised in the nucleus, where it participates in the regulation of transcription, splicing and miRNA biogenesis. It is capable of actively shuttling to the cytoplasm[2], where it regulates mRNA transport, mRNA stability and stress granule formation.[1] TDP-43 possesses a classical modular architecture (**Figure 1**), including an N-terminal oligomerization domain (NTD), a nuclear localization signal (NLS) and two RNA-recognition motifs (RRM1 and RRM2). Its disordered C-terminal low-complexity domain (LCD, residues 267-414), rich in glycine, glutamine/asparagine, and serine, bears features of a prion-like region, and plays a central role in self-assembly of the protein and in the pathogenesis of neurodegenerative diseases.^[3, 4]^ In post-mortem brains of amyotrophic lateral sclerosis (ALS) and frontotemporal dementia (FTD) patients, as well as 60% of Alzheimer’s disease (AD) patients, TDP-43 is consistently found mislocalized to the cytoplasm, where it accumulates in hyperphosphorylated, ubiquitinated, detergent-insoluble inclusions.[5,6] These aggregates have an amyloid-like core, as recently determined by cryo-electron microscopy [7,8], but were reported to contain a mixture of granular (amorphous) and fibrillar material [9], and mixtures of full-length TDP-43 and C-terminal fragments (CTFs) of 20-25 kDa comprised mostly of the LCD.^[10, 11]^ TDP-43 aggregates are thought to be pathological and cause neurotoxicity by inducing a nuclear TDP-43 loss of function, thus causing detrimental changes in RNA processing.^[12, 13, 14]^ Understanding the pathways that lead to TDP-43 aggregation is important for developing therapeutic strategies to target the build-up of these toxic protein inclusions.

**Figure 1.**
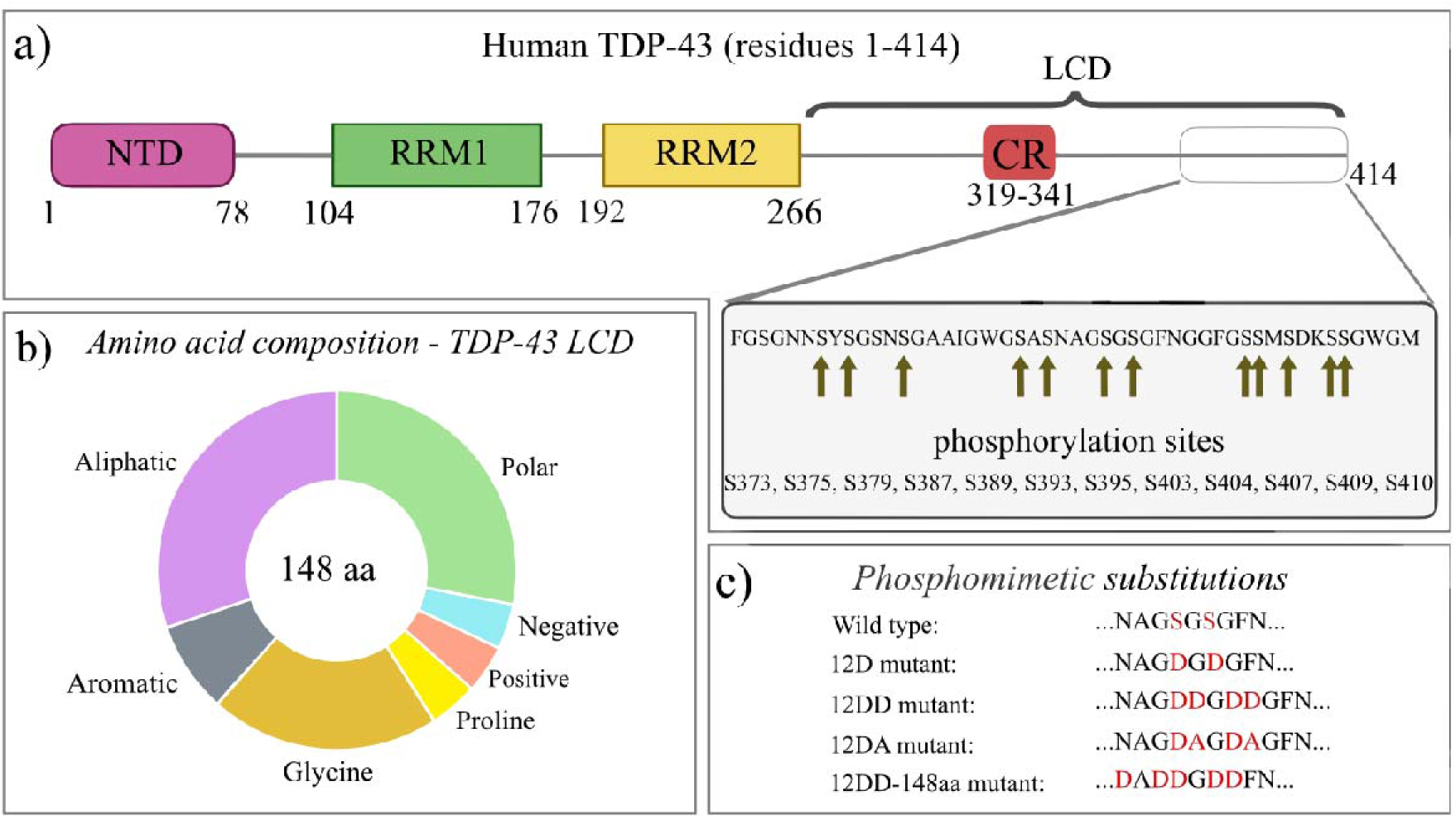
Full-length TDP-43 sequence. **a)** Representation of TDP-43 domains. The last 48 residues of the low complexity domain (LCD) are shown. The location of the phosphorylation sites relevant to this work are indicated by arrows. **b)** Amino acid composition of TDP-43 LCD. **c)** Schematic representation of the phosphomimetic substitution patterns used in this study, illustrated for a representative C-terminal sequence segment. In the wild-type sequence, the serine residues targeted for phosphorylation are highlighted in red. In the **12D** mutant, each of the 12 target serines is replaced by a single aspartate (S→D), preserving the wild-type LCD length of 148 residues and introducing a C-terminal block charge of −12e. In the **12DD** mutant, each serine is replaced by two aspartates (S→DD), increasing the C-terminal block charge to −24e but extending the chain by 12 residues (160 residues total). Two additional control constructs were designed to disentangle charge from chain-length effects: the **12DA** mutant, a length-matched analogue of 12DD (160 residues) in which one aspartate of each pair is replaced by alanine to restore the −12e charge of 12D while retaining the longer backbone; and the **12DD-148aa** mutant, a length-matched analogue of 12D in which native residues adjacent to the substitution sites are removed to restore the wild-type LCD length of 148 residues while preserving the −24e charge. Substituted residues are shown in red.

Phase separation (PS) or condensation, which is driven by multivalent interactions between macromolecules, has been suggested as one possible pathway towards TDP-43 aggregation.^[15, 16]^ While it can be mediated by intrinsically disordered regions (IDRs), which form multiple weak, transient contacts, RNA molecules and folded protein domains can also act as essential scaffolds and drivers of the process.^[16, 17]^ Condensates formed from recombinant protein *in vitro* often undergo a liquid-to-solid transition, whereby initially liquid-like droplets are converted into insoluble, fibrillar aggregates.^[18, 19]^ It has been proposed that phase separation could be a pathway to the formation of cytoplasmic inclusions observed in patients with ALS, FTD, and AD. However, the phase separation-to-fibril conversion pathway has not been definitively proven, and another possible route to amyloid fibril formation is direct misfolding into β-sheets, followed by “stapling” of misfolded monomers into an amyloid fibril ^[20, 21]^, independent of PS. In fact, several recent studies suggest that PS may even exert a protective effect against pathological amyloid formation.^[22, 23]^ Beyond phase separation, TDP-43 has recently been shown to adopt an additional mode of self-organisation. Shinn et al. demonstrated that full-length TDP-43 exhibits a block-copolymer-like architecture that encodes specific patterns of inter-domain homotypic and heterotypic attractions and repulsions among its domains, thereby driving microphase separation and the formation of ordered, size-limited assemblies of 12 molecules, yielding a structure ∼33 nm in diameter.[24]

One of the key, but understudied regulators is post-translational phosphorylation of TDP-43, which is detected in ALS/FTD patients ^[25, 26]^, but not in healthy cells and hence has long been considered a driver of TDP-43 aggregation. Notably, in disease-associated TDP-43 inclusions, phosphosites are highly enriched in the LCD: mass spectrometry analyses have revealed that 23 of the pathology-related phosphosites are localized in the C-terminal LCD, including the diagnostically significant Ser409/410 - a marker routinely used in post-mortem neuropathological diagnosis ^[25, 27]^ whereas only 5 disease-associated phospho-sites are found in the RRM domains and 2 in the NLS.[27] The addition of negatively charged phosphate groups to Ser/Thr residues within the LCD alters the sequence’s charge patterning. TDP-43 LCD is 148 amino acids long (aa: 267-414). Roughly half of these residues are hydrophobic (aliphatics, aromatics and glycine), the remaining being mostly polar, as well as 9 charged ones (6 cationic and 3 anionic) and 4 prolines (**Figure 1**). Phase separation of this domain is driven mainly by its aromatic and aliphatic residues[28], with an additional contribution of cooperative interchain interactions between residues in the conserved helical region (CR; aa 319-341).^[3, 29]^ Mass spectrometry analyses have revealed that phosphorylation sites are highly enriched in the C-terminal third of the LCD, beyond the conserved helical region: 13 of 13 serines in this region are phosphorylated in disease.^[27, 30, 31]^ Phosphorylation effectively converts the LCD into a block copolymer-like architecture consisting of a relatively hydrophobic N-terminal and a negatively charged, phosphorylated C-terminal “block.” Charged residues can exert a nuanced influence on PS: high net charge generally suppresses PS due to electrostatic repulsion, whereas segregation into blocks of same-sign residues can, in contrast, enhance it.^[32, 33, 34]^ Electrostatic interactions are extremely sensitive to the ionic strength of the solvent and pH.

Gruijs da Silva et al previously used a 12 C-terminal phosphomimetic serine-to-aspartate (S→D) substitution to make TDP-43 full-length condensates more liquid-like and dynamic, antagonize its aggregation and enhance its solubility in neurons.[35] In the present work, we aimed to investigate the effect of these 12 phosphorylations. As all of these 12(S→D) mutations are placed within the TDP43-LCD we focused on revealing the conformational ensemble and self-organization of TDP-43 LCD. Combining coarse-grained simulations with dynamic light scattering (DLS), negative-stain transmission electron microscopy (TEM), cryo-electron microscopy (cryo-EM), turbidity assays, fluorescence imaging, and fluorescence-based critical micelle concentration (CMC) measurements, we demonstrated that the phosphorylated form of TDP-43 LCD, and phosphomimetic variants, forms stable spherical micelles above the low-micromolar CMC. The hydrodynamic diameter of these micelles remains constant across a wide range of concentrations, consistent with classical core-shell micellar behavior. Phosphomimetic mutants (12D and 12DD LCD) form nearly identical assemblies, suggesting they capture the phosphorylation-induced charge redistribution (sequences of all protein constructs are shown in **Table S1**). In contrast, the unphosphorylated TDP-43 LCD (non-phLCD) forms dynamic liquid condensates that progressively mature into fibrils under comparable conditions. At high protein concentrations and elevated ionic strength, the 12D and 12DD variants - but not the phosphorylated LCD - were also found to form worm-like micelles. Both negative-stain and cryo-electron microscopy confirm the spherical architecture of these finite nanostructures. Collectively, these findings identify phosphorylation as a conformational switch that redirects TDP-43 LCD assembly to tune macrophase separation and potential fibril formation to micellization, revealing a previously unrecognized regulatory mechanism with potential therapeutic implications for TDP-43 proteinopathies.

## 2. Results

### 2.1 Phosphorylated TDP-43 LCD adopts a block-copolymer architecture that promotes micellization in coarse-grained simulations

To evaluate whether phosphorylation of the TDP-43 LCD could drive micellization, we first performed coarse-grained molecular simulations. Implicit-solvent coarse-grained models enable efficient simulation of condensates containing hundreds of protein chains by averaging-out solvent effects and representing each amino acid as a single interaction bead. They sacrifice atomic detail for computational performance, while nonetheless still capturing PS behaviour of IDRs.^[36, 37, 38, 39]^ We employed the original hydropathy scale (HPS) model, which is widely used for TDP-43 LCD and reproduces experimental trends qualitatively.^[4, 35, 39, 40, 41]^

Simulations were conducted for unmodified TDP-43 LCD, phosphorylated TDP-43 LCD-12pS (12 C-terminal phosphoserines), and the phosphomimetic variants 12D, as previously used by Gruijs da Silva et al.^[35]^ In addition, we replaced each phosphoserine with two aspartates to create another phosphomimic that we term 12DD, and is 12 amino acids longer than the TDP-43-LCD on its own. (**Figure 1c**, full sequences are shown in **Table S1).**

The pKa for the second deprotonation in phosphorylated serine (∼6.1) residues is very close to physiological pH, which causes an oscillation between −1 and −2 charge states.[42] Clustering of multiple same-charge residues has also been shown to shift the pKa up, bringing it even closer to pH 7. The experimental charge patterning of phosphorylated TDP-43 LCD thus likely lies somewhere between the two mutants. Sliding-window profiles of the hydrophobicity parameter (λ) and net charge per residue reflect differences between phosphorylated and unphosphorylated sequences (**Figure 2a**). The unmodified LCD contains few charged residues and remains hydrophobic along most of its length. In contrast, TDP-43-LCD-12pS displays a pronounced accumulation of negative charge in the C-terminal region, leading to a sharp local decrease in effective hydrophobicity. The 12D and the 12DD phosphomimetic variants recapitulate this redistribution of charge and hydropathy, consistent with the emergence of a block-copolymer-like architecture upon phosphorylation.

**Figure 2.**
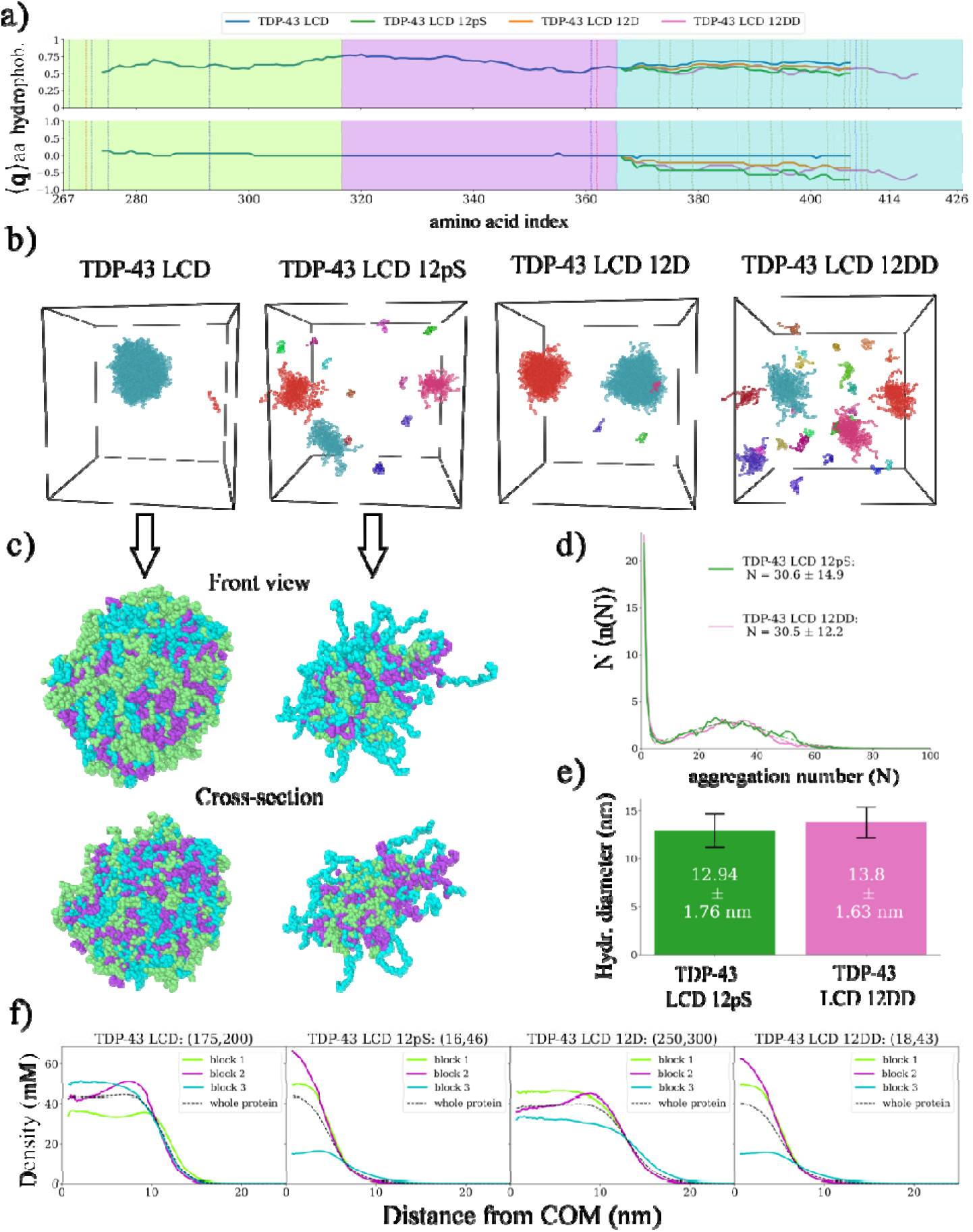
Coarse-grained simulations predict phosphorylation-induced micellization in TDP-43 LCD. **a)** Sliding average of mean hydrophobicity (top) and net charge per residue (bottom) along the sequence. The sequence is divided into 3 blocks of nearly identical length, shown in cyan, pink, and light green, representing the C-terminal, central, and N-terminal blocks. Dashed lines indicate sequence charges, blue and red represent positive and negative charges, respectively, and olive lines represent phosphorylation sites. **b)** Snapshots of simulation trajectories. Different colours represent different clusters. A zoomed-in snapshot of a TDP-43 LCD 12pS micelle is shown next to them. A front view and a cross-sectioned (sliced in half) view are provided. Amino acids are coloured according to the colour-coding in a and f. They provide a visualisation of the segregated internal architecture of the micelles. **c)** Zoomed-in snapshots of TDP-43 LCD and TDP-43 LCD 12pS assemblies. A frontal view and a cross-sectioned (sliced in half) view are provided. Amino acids in different sequence blocks are coloured according to the colour-coding in a and f. **d)** Size distribution (full lines) and corresponding fitting curves (dashed lines) for the 2 sequences (TDP-43 LCD 12pS and 12DD) forming finite-sized structures. Mean aggregation number and standard deviation are shown in the legend. **e)** Mean hydrodynamic diameter for both micellizing sequences. Results were obtained as a weighted average of the hydrodynamic radii of clusters of sizes within one standard deviation of the mean aggregation number of the micelles, as determined from the fit to the size distributions. Error bars represent the standard deviation of the distribution of hydrodynamic diameters. **f)** Monomer density profiles per block, measured as a function of the distance from the micelle’s (or cluster’s) centre of mass. Numbers in parentheses indicate the aggregation-number interval used to compute the density profiles. Images in **b** made with OVITO.[47]

Traditional slab simulations with a fixed number of chains are well suited to study macroscopic phase separation but may be less appropriate for detecting finite-sized micelle-like assemblies. The periodic boundary conditions in the x and y directions may cause micelles to interact with their own periodic images. A cubic box is therefore necessary in order to study such structures. The transition from a slab (1D) to a 3D simulation box reduces protein-protein contact probabilities. This effect is further amplified at the very low concentrations of TDP-43 LCD, where the likelihood of intermolecular encounters is intrinsically small, leading to significantly longer equilibration times. To overcome this limitation, we employed an in-house semi-grand canonical Monte Carlo (SGCMC) algorithm, which introduces a reservoir of noninteracting “virtual” chains that can be turned into interacting “real” chains, thereby enabling chain insertion and deletion moves between molecular dynamics (MD) steps. This approach allows the system to reach a steady state at very dilute concentrations and slow diffusion times. In standard canonical simulations, micelle size is sensitive to the overall number of molecules in the simulation box. Our SGCMC keeps the concentration of real chains constant in the dilute regions of the simulation box over long times, thereby minimising finite-size effects that may affect micelle size. Each simulation box contained 200 chains (considering both virtual and real), though the number of real chains fluctuates due to the SGCMC approach.

Under these conditions, unphosphorylated TDP-43-LCD (non-phLCD) readily formed continuously growing clusters (**Figure 2b**), consistent with its strong experimental propensity for phase separation.^[3, 43]^ In contrast, both 12pS and 12DD variants assembled into multiple small, roughly spherical clusters containing ∼20-40 chains that remained stable over extended simulation times, which is indicative of micellization. These clusters exhibited a dense core and a diffuse corona formed by chain ends pointing toward the solvent. The 12D phosphomimetic behaved differently. Preliminary MD simulations (**Figure S1**) showed that most chains partitioned into a single large droplet with a narrow, diffuse interface. To test whether this reflected an increased preferred micelle size due to the reduced electrostatic repulsion, we increased the maximum number of chains to 500. Even at this scale, 12D formed large, ever-growing droplets (**Figure 2b**) that contained almost up to 350 chains, but did not reach values close to 500 because a second droplet was nucleated. This hints at a preference for macroscopic phase separation over micellization.

Cluster size distributions (**Figure 2d**) for TDP-43 LCD-12pS and 12DD display distributions characteristic of micellization: a large monomer population, a rapidly decaying oligomer regime, and a pronounced Gaussian-like peak at intermediate cluster sizes.[44] The distributions were fitted with the equation: *f*(*N*) = *a e^-N/b^ +c e-(N-N_0_)^2^/2 σ^2^*, where f(x) denotes the probability density for a chain to be located in a cluster of aggregation number N. The variable N refers to the number of protein chains in the micelle and is usually referred to as the micelle’s aggregation number. The second (Gaussian) term yielded a mean aggregation number of 30.6 ± 14.9 chains for 12pS and 30.5 ± 12.2 chains for 12DD (results are given as *N*_0_ ± *σ*). We define these as the size intervals within which clusters are considered micelles in subsequent analyses of our simulations.

Micellization arises when attractive interactions within the hydrophobic block balance the repulsive electrostatic and solvation forces from the residues in the hydrophilic block. Growth beyond the preferred aggregation number leads to a tighter packing between amino acids in the corona, which comes with a high electrostatic interfacial penalty to the free energy.[44] Consistent with this mechanism, shape analysis revealed that larger micelles exhibit increased asphericity and shape anisotropy parameter (**Figure S2**), indicating deformation away from optimal spherical geometry, while acylindricity slightly decreased, reflecting a tendency toward unstable elongated elliptical shapes in oversized assemblies. Sequences prone to phase separation, on the other hand, show a monotonic decrease in asphericity, acylindricity and shape anisotropy for increasing aggregation number.

Fixed-size micelles possess a characteristic internal organisation, with hydrophobic residues typically forming a compact core and hydrophilic residues exposed to solvent. To quantify this architecture, we divided the LCD into three equal-length blocks (49 to 50 residues each), with all phosphorylation sites located in block 3 (residues 101-148 of the LCD and 367-414 of full-length TDP-43), while blocks 1 and 2 retain their compositions - and therefore hydrophobicity and net charge - in all 4 sequences (**Figure 2a**). The corresponding residue indices are detailed in Table 1 in Section 4.13.3.4 of the Methods. Per-block density profiles for the most probable cluster sizes (**Figure 2f**) reveal clear structural differences. In unphosphorylated LCD droplets, residues are distributed relatively uniformly, with slight enrichment of chain termini at the interface, as has been reported for small droplets of hydrophobic polymers.[45] In contrast, both 12pS and 12DD micelles exhibit pronounced asymmetry: the core density of blocks 1 and 2 is three to fourfold higher than that of block 3, which is preferentially localised at the surface.

**Table 1.** Intervals defining the 3 blocks of TDP-43 LCD. Amino acid indexes are given relative to the first residue of the LCD (left) and to the first residue of full-length TDP-43.

| Sequence \ Block | 1 | 2 | 3 |
| --- | --- | --- | --- |
| TDP-43 LCD | 1-50 / 267-316 | 51-100 / 317-366 | 101-148 / 367-414 |
| TDP-43 LCD 12pS | 1-50 / 267-316 | 51-100 / 317-366 | 101-148 / 367-414 |
| TDP-43 LCD 12D | 1-50 / 267-316 | 51-100 / 317-366 | 101-148 / 367-414 |
| TDP-43 LCD 12DD | 1-50 / 267-316 | 51-100 / 317-366 | 101-160 / 367-426 |

To further characterise the molecular organisation within the assemblies, we analysed interchain contact maps (**Figure S3**). In all sequences, contacts between residues in block 2 (2 −2) were the most frequent, consistent with its pronounced hydrophobicity. In unphosphorylated TDP-43 LCD, contacts are broadly distributed along the sequence, with 2-2 interactions dominating, followed by 2-3, while 1-1 contacts are less frequent, likely due to the modest net positive charge of block 1. In contrast, 3-3 contacts are nearly absent in phosphorylated LCD, as well as in the 12D and 12DD phosphomimetics, reflecting strong repulsive interactions among negatively charged residues in block 3. Contacts between blocks 2-2 and 2-3 also decrease, though to a lesser extent, while 2-1 contacts are minimally affected.

Notably, 1-1 contacts remain essentially unchanged and, although least frequent in the unphosphorylated sequence, become among the top three contact types in LCD-12pS and LCD-12DD, suggesting that block 1 contributes significantly to the formation of the micelle’s hydrophobic core. Finally, a localised increase in contacts between negatively charged residues in block 3 and the few positive residues near the N-terminus of block 1 is observed. Collectively, these contact maps highlight how phosphorylation reshapes the interaction network of TDP-43 LCD, promoting segregation of the charged C-terminal block and stabilising the hydrophobic N-terminal core of micelles.

Phosphoserines do not always exhibit charge −2.0e. The pKa for the second deprotonation is close to the physiological pH, giving rise to a non-negligible population of phosphoserines with charge −1e. To account for that, we conducted a set of simulations for TDP-43 LCD 12pS in which we assigned all phosphoserines a charge of −1.5e (**Figure S4**). Our goal was to approximate a slightly less negatively charged corona. Micellization still occurred; however, the mean aggregation number was higher and the dispersion was significantly larger (**Figure S4c**). This led to a large mean hydrodynamic diameter (**Figure S4d**). The computed per-block density profiles (**Figure S4e**) and interchain contact maps (**Figure S4f**) hint at a less “blocky” chain, with decreased segregation of blocks inside the micelle and more smoothly distributed contact frequencies for the 12pS variant with charge - 2.0e. The sequence is still more “blocky” than the 12D mutant. Analysis of the asphericity and shape anisotropy (**Figure S4g**) shine a light at the broadening of the aggregation number histogram. The decreased electrostatic repulsion in the corona significantly decreases the penalty for tighter packing of negatively charged amino acids in the corona, reflected by an almost fully flattened U-shaped profile of the asphericity, which allows the micelle to explore a wider array of aggregation numbers. We also observed the formation of short-lived oblong-shaped micelles, which could be precursors to worm-like chains (**Figure S4b**), a phenomenon not observed when phosphoserines had charge −2e and also contributed to the broadening of the aggregation number histogram.

We also used simulations to test whether the differing chain lengths between 12D/12pS and 12DD play a significant role. In order to test that we designed two new mutants, one containing 12 DA substitutions (named 12DA) and one containing 12 DD substitutions that is 148 amino acids long (named 12DD-148aa), created by deleting some residues around the DD substitutions. The 12DA sequence is a longer analog of the 12D sequence, whereas the 12DD-148aa serves as a model for the 12DD mutant but with identical length to 12D and 12pS. As we show in **Figures S5** and **S6**, the 12DA construct behaves very similarly to 12D, with both exhibiting phase separation (**Figure S5a**) and presenting nearly identical hydrodynamic diameters (**Figure S5c**), per-block density profiles (**Figure S5d**), shape descriptors (**Figure S6a**) and interchain contact maps (**Figure S6c**). We also observe identical phenotypes for 12DD and 12DD-148aa, with 12DD-148 presenting as slightly larger micelles (**Figure S5b and S5c**) and nearly identical per-block density profiles (**Figure S5e**), shape descriptors (**Figure S6b**) and interchain contact maps (**Figure S6c**) to those of TDP-43 LCD 12pS and 12DD. These results suggest that the 12-residue increase in protein length for the 12DD mutant does not play a significant role in determining supramolecular assembly.

Overall, the simulations strongly support micellization for TDP-43 LCD-12pS and TDP-43 LCD-12DD, consistent with a phosphorylation-induced block-copolymer architecture. The 12D variant instead seems to favor macroscopic PS; while larger preferred micelle sizes cannot be excluded, distinguishing them from phase-separated droplets would require substantially larger-scale simulations. It is important to highlight that, although the simulation methods employed here serve as an important prediction tool, they are intrinsically limited by the high amount of coarse-graining of the force field. Different parameterizations have been shown to over and underestimate the effective “stickiness” of certain amino acids and they completely ignore secondary structure - although alpha-fold[46] predicts preservation of disorder for the phosphomimetics as shown in **Figure S7** - as well as steric effects and residue ionization. They nonetheless provide significant insight into the phenomena underlying supramolecular assembly of phosphorylated and unphosphorylated TDP-43 LCD and the associated phosphomimetics.

### 2.2. Phosphorylation redirects TDP-43 LCD self-assembly from reversible PS to stable size-limited spherical micelle formation

To validate predictions from coarse-grained simulations, we compared the phase behaviour of unphosphorylated (non-phLCD) and *in vitro* phosphorylated TDP-43 LCD (phLCD). The phosphorylated LCD was generated following published protocols by incubating the purified LCD domain with casein kinase 1δ (CK1δ) under optimized conditions that achieve robust phosphorylation of key C-terminal sites, as confirmed by Western Blotting (see Methods Section 4.3 and **Figure S8**).^[25, 35]^ To assess the stoichiometry of phosphorylation, i.e. the number of phosphorylated serine residues, we used native mass spectrometry[48](see Methods Section 4.8). The acquired mass spectrum (**Figure S9**) revealed several charge state distributions corresponding to differently phosphorylated TDP-43 LCD populations, all consistent with a monomeric protein. Close inspection of the peak reveals a range of seven up to thirteen phosphorylation sites per population. Importantly, the highest intensities were observed for populations with 10, 11, or 12 phosphorylation sites (**Figure S9b**), consistent with extensive phosphorylation of the 12 C-terminal serine residues identified by Gruijs da Silva et al.[35] as the canonical CK1δ phosphorylation pattern in TDP-43 LCD. We adopted this 12-site pattern both for our coarse-grained simulations (Section 2.1) and for the design of the experimental phosphomimetic constructs characterised in the following section, focusing on the two mutants - 12D and 12DD - that bracket the effective C-terminal charge expected for the fully phosphorylated state (phosphoserines carry between −1e and −2e at physiological pH; **see Section 2.3** for detailed characterisation).

In physiological buffer (here defined as 150 mM NaCl, 50 mM HEPES, 10 mM MgCl_2_, 1mM DTT, pH 7.5, 25 °C), at low-micromolar concentrations (1 - 15 μM), the unmodified LCD assembled into large, polydisperse species with hydrodynamic diameters predominantly exceeding 100 nm, consistent with macrophase separation (**Figure 3a**). These droplets exhibited classical PS hallmarks: their intensity-weighted hydrodynamic diameter increased continuously with rising protein concentration, reflecting droplet growth through coalescence and Ostwald ripening, followed by progressive maturation into more viscous or gel-like states - a process implicated in the transition toward pathological aggregates.[4]

**Figure 3.**
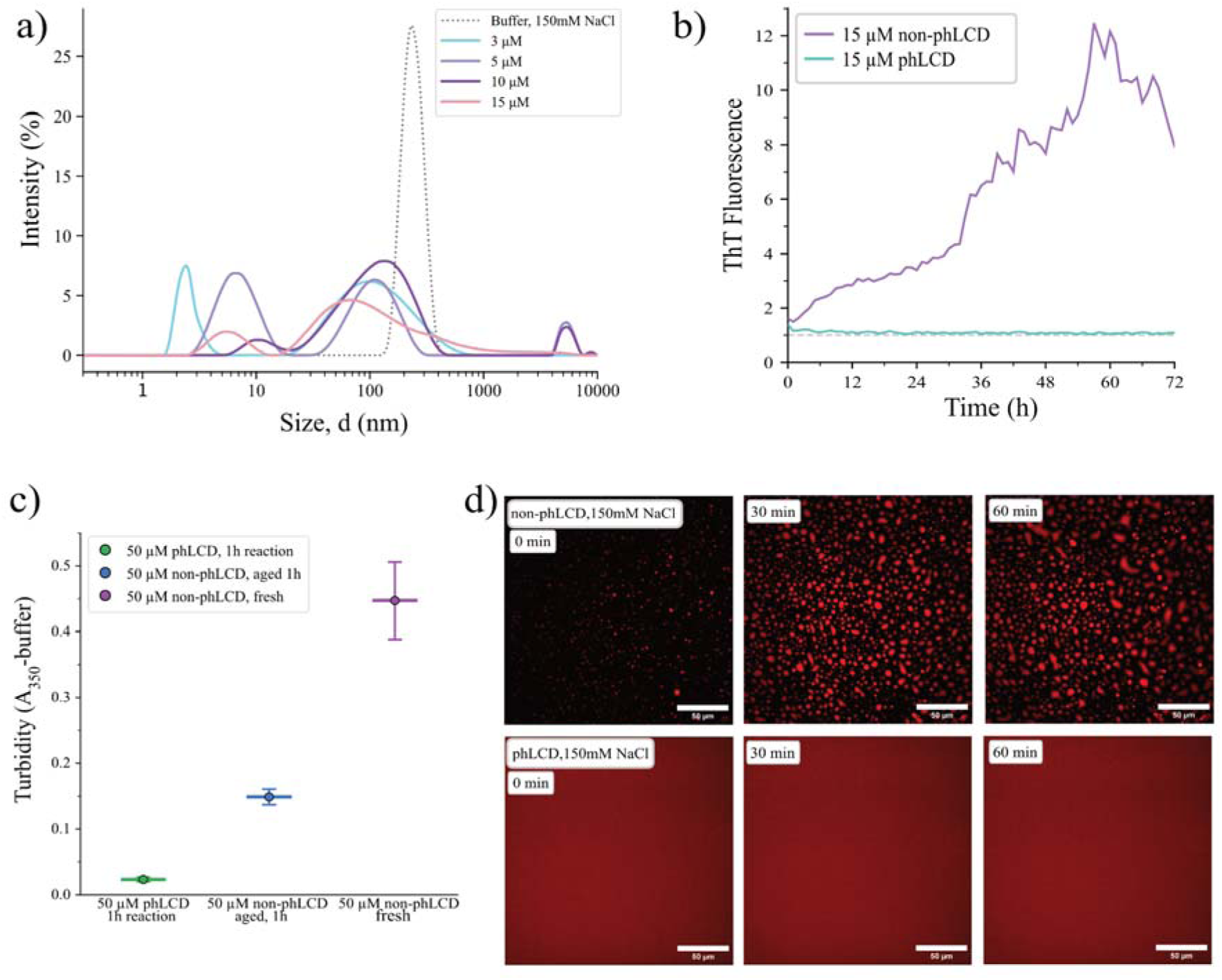
Phosphorylation suppresses macroscopic phase separation and amyloid-like aggregation of TDP-43 LCD. **a)** Intensity-weighted dynamic light scattering (DLS) size distributions (diameter in nm) of unphosphorylated TDP-43 LCD at indicated concentrations (3-15 μM) and buffer control (150 mM NaCl, 50 mM HEPES, 10 mM MgCl_2_, 1 mM DTT, pH 7.5, 25 °C). **b)** Thioflavin-T (ThT) fluorescence kinetics (fold change) over 72 h for 15 μM unphosphorylated LCD (non-phLCD) versus 15 μM phosphorylated LCD (phLCD). **c)** Turbidity (A_350nm_- buffer) of three samples at 50 μM total protein in Pluronic-passivated wells: phLCD after 1 h in-well reaction, “aged” non-phLCD (1 h), and “fresh” non-phLCD (mean ± s.d., n=3). **d)** Time-lapse spinning-disk confocal fluorescence microscopy of LD655-doped samples (10 nM LD655-A366C TDP-43 LCD, ∼1:5,000 molar ratio) at 50 μM total protein, in Pluronic-passivated glass bottom chambers at 0, 30 and 60 min. Upper panels: non-phLCD shows abundant droplets that progressively coalesce; lower panels: phLCD remains droplet-free throughout. Scale bars, 50 μm.

Consistent with this maturation, a Thioflavin-T (ThT) assay performed at 15 μM showed a progressive increase in fluorescence for unphosphorylated LCD over approximately 60 h, followed by a decline, likely reflecting sedimentation of mature fibrillar aggregates in the plate (**Figure 3b**). In contrast, the fluorescence signal for phosphorylated LCD remained at baseline throughout the time course, indicating the absence of amyloid-like assemblies.

To independently verify the contrasting phase behavior of unphosphorylated and phosphorylated LCD by a bulk scattering measurement, we performed a turbidity assay (**Figure 3c**). Three samples were compared in parallel under matched buffer conditions at 50 µM total protein and 150 mM NaCl, loaded into wells passivated with the detergent Pluronic), of a 384-well plate to suppress surface-induced nucleation (see Methods Section 4.6): (i) CK1δ-phosphorylated LCD (phLCD), incubated in the well for ∼1 h to allow the kinase reaction to proceed; (ii) “aged” unphosphorylated LCD (non-phLCD, prepared in parallel with phLCD without kinase and ATP, and incubated in the well for the same ∼1 h duration); and (iii) “fresh” non-phLCD, freshly diluted and measured immediately. Absorbance at 350 nm - a direct readout of light scattering by phase-separated droplets - was recorded simultaneously for all three samples. The phosphorylated sample showed no detectable turbidity above buffer baseline (A_350nm_ ≈ 0.02), whereas both unphosphorylated samples produced strong absorbance signals. The fresh non-phLCD sample displayed the highest turbidity (A_350nm_ ≈ 0.45), reflecting the abundant population of small-to-medium-sized droplets that form within minutes of dilution and efficiently scatter light. The aged non-phLCD sample, in contrast, showed reduced - but still substantial - turbidity (A_350nm_ ≈ 0.15), consistent with progressive coalescence into fewer, larger droplets and partial sedimentation of mature condensates over the 1h incubation window. This kinetic decrease in turbidity for the unphosphorylated LCD mirrors the DLS observation of continuously growing hydrodynamic diameters and is consistent with classical hallmarks of liquid-liquid phase separation followed by maturation. Crucially, under identical buffer composition and well passivation, phLCD shows no turbidity at either time point, demonstrating that the absence of phase separation in phLCD reflects a fundamental change in the assembly pathway rather than a kinetic delay.

To directly visualize this contrasting phase behavior, we performed time-lapse spinning-disk confocal fluorescence microscopy on samples doped with a small fraction of fluorescently LD655-labeled A366C TDP-43 LCD (10 nM, ∼1:5,000 molar ratio relative to unlabeled LCD; see Methods Section 4.7). Imaging was performed in Pluronic-passivated wells of a 15 well 3D glass bottom dish to suppress surface-induced nucleation, ensuring that any observed droplets reflect bulk-solution phase behavior. At 150 mM NaCl and 50 µM total protein, the unphosphorylated LCD rapidly assembled into a dense population of micron-scale fluorescent spherical droplets, already abundant within a few minutes of sample loading (**Figure 3d**). Over the 60min observation window the droplet population evolved progressively, with smaller droplets coalescing into fewer, larger droplets - directly visualizing the maturation kinetics inferred from the bulk turbidity and DLS measurements (**Movie S1**). Equivalent imaging of the unphosphorylated LCD at elevated ionic strength (300 and 500 mM NaCl; **Movies S2, S3**) revealed qualitatively the same phase-separation and coalescence behavior at all tested salt concentrations, confirming that the unphosphorylated LCD undergoes robust liquid-liquid phase separation followed by droplet maturation across the entire salt range tested in this study. In stark contrast, equivalent imaging of CK1δ-phosphorylated LCD at 150 mM NaCl - initiated after a 1h in-well incubation that allows the phosphorylation reaction to proceed to near-completion - yielded no detectable droplets throughout the entire 60 min observation window (**Figure 3d, Movie S4**), in direct agreement with the ThT and turbidity results. Together, these direct optical observations confirm that the suppression of phase separation in the phosphorylated form is robust and is not an artefact of either bulk-measurement geometry or ionic strength.

In sharp contrast to unphosphorylated LCD, which did not produce any population in the 10-100 nm range, phosphorylated LCD formed a distinct population of discrete, spherical nanoparticles (**Figure 4a,b**). Intensity-weighted DLS in backscattering geometry (173°) across a wide concentration range (0.05 - 50 μM) revealed a stable particle population centred around 30 nm that was absent in both control conditions: buffer alone (no protein, no kinase, no ATP) and buffer supplemented with kinase and ATP but lacking protein (**Figure 4a**). Large species exceeding 100 nm, detected in both unphosphorylated and phosphorylated TDP-43 LCD samples as well as in buffer-only controls, were attributed to background scattering rather than specific protein assemblies. The size of this population remained constant regardless of concentration and incubation time (**Figure S10**) and was therefore excluded from further analysis. This concentration-independent behavior strongly suggested the formation of finite-size supramolecular assemblies rather than growing phase-separated droplets. A direct side-by-side comparison of the distributions at selected concentrations highlights the dramatic difference: unphosphorylated LCD formed large polydisperse species, while phosphorylated LCD exhibited a stable population of ∼ 30 nm diameter size.

**Figure 4.**
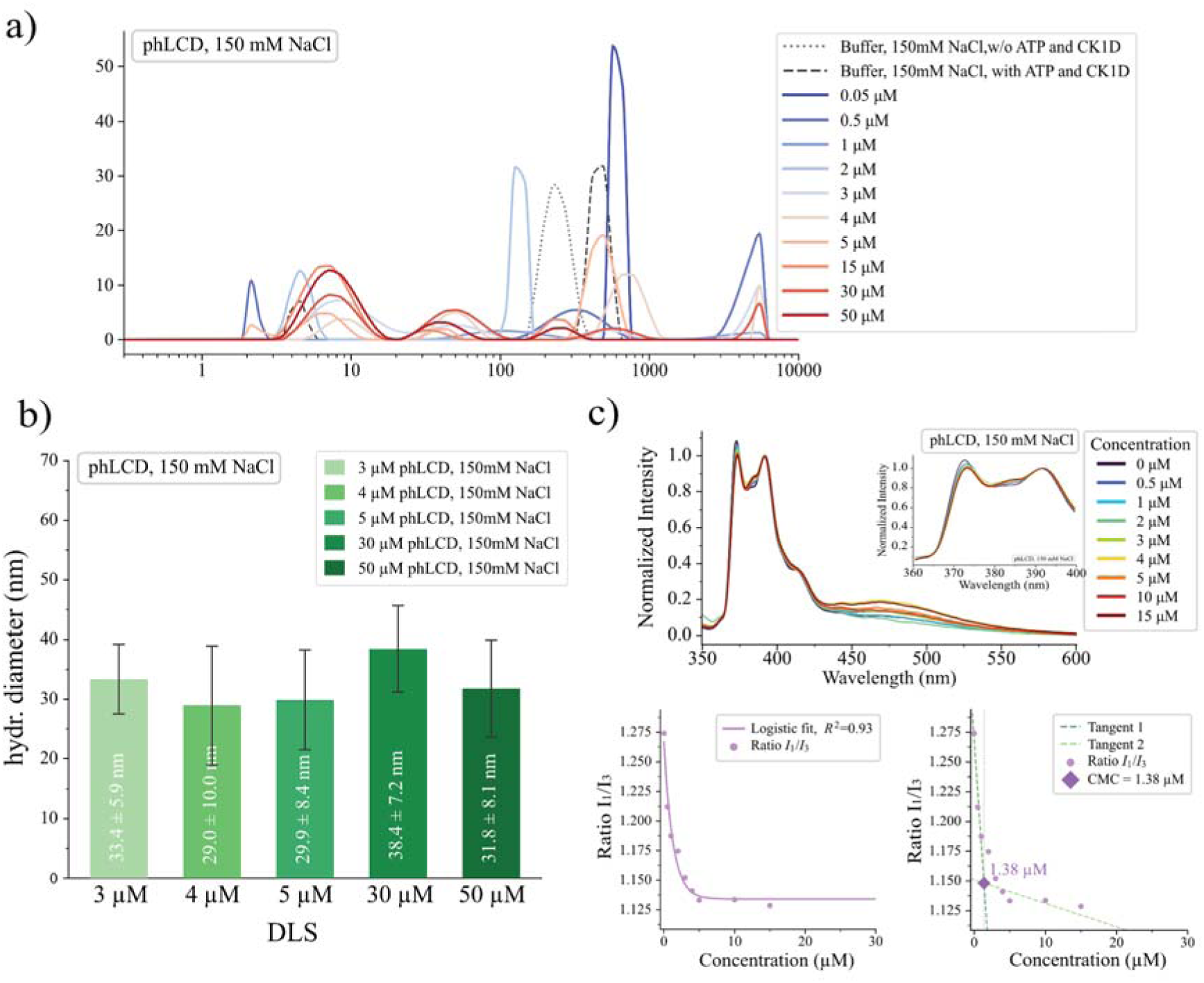
Phosphorylated TDP-43 LCD forms discrete, concentration-independent spherical assemblies with a low-micromolar critical micelle concentration. **a)** Intensity-weighted hydrodynamic diameter distributions of phLCD at concentrations ranging from 0.05 to 50 µM, together with two buffer controls: buffer alone (no protein, no kinase, no ATP) and buffer supplemented with CK1δ and ATP but lacking protein. A stable population centred around 30 nm is observed across the entire concentration range and is absent from both controls. **b)** Mean hydrodynamic diameters extracted from the protein-assembly peak of the DLS distributions in panel a for selected concentrations (3, 4, 5, 30 and 50 μM phLCD; mean ± s.d., n =12). **c)** Determination of critical micelle concentration using pyrene. Upper panel: normalized fluorescence emission spectra at increasing phLCD concentrations. Lower panel: Ratio of pyrene vibronic bands (I_1_/I_3_) plotted against phLCD concentration, fitted with a logistic function (R^2^ = 0.93) and tangent lines indicating CMC ≈ 1.38 μM.

The CMC was determined using the environment-sensitive fluorescent probe pyrene (**Figure 4c**).^[49, 50, 51, 52]^ Pyrene reports on local polarity through the ratio of its first to third vibronic emission bands (I_1_/I_3_): in pure water, the theoretical baseline is ∼1.87, but under physiological ionic strength (150 mM NaCl) it is compressed to ∼1.25, as salt lowers the aqueous solubility of pyrene and promotes weak self-association of the probe even in the absence of protein. Upon incorporation into a hydrophobic micellar core, I_1_/I_3_ decreases toward a lower plateau. For phosphorylated TDP-43 LCD at 150 mM NaCl, a progressive decrease from ∼1.27 (buffer baseline) to a plateau of 1.13 ± 0.002 above 5 µM was observed (Δ I_1_/I_3_ = 0.14; **Figure 4c**). The transition is gradual rather than sharp - characteristic of protein-based amphiphiles[44], where the intrinsic disorder of the LCD leads to partial hydrophobic exposure in monomeric and oligomeric species below the CMC, producing a continuous rather than cooperative I_1_/I_3_ decrease. The CMC was therefore estimated by the tangent-intersection method (linear fits to the descending and plateau regions; see Methods 4.12.1), yielding ∼1.4 µM (**Table S2**). This range is fully consistent with the independent onset of stable nanoparticle formation, as observed by DLS and TEM, between 1 and 3 µM. The convergence of three orthogonal methods brackets the CMC in the low-micromolar range and confirms micellar rather than continuous condensation behavior.

Negative-stain transmission electron microscopy (TEM) independently confirmed both the spherical morphology and the size of the assemblies. At a concentration of 5 µM, which is above the CMC established by pyrene and DLS, the phosphorylated TDP-43 sample appeared as a population of spherical nanoparticles (**Figure 5a**). Quantitative analysis of particle diameters from >200 particles yielded 27.2 ± 6.0 nm, in excellent agreement with the DLS value of 29.9 ± 8.4 nm (n = 12 biological replicates; **Figure 5b**). The slightly smaller diameter obtained by TEM relative to DLS is consistent with dehydration-induced deformation during negative-stain sample preparation, which can lead to partial collapse or shrinkage of soft protein assemblies, resulting in an underestimated projected diameter compared to the hydrated solution-state size. The observed size dispersity (coefficient of variation (CV) = 22% by TEM) reflects both the intrinsic distribution of aggregation numbers which is thermodynamically expected for protein-based amphiphiles in dynamic monomer-micelle equilibrium[44], and the variable degree of particle deformation during specimen preparation. Although CK1δ kinase is a processive enzyme that quantitatively phosphorylates TDP-43 LCDs at its preferred serine sites, some heterogeneity in charge patterning due to enzymatic processing is to be expected. Individual micelles will thus be made up of slightly different chain populations, which increases size heterogeneity. This polydispersity is further consistent with the broad aggregation number distributions (**Figure 2d**) and large standard deviation in hydrodynamic diameter estimates (**Figure 2e**) obtained from coarse-grained simulations (Section 2.1).

**Figure 5.**
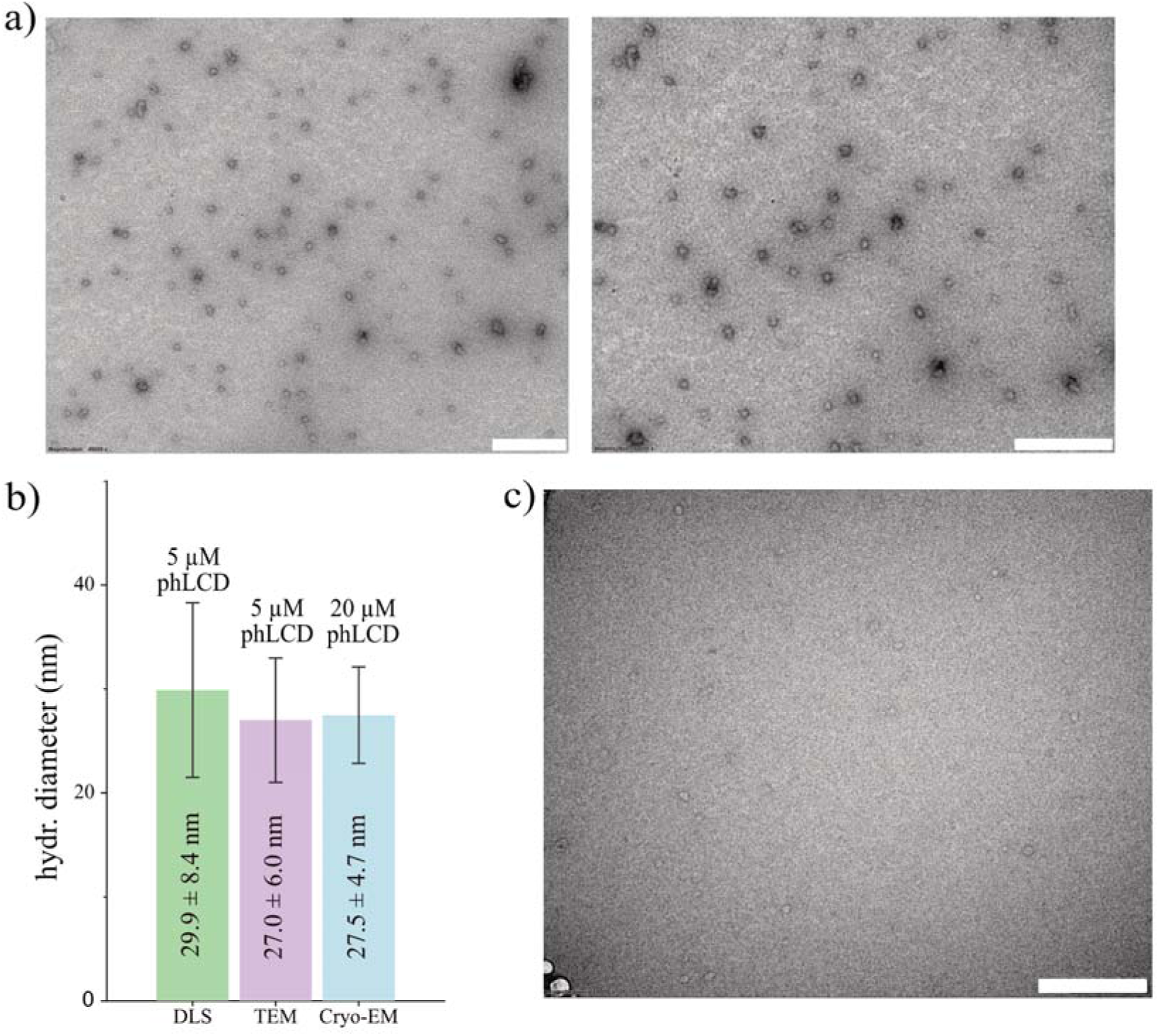
Direct visualization of phosphorylated TDP-43 LCD micelles by negative-stain TEM and cryo-EM confirms a uniform spherical architecture with native diameters of ∼30 nm. **a)** Representative negative-stain TEM images of 5 μM phLCD in 150 mM NaCl conditions, showing spherical nanoparticles. Scale bar, 300 nm. **b)** Direct comparison of mean micelle diameters obtained by three orthogonal methods: DLS (peak maximum of the intensity-weighted hydrodynamic diameters distribution; 5 μM phLCD, n = 12 biological replicates), negative-stain TEM (5 μM phLCD, >200 particles analyzed), and cryo-EM in vitreous ice (20 μM phLCD, n >100 particles). Bars indicate mean ± s.d. **c)** Representative cryo-EM micrograph of phLCD at 20 μM in vitreous ice, showing well-dispersed spherical particles consistent with the negative-stain TEM and DLS measurements. Scale bar, 300 nm.

To further confirm the native hydrated morphology of phLCD assemblies and to rule out drying-induced artefacts as a source of any size variation between methods, we performed cryo-electron microscopy (cryo-EM) at 20 µM phLCD in vitreous ice at 150 mM NaCl (**Figure 5c**) (see Methods Section 4.11). Cryo-EM micrographs revealed monodisperse spherical particles with a mean diameter of 27.5 ± 4.7 nm, in excellent agreement with both negative-stain TEM (27.2 ± 6.0 nm) and DLS (29.9 ± 8.4 nm) (**Figure 5b**). The cross-validation of phLCD micelle size by three independent techniques - DLS in solution, negative-stain TEM under dehydration, and cryo-EM under native hydrated conditions - establishes the spherical micellar morphology of phLCD as a robust, drying-artifact-free observation. Importantly, no fibrillar structures were observed in the cryo-EM micrographs of phLCD at this elevated concentration, consistent with the absence of a ThT signal (**Figure 3b**) and supporting the interpretation that phosphorylation effectively suppresses the amyloid pathway.

Collectively, these biophysical and imaging approaches demonstrate that phosphorylation of the TDP-43 LCD fully redirects its self-assembly pathway: from macro PS and amyloid-like aggregation in unphosphorylated LCD to micro PS and formation of stable, size-limited spherical micelles whose dimensions are independent of concentration and time.

### 2.3. Phosphomimetic mutants TDP-43 LCD 12D and 12DD recapitulate the phosphorylated phenotype and form size-limited spherical micelles above CMC at physiological conditions

Analogously to the simulations above (Section 2.1), we next constructed the 12D LCD mutant (with single Ser→Asp substitutions at 12 sites) that mimics this C-terminal hyperphosphorylation which has been previously studied by Gruijs da Silva et al. 2022[35], and the 12DD LCD mutant (see **Figure 1c** and **Table S1** for protein sequences). These two constructs were chosen for experimental characterisation because they bracket the effective C-terminal charge expected for the fully phosphorylated state, given that phosphoserines carry between −1e and −2e at physiological pH, as was mentioned above. We note that the two additional simulation-only constructs introduced in Section 2.1 (12DA and 12DD-148aa) serve as in silico controls for chain-length effects (**Figures S5** and S**6**) and are not characterised experimentally, as the chain-length question they address is most directly resolved by comparing length-matched analogues in simulation. Native mass spectrometry confirmed the expected molecular masses of both phosphomimetic mutants (**Figure S9c, d**). To directly position phLCD on the same effective-charge axis as the two phosphomimetics, we ran all three constructs together with non-phLCD side-by-side on a single SDS-PAGE gel (**Figure S11**). Although SDS-PAGE primarily separates proteins by molecular weight, highly negatively charged and phosphorylated proteins are well known to migrate slower relative to neutral globular standards of the same mass. Consistent with this expectation, phLCD migrated as a single band located between the 12D and 12DD bands, providing an independent biochemical confirmation that the effective C-terminal charge of CK1δ-phosphorylated LCD lies between that of the 12D (-12e) and 12DD (-24e) constructs, in agreement with 10-12 phosphate groups whose protonation lies between −1e and −2e at physiological pH. Together, the overlapping native-MS charge envelopes and the intermediate SDS-PAGE mobility of phLCD support the use of 12D and 12DD as faithful charge-based mimics that bracket the phosphorylated state obtained by CK1δ treatment.

Both phosphomimetics in a physiological buffer (150 mM NaCl, 20 mM Tris, pH 7.5, 25 °C; the simpler Tris-based composition is used here because the phosphomimetic constructs do not require the kinase reaction mixture) reproduce the essential hallmarks of *in vitro* phosphorylated LCD micellization. Bar graphs of mean hydrodynamic diameters (**Figure 6a**) show the formation of stable nanoparticles of finite size above the threshold concentration: 48.5 ± 13.7 nm for 12D and 42.6 ± 19.8 nm for 12DD (mean ± s.d., n = 12 biological replicates, at 5 μM). The intensity-weighted DLS diameters of both phosphomimetics are noticeably larger than that of phLCD (29.9 ± 8.4 nm); the basis of this difference is addressed below in the context of the cross-method comparison. Like phLCD, the 12D micellar population was temporally stable: the size distribution remained essentially unchanged over a 5-day incubation at room temperature at 150 mM NaCl (**Figure S10b**), indicating that the spherical 12D assemblies represent a stable equilibrium state rather than a kinetically trapped intermediate.

**Figure 6.**
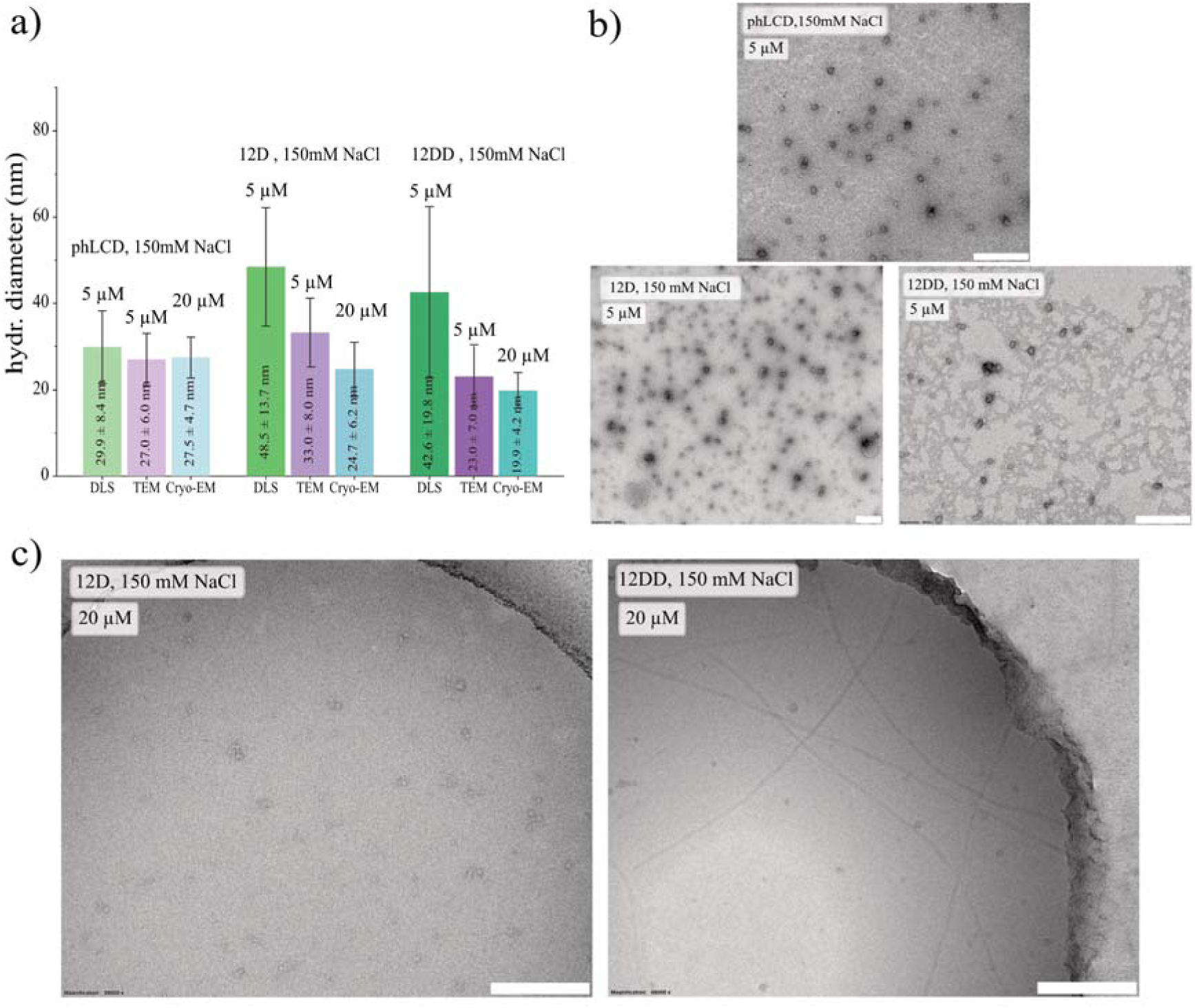
Phosphomimetic substitutions 12D and 12DD fully recapitulate the phenotype of phosphorylated TDP-43 LCD, forming stable spherical micelles of defined size above a low-micromolar critical micelle concentration. **a)** Direct comparison of mean diameters obtained by dynamic light scattering (DLS, 173° backscattering geometry, peak maximum of the intensity-weighted size distribution, n = 12 biological replicates), negative-stain TEM (>200 particles per construct) and cryo-EM in vitreous ice (>100 particles per construct) for phLCD, 12D and 12DD in physiological buffer (150 mM NaCl, pH 7.5, 25 °C). DLS and TEM measurements were performed at 5 µM; cryo-EM at 20 µM. Data are mean ± s.d. **b)** Representative negative-stain TEM images at 5 μM for phLCD (upper), 12D (lower left), and 12DD (lower right). Scale bars, 300 nm. **c)** Representative cryo-EM micrographs at 20 μM for 12D (left) and 12DD (right). The 12DD sample additionally shows occasional thin fibrillar structures coexisting with the dominant spherical micellar population - a minor subpopulation not detected by negative-stain TEM at 5 µM, indicating that, unlike phLCD, the doubly-charged 12DD phosphomimetic does not fully abolish access to fibrillar morphologies. Scale bars, 300 nm.

Negative-stain TEM images independently confirmed the spherical morphology of both variants at 5 µM (**Figure 6b**), with mean diameters measured over >200 particles of 33.0 ± 8.0 nm (CV = 24%) for 12D and 23.0 ± 7.0 nm (CV = 30%) for 12DD (**Figure 6a,b**). The size dispersity observed for both constructs is consistent with the broad aggregation number distributions expected for protein-based amphiphiles in dynamic monomer-micelle equilibrium[44], and is further supported by simulation data (Section 2.1).

To corroborate these measurements under natively hydrated conditions, we performed cryo-electron microscopy on both phosphomimetic constructs at 20 µM and 150 mM NaCl, following the same protocol used for phLCD in Section 2.2 (**Figure 6c**). Cryo-EM yielded mean diameters of 24.7 ± 6.2 nm for 12D (n >100 particles) and 19.9 ± 4.2 nm for 12DD (n > 100 particles), in close agreement with the corresponding negative-stain TEM measurements and substantially smaller than the intensity-weighted DLS values reported above. Because vitrification preserves the native hydrated state and avoids the dehydration inherent to negative staining, the close agreement between cryo-EM and TEM diameters (within ∼3 nm for all three constructs) excludes drying-induced shrinkage as a significant source of size underestimation by TEM and establishes a narrow native diameter range of ∼20-27 nm. The persistent gap between this microscopy-based range and the larger intensity-weighted DLS values - most pronounced for the phosphomimetic constructs - has two contributions. First, DLS reports a hydrodynamic diameter that includes the diffuse, hydrated, charged corona, whereas TEM and cryo-EM resolve only the dense, electron-scattering core of the micelle; the more extended the corona, the larger the expected DLS-microscopy difference. Second, intensity-weighted DLS scales with the sixth power of particle diameter, which strongly amplifies the contribution of minor populations of larger oligomers or sub-stoichiometric aggregates - a contribution that is more pronounced for 12D and 12DD, whose TEM size distributions are broader (CV = 24 % and 30 %, respectively) than that of phLCD (CV = 22 %). Together, these two effects account for the DLS/microscopy size difference. The slightly smaller cryo-EM diameter of 12DD relative to 12D is consistent with the higher charge density of its C-terminal block (-24e for 12DD vs. −12e for 12D), which increases electrostatic repulsion within the corona (**Figure S3**) and limits the number of monomers that can be incorporated into a single micelle. The phLCD value falls between those of 12D and 12DD, consistent with the partial protonation of phosphoserines at physiological pH (effective average charge between −1e and −2e per phospho-site), placing the effective C-terminal block charge of phLCD between −12e and −24e. Both phosphomimetics thus reproduce the micellization behavior of the phosphorylated LCD, with the 12DD construct providing a closer match to the fully phosphorylated (-2e per site) limiting case.

Interestingly, cryo-EM imaging of 12DD at elevated concentration (20 µM) additionally revealed occasional thin fibrillar structures coexisting with the dominant spherical micellar population (**Figure 6c**), a feature not observed in cryo-EM micrographs of phLCD or 12D acquired under matched conditions. This minor fibrillar subpopulation was not detected in our TEM experiments performed at lower concentrations (5-15 µM), and we therefore do not include it in the quantitative size analysis of 12DD micelles. We note this observation as a qualitative finding indicating that, while the bona fide phosphorylated LCD effectively suppresses access to fibrillar morphologies even at elevated protein concentration, the doubly-charged 12DD phosphomimetic does not fully abolish this pathway under these conditions - an interesting limitation of phosphomimetic substitution that we revisit in the Discussion.

The CMC was determined for both variants (12D and 12DD) by the pyrene fluorescence method, using the same tangent-intersection approach as shown in **Figure 4c** for phLCD (**Figures 7a**, **7b**). As with phLCD, the I_1_/I_3_ transition was gradual rather than sharp for both 12D and 12DD, consistent with partial hydrophobic exposure of sub-micellar species below the CMC characteristic of intrinsically disordered amphiphiles. The buffer baseline I_1_/I_3_ values were comparable across all three constructs (1.26 for 12D, 1.25 for 12DD, 1.27 for phLCD), confirming equivalent salt-induced compression of the pyrene dynamic range at 150 mM NaCl. The CMC was ∼1.50 µM for 12D and ∼1.93 µM for 12DD - both in the low-micromolar range and close to the CMC determined for phLCD (∼1.38 µM) (**Table S2**). Notably, ΔI_1_/I_3_ was smaller for 12D (0.10) than for 12DD and phLCD (0.14-0.15), indicating a more polar pyrene environment within 12D assemblies. This is consistent with a less well-defined core-corona architecture in 12D, induced by the weakened contrast between the hydrophobic N-terminal block and the charged C-terminal corona. This results in greater water accessibility to the pyrene probe. A logistic function (a sigmoidal fit that describes the gradual transition from monomeric species to micelle formation with increasing concentration) is shown in **Figures 7a** and **7b** for visual guidance only; CMC values were extracted exclusively by the tangent-intersection method.

**Figure 7.**
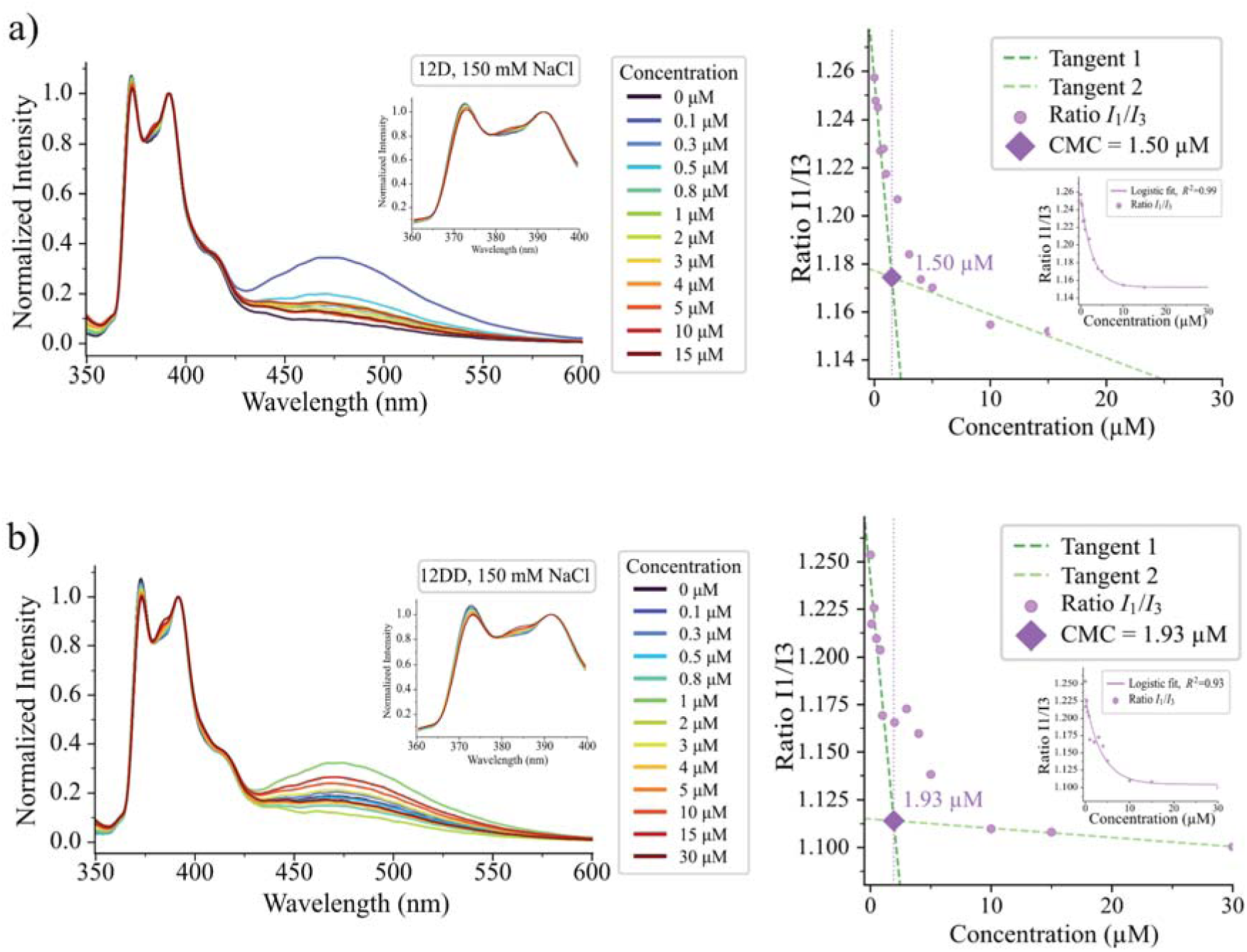
Pyrene CMC determination for 12D and 12DD phosphomimetics. **a)** 12D in physiological buffer (150 mM NaCl). Left: normalized pyrene fluorescence emission spectra (350-600 nm) at 0-15 µM, inset shows zoom (360-400 nm). Right: I_1_/I_3_ ratio versus 12D concentration with tangent-intersection analysis, CMC ≈ 1.50 µM; inset shows logistic fit (R^2^ = 0.99). **b)** Same analysis for 12DD (0-30 µM): CMC ≈ 1.93 µM; logistic fit R^2^ = 0.93.

Collectively, these results demonstrate that the phosphomimetic LCD variants 12D and 12DD reproduce the essential feature of phosphorylation-driven micellization, forming spherical micelles above a low-micromolar CMC with concentration-independent sizes, underscoring the role of C-terminal charge in redirecting self-organization. The additional observation of occasional fibrillar structures in cryo-EM of 12DD at elevated concentration suggests that, while the phosphomimetic substitutions successfully capture the micellization pathway, only the bona fide phosphorylated LCD fully suppresses the alternative fibrillar pathway under the conditions probed.

### 2.4. Salt-dependent morphological landscape: From spheres to worm-like micelles at 300 mM and 500 mM NaCl

As TDP-43, which can also bind RNA, can shuttle between the nucleus and the cytoplasm, it might experience vastly different electrostatic environments.[53] To probe how ionic strength modulates TDP-43 LCD micelle-like assemblies, we examined the phosphorylated form and the phosphomimetic mutants 12D and 12DD under elevated salt conditions (300 mM and 500 mM NaCl). These ionic strengths are higher than the bulk cytosolic value (∼150 mM) but were selected to probe the regime in which electrostatic screening becomes sufficient to alter assembly morphology and to systematically map the morphological landscape accessible to these constructs as a function of charge screening.

Increasing salt concentration screens electrostatic repulsion between the negatively charged residues (phosphate groups or Asp side chains), which is expected to lower CMC and may promote a transition from spherical micelles to more elongated morphologies, such as cylindrical or worm-like micelles. Such behavior is known from non-biological polymers with charged amphiphilic systems, and is driven by the electrostatic screening of repulsive forces between like-charged residues in the corona, allowing for a tighter packing of the hydrophilic blocks.[44, 54, 55]

For phLCD and 12DD, CMC values at elevated ionic strength were determined by the pyrene fluorescence tangent-intersection method described in Section 2.2 (**Figures 8a,b**; **Figures 10a,b**). For phLCD, the CMC remained approximately consistent across salt concentrations, at ∼1.38 μM at 150 mM NaCl, ∼1.6 μM at 300 mM NaCl, and ∼1.45 μM at 500 mM NaCl. For 12DD, the CMC decreased from ∼1.93 µM at 150 mM to ∼0.52 µM at 300 mM and ∼0.89 µM at 500 mM NaCl, confirming the salt dependence of the assembly onset across all constructs (**Table S2**). A logistic function is shown in figures for visual guidance only; all CMC values were extracted by the tangent-intersection method.

**Figure 8.**
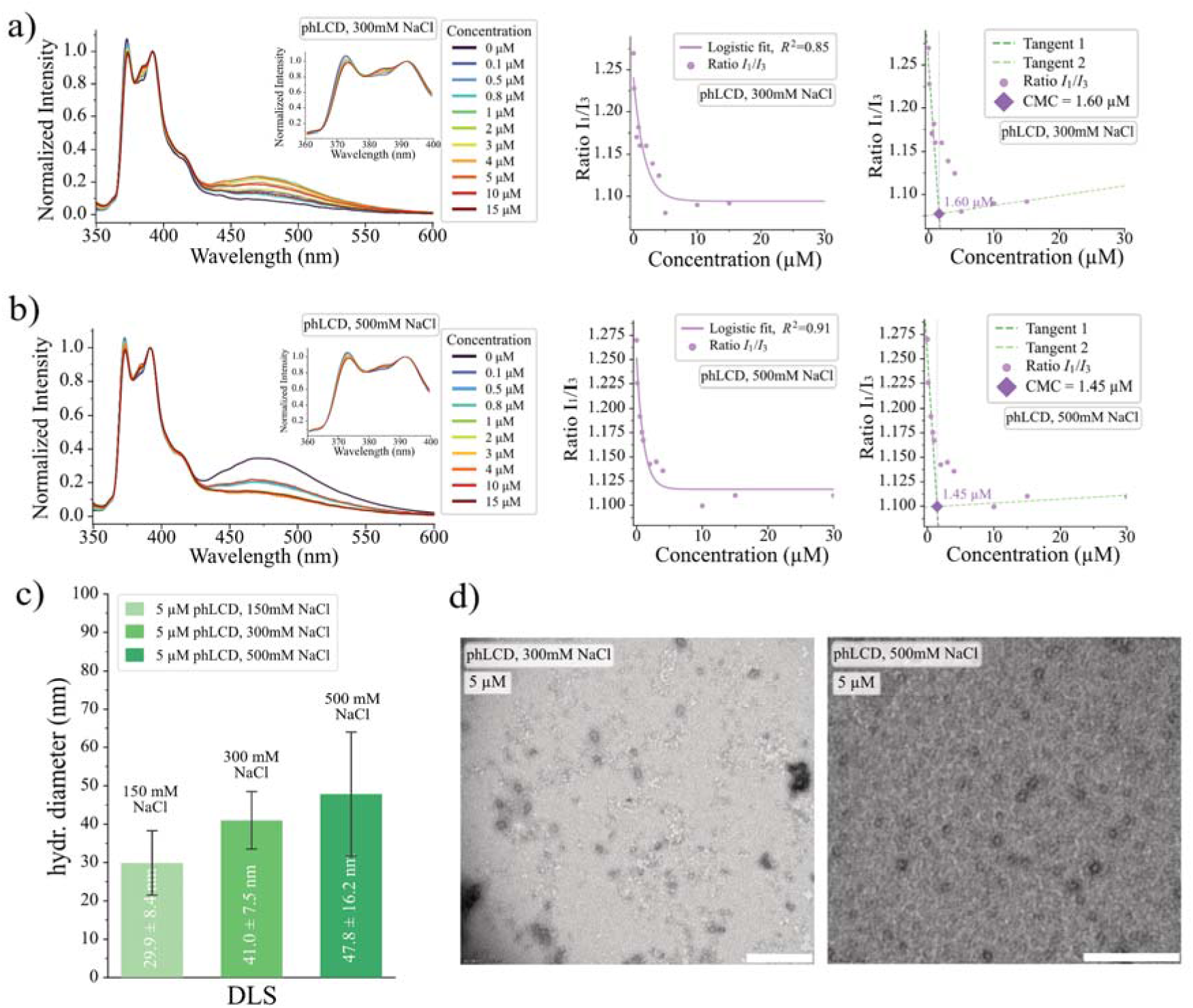
Salt-dependent self-assembly of phosphorylated TDP-43 LCD. **a)** Determination of critical micelle concentration using pyrene. Left: normalized fluorescence emission spectra at increasing phLCD concentrations at 300 mM NaCl buffer. Right: Ratio of pyrene vibronic bands (I_1_/I_3_) plotted against phLCD concentration, fitted with a logistic function (R^2^ = 0.85) and tangent lines indicating CMC ≈ 1.60 μM. **b)** Determination of critical micelle concentration using pyrene. Left: normalized fluorescence emission spectra at increasing phLCD concentrations at 500 mM NaCl buffer. Right: Ratio of pyrene vibronic bands (I_1_/I_3_) plotted against phLCD concentration, fitted with a logistic function (R^2^ = 0.91) and tangent lines indicating CMC ≈ 1.45 μM. **c)** Mean hydrodynamic diameters of 5 μM phLCD determined by DLS at 150, 300 and 500 mM NaCl (mean ± s.d., n = 12). **d)** Representative negative-stain TEM micrographs of 5 μM phLCD at 300 mM and 500 mM NaCl. Scale bar, 300 nm.

DLS measurements at 5 µM (**Figure 8c**) revealed a modest size increase of phLCD micelles with increasing ionic strength (29.9 ± 8.4 nm at 150 mM, 41.0 ± 7.5 nm at 300 mM, and 47.8 ± 16.2 nm at 500 mM NaCl; mean ± s.d., n = 12 biological replicates). This size increase is consistent with electrostatic screening of the corona charges, which allows additional protein chains to be recruited into the micelle, increasing its aggregation number and consequently its diameter. Notably, this corona compaction does not progress far enough to trigger a transition from spherical to worm-like morphology: TEM imaging confirms that only spherical micelles are present at all tested salt concentrations (**Figure 8d**), in marked contrast to 12D and 12DD, which transition to worm-like morphologies at the same ionic strengths (see below; **Figures 9b**, **10d**). A possible explanation for this morphological robustness of phLCD lies in the chemical nature of the phosphoserine side chain itself: the phosphate group is bulky and heavily hydrated, which sterically and entropically resists the close corona packing required to push the packing parameter (*p*) above the spherical-to-cylindrical transition threshold (*p* ≈ 1/3). Aspartate side chains, in contrast, are smaller and less hydrated, permitting the more pronounced corona compaction that drives the sphere-to-worm transition observed in 12D and 12DD at elevated ionic strength.

**Figure 9.**
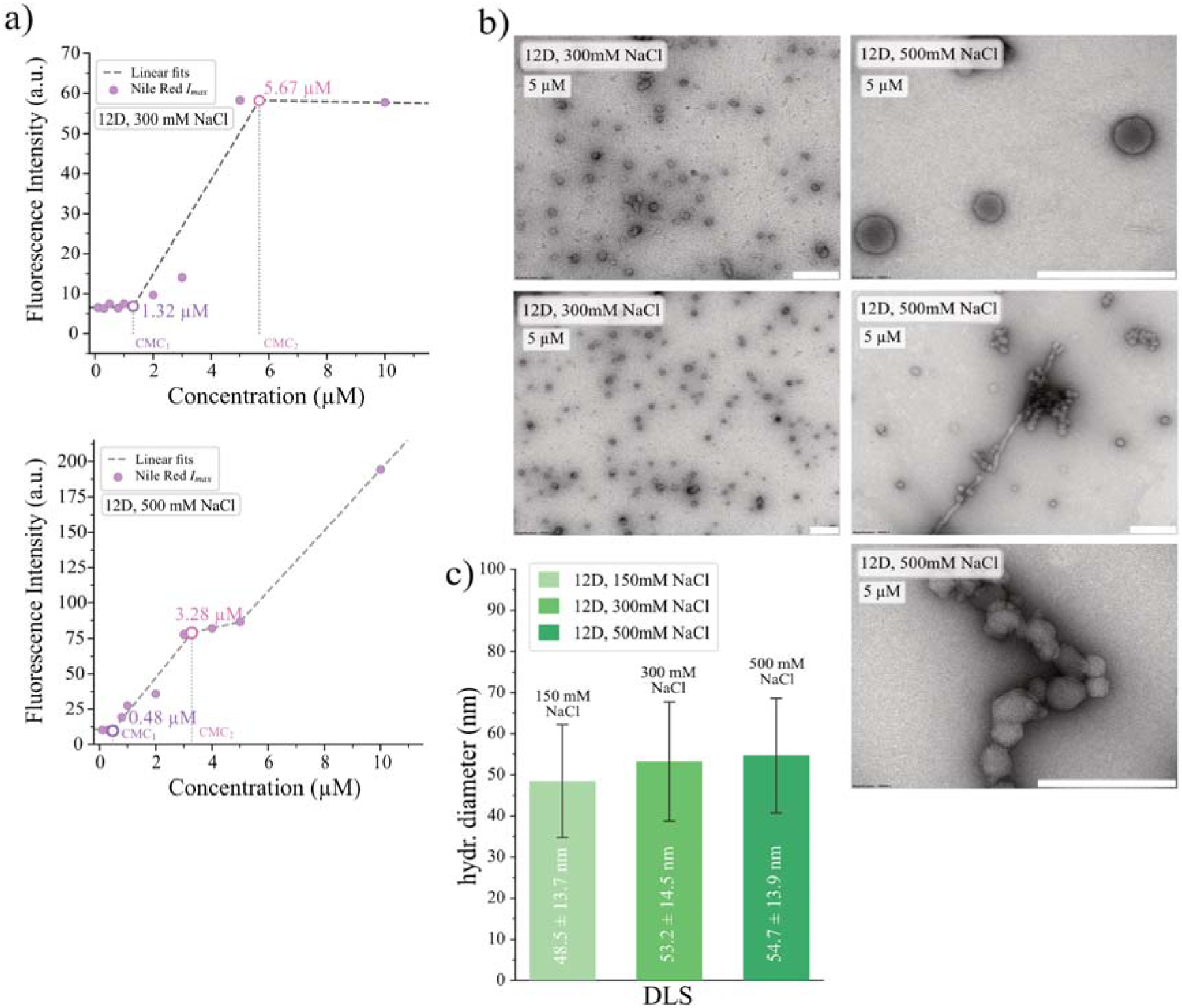
Salt-dependent self-assembly of phosphomimetic 12D TDP43-LCD. **a)** Nile Red fluorescence intensity versus protein concentration for 12D LCD at 300 mM NaCl (top) and 500 mM NaCl (bottom), showing two distinct CMC transitions at high salt **b)** Negative-stain TEM images of 5 μM 12D at 300 mM NaCl (spherical micelles) and 500 mM NaCl (spherical and worm-like structures). **c)** Mean hydrodynamic diameters of 5 μM 12D determined by DLS at 150, 300 and 500 mM NaCl (mean ± s.d., n = 12).

**Figure 10.**
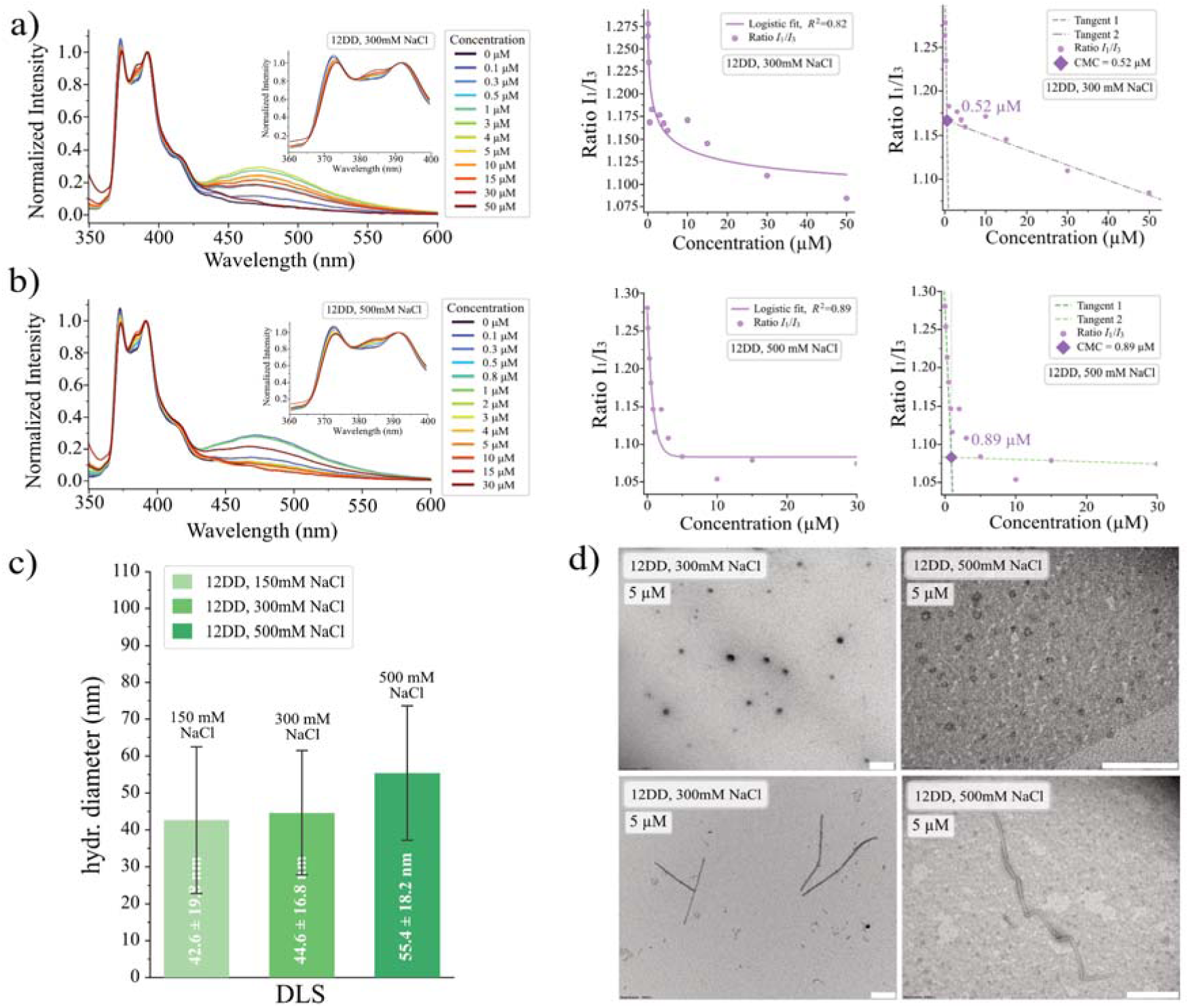
Salt-dependent self-assembly of phosphomimetic 12DD LCD. **a)** Determination of critical micelle concentration using pyrene. Left: normalised fluorescence emission spectra at increasing 12DD LCD concentrations at 300 mM NaCl buffer. Right: Ratio of pyrene vibronic bands (I_1_/I_3_) plotted against 12DD LCD concentration, fitted with a logistic function (R^2^ = 0.82) and tangent lines indicating CMC ≈ 0.52 μM. **b)** Determination of critical micelle concentration using pyrene. Left: normalised fluorescence emission spectra at increasing 12DD LCD concentrations at 500 mM NaCl buffer. Right: Ratio of pyrene vibronic bands (I_1_/I_3_) plotted against 12DD LCD concentration, fitted with a logistic function (R^2^ = 0.89) and tangent lines indicating CMC ≈ 0.89 μM. **c)** Mean hydrodynamic diameters of 5 μM 12DD LCD determined by DLS at 150, 300 and 500 mM NaCl (mean ± s.d., n = 12). **d)** Negative-stain TEM images of 5 μM 12DD at 300 mM NaCl and 500 mM NaCl. Scale bar, 300 nm.

In contrast, 12D displays a markedly different response. Pyrene fluorescence yielded broad and poorly defined I_1_/I_3_ transitions at elevated ionic strength, most likely because the lower charge density of single Asp substitutions (-12e versus −24e for 12DD) reduces the contrast between the hydrophobic N-terminal block and the charged C-terminal corona, yielding micelles with a less well-defined core-corona architecture that fails to sequester pyrene efficiently. We therefore used Nile Red, a solvatochromic dye whose fluorescence intensity increases sharply upon partitioning into hydrophobic environments (excitation ∼550 nm, emission ∼620-650 nm), and which provides a strong signal even for partially hydrated assemblies. The CMC curves indicate a two-step process (**Figure 9a**): a first CMC for formation of spherical micelles (∼1.32 μM at 300 mM and ∼0.48 μM at 500 mM), and a second CMC (∼5.67 μM at 300 mM and ∼3.28 μM at 500 mM) (**Table S2)**, at which micelles elongate into worm-like structures whose cross-sectional diameter remains comparable to that of the spherical micelles, as confirmed by TEM (**Figure 9b**). Mechanistically, we interpret our observations as follows: At 300 mM NaCl, electrostatic repulsion between Asp-rich segments remains sufficiently strong that assembly proceeds stepwise. Upon reaching CMC_1_, relatively small spherical micelles form, with hydrophobic segments partially shielded while charged side chains remain on the surface, maintaining colloidal stability. As protein concentration and/or ionic strength increase, electrostatic screening becomes more pronounced, increasing inter-micelle contacts. This yields a second characteristic point, CMC_2_, beyond which a transition to cylindrical packing becomes favorable: spheres elongate and “fuse” into thin, flexible worm-like micelles. Increasing NaCl to 500 mM particularly enhances this second stage. DLS measurements at 5 µM showed a gradual increase in the intensity-weighted hydrodynamic diameter of 12D assemblies with increasing salt (48.5 ± 13.7 nm at 150 mM, 53.2 ± 14.5 nm at 300 mM, 54.7 ± 13.9 nm at 500 mM NaCl; **Figure 9c**); the relatively modest change reflects the predominantly cross-sectional sensitivity of intensity-weighted DLS to elongating micelles, while the transition to a true worm-like morphology is more clearly visualized by TEM. The size distributions of 12D measured immediately after sample preparation and after 5 days of incubation are essentially identical at all three salt concentrations (**Figure S10b-d**), indicating that the morphologies observed at elevated ionic strength, including the worm-like structures, represent equilibrium states rather than transient kinetic intermediates

The 12DD introduces −24e charge to TDP-43 LCD, twice the amount of 12D. It thus possesses a more hydrophilic and repulsive corona, which requires more salt ions to achieve similar levels of electrostatic screening. The 12DD sample showed a characteristic “delayed” micellization (higher CMC at 150 mM salt) (**Table S2**), followed by formation of rigid, relatively uniform cylindrical assemblies. At moderate ionic strength (∼300 mM NaCl), micelle nucleation occurred at ∼0.5 µM; once this threshold was exceeded, subsequent self-organisation became essentially one-dimensional, with spherical precursors rapidly elongating into short cylinders. In DLS, the spherical micellar precursor population gave rise to an intensity-weighted hydrodynamic diameter distribution with a peak in the ∼10-100 nm range (**Figure 10c**). The cylindrical morphology and dimensions of these assemblies are established directly from negative-stain TEM (**Figure 10d**), which reveals cylinders of cross-sectional diameter ∼20 nm. Further increasing ionic strength to 500 mM NaCl promotes longitudinal growth of these cylinders to lengths exceeding ∼300 nm. Rather than transitioning into a flexible, network-like morphology as in 12D, 12DD instead favours further ordering via end-to-end alignment, without pronounced gelation or a strong increase in macroscopic solution viscosity. We also note that the double DD mutation made the protein chain 12 amino acids longer, and additional effects because of this ∼10% chain length increase, cannot be excluded, although simulations suggest the increase in chain length plays a minor role (**Figures S5 and S6**). However, most of what we measured experimentally can be well explained by the effect of charges. Mechanistically, the behaviour we observed is consistent with Asp mutations exerting a dual effect. During micelle nucleation, they increase the barrier to primary association, requiring higher protein concentration and/or ionic strength to initiate micellization. After a cylindrical core forms, the same groups act as stabilizing “fasteners”: screened carboxylates can form ion pairs and hydrogen bonds that “lock” the lateral surface of the cylinder, preventing twisting and back-conversion (“rounding”) into spheres. Consequently, 12DD forms rigid, narrowly distributed nanocylinders whose length is finely tuned by ionic strength at an essentially constant diameter, making this variant a convenient model for controllable formation of stable rod-like nanostructures.

In order to simulate the effect of salt concentration on micelle size we performed simulations at 150, 300 and 500 mM salt (**Figures S12** and **S13**). We chose the 12DD sequence as a model since it behaves almost identically to TDP-43 LCD 12pS in simulations. We observe the same tendency towards larger aggregation numbers and hydrodynamic diameters with increasing salt concentration (**Figures S12a**, **S13a**, and **S13b**), as reported experimentally for phLCD and 12DD. There is a modest increase of the micelle between 100 and 150 mM salt, followed by a significantly larger assembly at 300 mM. At a 500 mM salt concentration, it is impossible to tell whether the system shifts to macrophase separation or continues to form finite-size assemblies with an aggregation number exceeding the number of chains in the box. Analysis of the contact maps (**Figure S12b**) suggests that electrostatic screening makes the protein less “blocky”, as contacts become more evenly distributed. This is also reflected in the per-block density profiles (**Figure S12c**), where we observe a less segregated internal architecture of the micelles at higher ionic strength (a visual representation of this is given in the cross-section snapshots of the micelles in **Figure S12a**). Analysis of the shape anisotropy (**Figure S13c**) and asphericity parameters offers insight into the effects of electrostatic screening on chain packing. Increased salt concentration significantly flattens the asphericity curves for higher aggregation numbers, suggesting that the penalty for tighter packing of negatively charged amino acids in the corona decreases with electrostatic screening, allowing for larger aggregation numbers (**Figure S13a**). This can also be observed in the contact maps where contacts between the negatively charged C-terminal increase - even if slightly - due to the weakened repulsion (**Figure S12b**). The consequence increase in packing parameters can lead to the formation of wormlike structures. We did not observe worms in simulations, this is not surprising as simulating worms would require a significantly larger number of chains in simulations. However, short-lived oblong-shaped micelles were observed when we gave phosphoserines a charge of −1.5e (**Figure S4b**). This system also exhibits decreased electrostatic repulsion in the corona and is in close analogy to simulations at higher salt concentrations. We ran simulations for the 12pS sequence at 150 mM and 500 mM to confirm that it behaves similarly to 12DD. We observe in **Figures S14** and **S15** that it yields the same phenotypes at the considered salt concentrations, as well as qualitative (and near quantitative) agreement between contact maps (**Figure S14b**), per-block density profiles (**Figure S14c**), size distribution histograms (**Figure S15a**), hydrodynamic diameters (**Figure S15b**) and shape descriptors (**Figure S15c**). Although phosphorylated TDP-43 LCD is chemically distinct from 12DD and exhibits different phenotypes in experiments for higher salt concentrations, this is not captured by simulations. The degree of coarse-graining of the HPS model sacrifices significant details on protein conformations. Phenomena such as steric clashes or protonation/deprotonation of the side chain are not present. Residues are represented as spheres and are fully characterised only by a mass, radius (σ), a stickiness parameter (;\) and a fixed point charge, making 12pS and the 12DD sequences nearly identical species in our simulations.

Although we mostly discussed the spherical to wormlike micelle transition in terms of screened electrostatic repulsion for increasing ionic strength, one cannot discard salting out as a contributor. The increase in the concentration of ions in solution can lead to an effective decrease in affinity between the protein and the solvent. This leads to an increase in effective hydrophobicity, which thus increases the protein’s propensity for supramolecular assembly, leading to tighter packing and potential wormlike micelles. The Debye length changes modestly between 300 mM and 500 mM salt concentration (0.55 nm to 0.43 nm). This represents, however, a significant increase in the number of ions in solution. The spherical-to-elongated morphology transitions observed for both 12D (worms) and 12DD (longer cylinders) between 300 and 500 mM NaCl may therefore be driven by a combination of electrostatic screening and salting out.

### 2.5 Effect of minute amounts of detergent on the morphology of 12D TDP-43 LCD worm-like micelles

The use of very low concentrations of the non-ionic surfactant Tween-20 is common when working with proteins, particularly IDPs, as it is known to reduce surface adsorption. We examined the TDP-43 12D LCD at a protein concentration of 70 μM in a buffer containing 500 mM NaCl (pH 7.5), comparing conditions without and with 0.01% (v/v) Tween-20. It should be noted that, unlike the samples in Section 2.4, which were centrifuged at 20,000 × g for 10 min prior to TEM grid preparation, the samples examined here were applied directly without centrifugation. We chose not to centrifuge our samples to avoid losing large worm-like chains. The two sets of experiments, therefore, differ in both, protein concentration (5 µM versus 70 µM) and sample preparation, and the TEM images should be interpreted with caution. TEM observations revealed an unexpected inversion of morphology: in the absence of Tween-20, spherical micelles dominate, whereas upon addition of the surfactant a large number of small worm-like micelles with lengths of ∼ 100-200 nm form (**Figure 11a** and **11b**).

**Figure 11.**
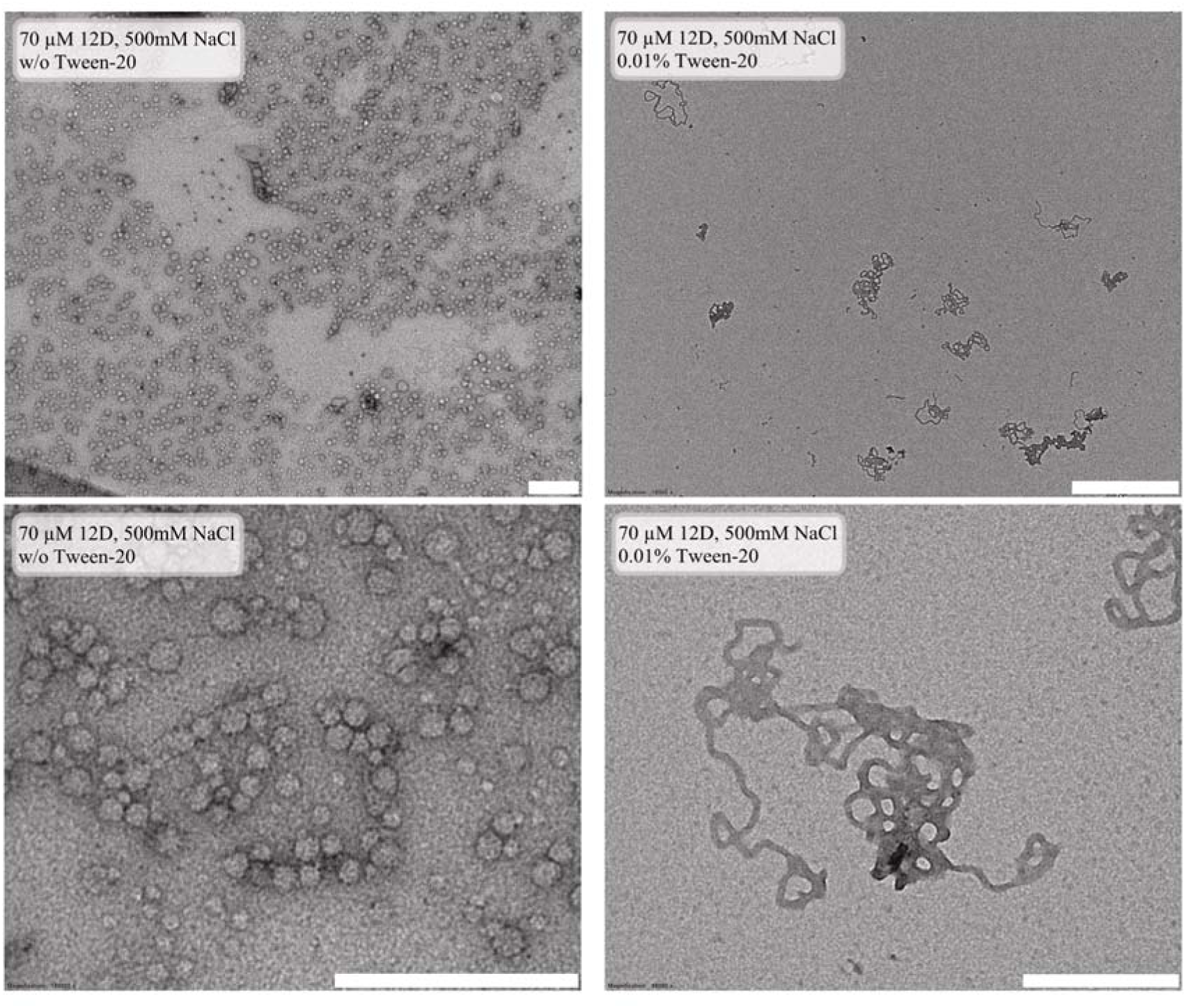
Representative TEM images of 12D assemblies at 70 μM in 500 mM NaCl. **a)** Without Tween-20, showing spherical micelles with a size histogram. Scale bar, 300 nm (top and bottom). **b**) with 0.01% Tween-20, showing small worm-like micelles with histograms of lengths and diameters. Scale bar, 300 nm (bottom), 1µm (top).

A mechanistic interpretation is as follows: The Tween-20 concentration (∼ 0.08 mM) exceeds its own critical micelle concentration (CMC ∼ 0.06 mM), enabling the surfactant to form pure Tween-20 micelles and those, as well as the Tween-20 monomers can also interact with hydrophobic regions of proteins. We speculate that Tween-20 is incorporated into the protein micelles, forming mixed structures in which the hydrophobic tail of Tween-20 (the laurate) inserts into the core of the protein micelle, lowering the packing parameter and promoting a transition from spheres to cylinders. At the same time, the hydrophilic headgroup (polyoxyethylene sorbitan) of Tween-20 strengthens the corona but, in combination with high salt, promotes permeabilization and fragmentation, breaking larger aggregates into many small elongated particles. This results in swelling of spherical assemblies, their elongation, and subsequent growth limitation, preventing the formation of extended networks. Such an effect is typical for low surfactant concentrations, where the surfactant competes for hydrophobic surfaces, increasing the total number of assemblies and making them more dynamic yet length-limited. In contrast to the stabilising action at higher concentrations, 0.01% Tween-20 here acts as a remodelling agent, analogous to its effects on lipid vesicles, where it induces transitions to cylindrical or fragmented morphologies prior to lysis.

We hypothesize that the shift from spherical to wormlike micelles caused by the addition of Tween-20 is linked to changes in the packing of protein chains. Its hydrophobic tail is very short, which would allow it to partition into the spherical micelles without causing too much disruption to the cohesive hydrophobic forces. The hydrophilic block is significantly bulkier - it constitutes roughly 75 % of the molecular mass and has a hydrated size of ∼ 3 nm. Since it is uncharged, the hydrophilic group may serve as a spacer between the negatively charged Aspartate residues in the corona. This decreases electrostatic repulsion, which allows for lower surface curvature in the micelles, which is necessary for the formation of wormlike chains.

The net effect is an increase in *p* toward the worm-like regime (1/3 < *p* < 1/2): endcaps are effectively “capped” by Tween molecules, elongation becomes thermodynamically accessible, and coalescence into spheres is kinetically slowed. The result is a coexistence of spherical micelles and worm-like structures - the latter appearing as relatively straight, moderately thick cylinders, rather than the thin, highly curved worms typical of strongly segregating surfactant systems.[44] This morphological heterogeneity reflects the fact that at 70 µM 12D, the system is poised near the sphere-to-cylinder boundary: Tween-20 shifts *p* into the cylindrical regime for a fraction of the assemblies, while the remaining protein chains retain sufficient head-group area to sustain spherical geometry.

## 3. Discussion

Nanostructure formation in IDPs has emerged as an exciting frontier in protein biophysics. In particular, subcritical nanoclusters and well-defined, finite-sized assemblies have recently garnered attention for their unique structural and functional properties.^[24, 55, 56]^ In this work, we showed that TDP-43 LCD organises into micelles upon C-terminal phosphorylation. This behaviour arises from the block-copolymer architecture conferred by clustered phosphosites, which create a hydrophilicity asymmetry between the N-terminal hydrophobic block and the charged C-terminal block. Coarse-grained simulations indicated that micellization is favoured under physiological conditions, and these predictions were confirmed by *in vitro* experiments for the phLCD. The phosphomimetic variants 12D and 12DD qualitatively recapitulate this trend. Notably, phLCD micelles proved morphologically robust, retaining spherical geometry across a range of ionic strengths. The phosphomimetic variants, by contrast, undergo salt-dependent morphological transitions, from spheres to worm-like micelles (12D) or rigid nanocylinders (12DD), consistent with charge-controlled packing parameter effects, and illustrating that it is the precise magnitude and chemical nature of C-terminal charge, rather than its mere presence, that determines the accessible morphological space.

Although we restricted ourselves to the study of TDP-43 LCD, we conjecture that these findings may extend to full-length phosphorylated TDP-43, as it has been shown to be one of the main contributors to macro PS in the full-length protein.^[3, 4]^ Our findings expand the known manifestations of TDP-43 LCD supramolecular assembly by adding a phosphorylation-dependent micelle state and a related worm-like morphology. C-terminal fragments of TDP-43 of 20 - 25 kDa that comprise mainly the LCD are consistently found in ALS/FTD patients ^[7, 10, 57]^ suggesting that our model system may have a naturally occurring analogue in disease-relevant contexts.

The role of phosphorylation in TDP-43 pathology is not fully understood. C-terminal phosphorylation is a well-established occurrence in ALS and FTD.^[25, 26]^ Whether that is a driver or antagonizer of pathological TDP-43 aggregation, or simply a secondary effect, remains an open question and is highly debated.^[27, 58, 59]^ Recent studies have challenged the aggregation-inducing role of phosphorylation, rather positioning it as a protective regulator against amyloid formation by suppressing PS and TDP-43 insolubility *in vitro* and in cells.[35] PS has been suggested as a precursor of fibrilization in proteins, therefore getting a more complete understanding of its multiple drivers is key to answering that process. Recent findings from Cryo-EM have resolved the structure of TDP-43 fibrils isolated from ALS/FTD brains.^[7, 8, 60]^ They indicate that the amyloid core is formed by amino acids 282-360, roughly equivalent to our blocks 1 and 2. In our phosphorylated TDP-43 LCD model, they are positioned within the micelle interior, whereas block 3 resides on the micelle surface. This structural arrangement indicates that the amyloid core is strongly compositionally similar to the simulated core of our phosphorylated micelle. One can speculate that the phosphorylated micelle may serve as a precursor to amyloid fibrils, highlighting its potential clinical relevance.^[18, 23]^ Experimentally, however, phosphorylated TDP-43 LCD micelles remained non-fibrillar and colloidally stable over days, in marked contrast to the unphosphorylated sequence, which matures into fibrils.

Our findings also highlight the power of coarse-grained simulations as a predictive tool for protein-protein interactions. Through them we identified micellization as a potential form of assembly for phosphorylated TDP-43 LCD, which was later confirmed by experiments. This shows that even very simplified models can capture intricate molecular phenomena happening in IDRs. It is nonetheless necessary to remark the limitations of such techniques, we did not observe micellization in the 12D construct in our simulations, and 12DD and 12pS (phLCD) micelles remained stable at high salt concentrations *in vitro*, while simulations seemed to hint at a shift to the formation of macroscopic droplets. Whether or not these discrepancies were caused by the force field or by our limitation in the number of chains in the simulation box is beyond the scope of this work. There was also a quantitative mismatch between simulated and experimental hydrodynamic diameter sizes, where our simulated hydrodynamic diameters are roughly half of those measured in the experiment. This is likely attributed to the excessive “stickiness” of the HPS model, which may overestimate the inner density of the micelles. While newer iterations of this force field exist which address this issue, none have been consistently applied to TDP-43 LCD like HPS.

Our work also sheds some light on the usefulness of phosphomimetic mutations in the study of phosphorylation. The real charge of phosphoserines is pH dependent and oscillates between −1e and −2e, with a preference for −2e. This means that the phosphorylated LCD lies somewhere between the 12D and 12DD constructs. Remarkably, we found that the size of the phosphorylated LCD micelles lies between those of the 12D and 12DD mutants at physiological salt. We designed point mutations in the phosphomimetics to reproduce the average phosphorylation pattern derived from computational and experimental studies. The distribution of phosphorylation sites from kinase activity might not be uniform and vary across individual proteins. In addition, phosphoserines are slightly larger than aspartates and thus aspartates may underestimate steric clashes. This supports the interpretation of 12D and 12DD as reliable surrogates for the negatively charged C-terminal block of the phosphorylated LCD. Although the agreement with the phosphomimetics in our study is remarkable, it should be taken with caution. We did not, for instance, observe the formation of worm-like structures in the phosphorylated LCD, although they were observed in both 12D and 12DD at high enough concentrations. We emphasize that these worm-like morphologies were observed exclusively under *in vitro* conditions of elevated ionic strength (300-500 mM NaCl) or in the presence of a non-ionic surfactant (0.01% Tween-20), and we make no claim regarding their accessibility within the cellular environment; whether comparable morphologies can form *in vivo* under specific local conditions remains an open question that will require dedicated cellular studies. Furthermore, cryo-EM imaging at elevated protein concentration (20 µM) revealed that the 12DD construct retains residual access to a fibrillar pathway, with thin fibrillar structures occasionally coexisting with the dominant spherical micellar population, a feature not observed for phLCD or 12D under matched conditions. This indicates that, while phosphomimetic substitutions successfully redirect TDP-43 LCD assembly toward micellization, only the bona fide phosphorylated LCD appears to fully suppress access to fibrillar morphologies at the concentrations probed, suggesting that the precise chemistry of the phospho-group, beyond its formal charge alone, may contribute to the kinetic stabilisation of the micellar state.

We identified electrostatics as the main driver of the conformational assemblies in this work. Results from simulation capture the electrostatic screening induced by the increase in ionic strength, which lead to decreased repulsion between the negatively charged residues forming the micelle’s coronas (**Figure S12b**). The mechanisms that drive the transition from spherical to wormlike micelles in 12D and 12DD is, however, not fully elucidated. While electrostatic screening is definitely necessary for such a charged molecule, it may not be sufficient. To fully describe this phenomenon we would also need to take into account secondary effects like salting out. This is however beyond the scope of this work.

TDP-43 LCD micellization is driven by phosphorylation-induced hydrophobicity asymmetry, with clustered phosphosites creating a charged C-terminal block. This patterning is observed in full-length TDP-43 and other IDRs, suggesting micellization may be widespread. Simulations and experiments showed consistent spherical shapes, core-corona organization, and concentration-independent sizes above CMC, validating the approach.

Although Tween-20 is not a naturally occurring cellular component, the observation that 0.01% (v/v) of this surfactant dramatically remodels the assembly landscape of 12D TDP-43 LCD, converting a predominantly spherical population into well-defined worm-like micelles, highlights the *in vitro* sensitivity of these assemblies to amphiphilic co-solutes. We note that this observation does not establish that comparable morphologies are accessible *in vivo*, and we present it cautiously as a starting point for future experiments. Whether endogenous amphiphilic molecules with related physicochemical properties, such as certain lipids or fatty acids, could exert analogous effects within the cell remains an open and speculative question that we did not test here and that would require dedicated experiments in physiologically relevant systems. That said, TDP might be exposed to dramatically different cellular environments in the course of its multiple functions. Furthermore, the amount of Tween that makes the difference is very low. Most important, however, is to realise that such structures are accessible within the phase diagram of TDP-43 LCD.

TDP-43 is implicated in neurodegenerative diseases, such as ALS, FTD and AD. Despite decades of intense research, the actual pathological species in these diseases is still debated. For similar proteins, much early work focused on understanding how amyloid fibres might emerge from self-templating and primary nucleation, and then propagate through secondary nucleation.^[61, 62]^ For TDP-43, also non-fibrillar, non-amyloid architectures have been observed in patients and model systems^[9, 63, 64]^, suggesting that other assembly forms may be pathologically relevant and not only the amyloid state might contribute to nuclear TDP-43 loss-of-function. Since PS has been suggested as a relevant mechanism in biology, the initial belief was that its high concentration in the dense state might contribute to and facilitate pathological aggregation and potential amyloid formation. More recently, droplets have also been suggested to stabilise the protein, while aggregation phenomena seem largely catalysed at the droplet/buffer interface.^[23, 65, 66, 67]^ To understand the role of TDP-43 aggregation, we need to know all states.

How this phospho-switch operates in the context of the full-length protein remains a particularly intriguing open question, especially with regard to the architecture of the resulting assemblies. Shinn et al. recently demonstrated that unphosphorylated full-length TDP-43 already adopts a microphase-separated, size-limited assembly of ∼12 molecules forming ordered structures of ∼33 nm in diameter, driven by inter-block attractions and repulsions encoded in its multi-domain architecture.[24] In their model, the N-terminal oligomerization domain (NTD) and the two RNA-recognition motifs (RRM1 and RRM2), together with the LCD, act as the blocks of an effective block copolymer, with NTD-driven head-to-tail homo-oligomerization and the documented stacking of the NTD onto the RRMs likely contributing a structural inner subdomain, while the LCD provides a separately segregating hydrophobic subdomain driven by its aromatic, aliphatic and helix-forming residues. Importantly, the 12 phosphorylation sites we examined here lie within residues 369-414 of full-length TDP-43, deep within the LCD and well beyond the conserved helix and the hydrophobic LCD core. Phosphorylation therefore introduces a strongly negatively charged segment (shifting the local net charge from approximately +3 to approximately −21, depending on phosphoserine protonation) at the very C-terminal tip of the LCD, without altering the chemistry of the folded NTD, the RRMs, the NLS, or the conserved helix.

Based on the block-copolymer logic established here for the isolated LCD and on the architecture reported by Shinn et al., and acknowledging that TDP-43 full length is a complex multidomain protein, we can speculate that phosphorylation of the full-length protein would reinforce, rather than disrupt, the existing microphase architecture: the NTD/RRM module would likely remain at the interior of the assembly as a structural kernel; the hydrophobic N-terminal half of the LCD would form an intermediate hydrophobic shell surrounding this kernel; and the phosphorylated C-terminal tail of the LCD would project outward as a densely charged outer corona. This “kernel-shell-corona” architecture differs in topology from the simpler core-corona micelles of the isolated phosphorylated LCD reported here, but follows the same underlying physical principle of phosphorylation-induced charge patterning driving block segregation. Several testable consequences follow: the assembly size should remain comparable to, or modestly larger than, the ∼33 nm reported by Shinn et al.; the aggregation number is likely to decrease as the charged corona increases steric and electrostatic crowding at the surface, in line with the trend we observe between 12D and 12DD; access to the LCD-driven amyloid pathway should be further suppressed, as the hydrophobic LCD segments become sequestered behind both the NTD/RRM kernel and the phospho-corona; and RNA binding to the RRMs, now embedded inside the assembly, may be sterically restricted, suggesting a potential link between C-terminal phosphorylation and modulation of TDP-43 RNA-processing function. Each of these predictions provides a concrete starting point for future biophysical and structural studies on phosphorylated full-length TDP-43.

For understanding TDP pathology, it is important to know all supramolecular states that can, in principle, be formed due to its physicochemical characteristics. This study added two more supramolecular states in the LCD domain observed only for the charged variants, i.e. the spherical micelle as well as worm-like micelles. While there is much to do to connect and understand all these different states and their assembly pathways and how they can form and transition into each other, our study is fully in line with a view that by shifting TDP-43 LCD assembly from phase-separated droplets to stable micelles, one can suppress - or at least significantly delay - amyloid fibril formation, hence phosphorylation may offer a therapeutic avenue to mitigate pathological aggregation.

This discovery is particularly exciting because it challenges traditional views of protein structure-to-function, revealing that IDP’s dynamic nanostructures - like micelles - may be key to biological regulation and disease.

## Experimental Section/Methods

### 4.1. Protein Expression and Purification

#### 4.1.1 Protein Expression and Purification of TDP-43 LCD

The TDP-43 LCD (residues 267-414) was expressed in *E. coli* BL21(DE3) cells from the expression vector pJ411 (Addgene #98669) encoding a His-TEV-TDP-43 LCD fusion protein. Cells were grown in a total culture volume of 12 L Luria-Bertani (LB) medium containing 30 mg L[¹ kanamycin at 37°C, 180 rpm until an OD_600_ of approximately 0.7 was reached. Cultures were cooled to 18°C and protein expression was induced with 0.5 mM IPTG. Expression proceeded overnight for approximately 16-18 h at 18 °C with shaking at 180 rpm.

Cells were harvested (8 000 × *g*, 15 min, 4 °C), resuspended in ice-cold lysis buffer (4 M guanidine hydrochloride (Gdn-HCl), 20 mM Tris-HCl pH 8.0, 500 mM NaCl, 10 mM imidazole, 4 mM β-ME, 1 mM PMSF) and lysed by sonication with a Branson Sonifier (4 cycles of 4 min at 40 % amplitude) with 4 min cooling intervals between cycles to prevent overheating. Cell lysates were cleared by centrifugation (20000 × *g*, 1 h at 4 °C). The supernatant was then incubated for 2 h at room temperature with 10 mL HisPur™ Ni-NTA resin (Thermo Fisher Scientific) under gentle rotation. Beads were washed with 30 CV washing buffer (4 M Gdn-HCl, 20 mM Tris-HCl pH 8.0, 500 mM NaCl, 30 mM imidazole) and protein was eluted with elution buffer (4 M Gdn-HCl, 20 mM Tris-HCl pH 8.0, 150 mM NaCl, 500 mM imidazole).

The eluate was buffer-exchanged into 0.5 M Gdn-HCl (50 mM Tris-HCl pH 8.0, 150 mM NaCl) using an Amicon Ultra-15 centrifugal filter unit (3 kDa MWCO, Merck Millipore) and then cleaved overnight at room temperature with His-TEV protease (1 : 20 w/w). Guanidine hydrochloride was then added to a final concentration of 4 M to dissolve precipitate. The solution was again incubated with fresh Ni-NTA to bind residual His-tag and TEV protease for 2 h, and the flow-through containing untagged TDP-43 LCD was collected. The flow-through containing untagged TDP-43 LCD was spin-concentrated using Amicon Ultra-15 centrifugal filter (3 kDa MWCO, Merck Millipore) and subjected further purified by size-exclusion chromatography on a Superdex 200 10/300 column using an ÄKTA system (Cytiva). The column was equilibrated and run in a buffer containing 4 M Gdn-HCl, 20 mM Tris-HCl, 150 mM NaCl, pH 8.0. Fractions were analyzed by SDS-PAGE using 4-12% gradient gels with MES running buffer and stained with Coomassie Brilliant Blue, and peak fractions with sufficient purity were pooled and spin-concentrated to 2.6 mM. Protein concentration was determined using a bicinchoninic acid (BCA) assay with bovine serum albumin (BSA) as a standard. The purified protein was aliquoted, flash-frozen in liquid nitrogen and stored at -80°C.

A single-cysteine variant of TDP-43 LCD (A366C) was generated from the WT TDP-43 LCD plasmid (pJ411, Addgene #98669) by site-directed mutagenesis using forward and reverse mutagenic primers and Platinum™ SuperFi™ II DNA Polymerase (Thermo Fisher Scientific), introducing a cysteine residue in place of alanine at position 366 in the otherwise cysteine-free WT background. The A366C variant was expressed and purified following the same protocol as for the TDP-43 LCD, with the addition of 1 mM DTT to the size-exclusion chromatography buffer to maintain the cysteine in its reduced state.

Constructs encoding His-TDP-43 LCD 12D and 12DD were generated from the His–TDP-43 LCD construct in the pJ411 vector obtained from Addgene (plasmid #98669). D and DD substitutions have been placed for S at amino acid position 373, 375, 379, 387, 389, 393, 395, 403, 404, 406, 409, 410, based on Gruijs da silva et al. work.[35] We purified similarly to the LCD described above, except that all buffers were adjusted to pH 7.5.

### 4.1.2 Protein Expression and Purification of CK1δ

Chemically competent *E. coli* (Rosetta 2(DE3) pλPP, ^[68]^ were transformed with pET vectors containing MBP-His6-TEV-CK1δ and untagged λPP-phosphatase and grown in 4 to 8 L TB media (Formedium) containing 100 mg/l Ampicillin at 37°C, 140 rpm until an OD_600_ of 0.6. Cultures were cooled on ice to 18°C and protein expression was induced using 0.5 mM IPTG with further incubation at 18°C, 140 rpm for 21 h. Cells were harvested by centrifugation (4000 × *g*, 15 min), re-suspended in ice cold lysis buffer (50 mM Tris-Cl, 500 mM NaCl, 10 mM imidazole, 5 mM MgCl_2_, 1 mM TCEP, 5% glycerol, 1X Roche EDTA-free cOmplete protease inhibitor, 100 U/ml Millipore Benzonase, pH 8.0)) and lysed by high pressure homogenization at 1.9 kbar (Constant Systems CF1 cell disruptor). Cell lysates were cleared by centrifugation (40000 × *g*, 30 min at 4°C). Lysate containing MBP-His6-TEV-CK1δ was applied to a HisTrap FF 5 ml column (Cytiva), using an automated chromatography system (Biorad NGC Quest Plus, used for all chromatography steps). The column was washed with 20 CV (column volumes) of wash buffer (50 mM Tris-Cl, 500 mM NaCl, 10 mM imidazole, 5 mM MgCl_2_, 5 % glycerol). Tagged CK1δ was eluted from the Ni-NTA column by applying a linear gradient of 10-500 mM imidazole in wash buffer over 10 CV. Elution fractions containing MBP-His6-TEV-CK1δ were dialyzed (50 mM Tris-Cl, 200 mM NaCl, 5 mM MgCl_2_, 1 mM DTT, 5 % glycerol) in the presence of His6-TEV protease (1:50 w/w) at 4°C overnight. The dialyzed solution was supplemented with NaCl (500 mM), imidazole (10 mM) and passed over a HisTrap FF 5 ml column to separate untagged CK1δ (eluting at 25 mM imidazole) and MBP-His6, as well as His6-TEV protease (eluting at imidazole concentrations higher than 50 mM imidazole). Untagged CK1δ were spin-concentrated (Amicon Ultra 15 3K, Merck Millipore) and subjected to size exclusion chromatography (Superdex 200 16/600 pg, Cytiva, 50 mM Bis-Tris pH 6.5, 50 mM NaCl, 10 mM MgCl_2_, 0.5 mM TCEP). Peak fractions were pooled and spin-concentrated to 120 µM, aliquoted, flash-frozen in liquid nitrogen and stored at -80°C.

### 4.2. Site-Specific Fluorescent Labeling of TDP-43 LCD

For fluorescence microscopy experiments, the WT A366C variant of TDP-43 LCD (expressed and purified as described in Section 4.1.1) was site-specifically labeled using thiol-reactive maleimide chemistry.

Purified A366C LCD was first reduced by incubation with 10 mM 1,4-dithiothreitol (DTT) at room temperature for 10 min, and subsequently buffer-exchanged into maleimide labeling buffer (4 M guanidine hydrochloride, 1× PBS, 0.1 mM EDTA, 0.2 mM TCEP, pH 7.0) using Amicon Ultra centrifugal filters (3 kDa MWCO; Merck Millipore) with five sequential wash cycles. Labeling was performed overnight at 4 °C with LD655-maleimide (Lumidyne Technologies) at a 1:2 protein-to-dye molar ratio. The dye was first dissolved in DMSO and then added to the protein solution while gently vortexing to ensure rapid mixing and prevent precipitate formation.

After incubation, the reaction was quenched by addition of DTT to a final concentration of 10 mM for 5 min. Free dye and DTT were removed by seven rounds of buffer exchange against labeling buffer (without TCEP) using Amicon Ultra centrifugal filters (3 kDa MWCO; Merck Millipore), followed by size-exclusion chromatography on a Superdex 200 10/300 column equilibrated in storage buffer (4 M guanidine hydrochloride, 20 mM Tris-HCl, 150 mM NaCl, pH 8.0). Fractions corresponding to monomeric labeled protein were pooled and analyzed by SDS-PAGE. The final labeled A366C LCD was concentrated to 22.3 µM in a storage buffer, aliquoted (3 µL), flash-frozen in liquid nitrogen, and stored at -80 °C until use.

### 4.3. In Vitro Phosphorylation Reaction Assay

Recombinant TDP-43 LCD (residues 267-414) samples at various concentrations were prepared independently and phosphorylated *in vitro* using recombinant human casein kinase 1δ (CK1δ) in the appropriate proportions for each concentration. The reactions were carried out separately in kinase buffer (50 mM HEPES pH 7.5, 150 mM NaCl, 10 mM MgCl_2_, 0.75 mM ATP, 1 mM DTT) at a 1:10 molar ratio of CK1δ to LCD. Samples were incubated at room temperature overnight (16 h).

Phosphorylation of TDP-43 was assessed by SDS-PAGE followed by Western blotting using a phospho-specific antibody against TDP-43 (anti-pSer409/410, Proteintech, 80007-1-RR). Samples were separated on 4-12% Bis-Tris gradient gels (Invitrogen, NP0329BOX) using 1×MES running buffer (Invitrogen, NP0002). Proteins were transferred to nitrocellulose membranes (Cytiva, 10600001) by wet transfer at 400 mA for 60 min in transfer buffer containing 192 mM glycine (Roth, 0079.5), 24.8 mM Tris (Roth, 0188.4), and 20% ethanol (Merck, 1.00983.1000).

Membranes were incubated overnight at 4 °C with the primary antibody, followed by incubation with IRDye 800CW donkey anti-rabbit secondary antibody (LI-COR, 926-32213). Fluorescence signals were detected using an Odyssey CLx Infrared Imaging System (LI-COR).

Total protein staining was performed on gels following electrophoresis. Gels were fixed for 30 min at room temperature in 50% ethanol (Merck, 1.00983.1000) and 10% acetic acid (Sigma, A6283), stained overnight with SYPRO Ruby (Sigma, S4942), and destained for 30 min in 10% ethanol and 10% acetic acid. Images were acquired using a ChemiDoc MP Imaging System (Bio-Rad) (**Figure S8**).

### 4.4. Sample preparation

Throughout this work, “physiological buffer” refers to a buffer containing 150 mM NaCl at neutral pH (HEPES pH 7.5 or Tris pH 7.5) at 25 °C. The use of HEPES for phLCD reflects the requirements of the CK1δ kinase reaction (Mg^2+^, ATP); the simpler Tris-based buffer is used for 12D and 12DD, where the kinase reaction is not required.

Protein stocks (2.3-2.6 mM in 4 M Gdn-HCl) were thawed on ice immediately before use. For samples at final concentrations ≥10 μM, the stock was diluted directly into the respective assay buffer. For samples at concentrations below 10 μM, the stock was first pre-diluted into an intermediate buffer containing 2 M Gdn-HCl (with the same components and pH as the SEC buffer used for each construct) to achieve a suitable intermediate concentration, from which the required volume was then diluted into the assay buffer to reach the desired final concentration. Each concentration point was prepared as an independent sample. In all cases, the residual Gdn-HCl concentration in the final sample did not exceed 20 mM. This dilution protocol was applied consistently across all experiments, including DLS, TEM, pyrene fluorescence, Nile Red fluorescence, and *in vitro* phosphorylation reactions, and was used for both the phosphorylated LCD (phLCD) and the phosphomimetic variants (12D and 12DD).

Buffer compositions were construct-specific. For the phosphorylated LCD (phLCD), phosphorylation reactions were performed in 50 mM HEPES, pH 7.5, 150 mM NaCl (or adjusted to 300 or 500 mM as indicated), 10 mM MgCl_2_, 0.75 mM ATP, and 1 mM DTT. The reactions were not quenched and were incubated overnight (16 h) at room temperature. Following incubation, samples were centrifuged at 20,000 × *g* for 10 min at 22°C to remove large aggregates and debris. The supernatants were then transferred to low-binding 1.5 mL reaction tubes (Sarstedt, Nümbrecht, Germany; cat. no. 72.690.001) and kept until measurements.

For phosphomimetic variants (12D and 12DD), samples were diluted into 20 mM Tris pH 7.5, 150 mM NaCl (or varied to 300/500 mM as indicated) and incubated overnight (16 h) at room temperature. Following incubation, samples were centrifuged at 20,000 × *g* for 10 min at 22°C to remove large aggregates and debris, then transferred to low-binding 1.5 mL reaction tubes (Sarstedt; cat. no. 72.690.001) prior to measurements.

For surfactant experiments, assay buffers were prepared separately with the same components as above but with Tween-20 added to a final concentration of 0.01% (v/v) from a 10% stock solution. Protein samples were diluted directly into these Tween-20-containing buffers, followed by overnight incubation (16 h) at room temperature. Samples were then centrifuged and transferred as described above. All samples were prepared freshly and used within 24 h.

For TEM and Cryo-EM sample preparation in Sections 2.2-2.4, samples were centrifuged at 20,000 × g for 10 min at 22°C prior to grid preparation. For Section 2.5, samples were applied directly without centrifugation to preserve the full assembly population.

### 4.5. Thioflavin-T (ThT) aggregation assay

Wild-type (WT) and *in vitro* phosphorylated TDP-43 LCD (15 μM final concentration) were incubated with 5 μM ThT in assay buffer (50 mM HEPES pH 7.5, 150 mM NaCl, 10 mM MgCl[, 0.75 mM ATP, 1 mM DTT) at 25 °C. Measurements were set up in a sealed black 96-well plate (Corning, low-binding) to minimize evaporation and non-specific binding. ThT fluorescence was monitored every 60 min for 72 h using a plate reader (excitation 440 nm, emission 480 nm) with 3 s orbital shaking before each reading. Data are expressed as fold change relative to the fluorescence intensity at t = 0 after subtraction of buffer-only background. All experiments were performed in at least three independent replicates.

### 4.6. Turbidity Assay

To compare the phase-separation propensities of phosphorylated and unphosphorylated TDP-43 LCD, turbidity measurements were performed in parallel on three samples: (i) CK1δ-phosphorylated TDP-43 LCD (phLCD); (ii) “aged” unphosphorylated TDP-43 LCD (non-phLCD prepared in parallel with phLCD without kinase and ATP, allowed to age for the same duration); and (iii) “freshly” prepared non-phLCD.

#### 4.6.1 Well passivation

Prior to sample loading, the wells of a clear-bottom 384-well plate used for measurement were passivated to minimize non-specific protein adsorption to the polystyrene surface and to prevent surface-induced heterogeneous nucleation of phase separation. Each well was filled to the brim with 10% (w/v) Pluronic and incubated at room temperature for approximately 1 hour to allow self-assembly of a PEO-based hydrophilic coating on the polystyrene surface. The Pluronic solution was then aspirated, and each well was washed 10 times with the respective experimental buffer to remove any non-adsorbed Pluronic prior to sample loading.

#### 4.6.2 Sample preparation and measurement

The purified TDP-43 LCD was rapidly diluted from the 4 M GdmCl storage buffer into the experimental buffer to a final protein concentration of 50 µM. For phLCD, the buffer composition was 50 mM HEPES pH 7.5, 150 mM NaCl, 10 mM MgCl_2_, 0.75 mM ATP, 1 mM DTT, with CK1δ added at a 1:10 molar ratio (kinase:LCD). For the “aged” non-phLCD sample, the buffer composition was identical except that CK1δ and ATP were omitted to control for buffer effects.

Immediately after dilution, 30 µL of each freshly prepared phLCD and “aged” non-phLCD sample was transferred to separate passivated wells and incubated in the plate at 23 °C. After approximately 1 hour of incubation, the “fresh” non-phLCD sample was prepared (diluted from 4 M GdmCl into the same buffer) and immediately transferred to adjacent passivated wells of the same plate. Buffer-only controls matching the composition of each sample were included in parallel wells for blank subtraction.

Turbidity was then measured for all samples simultaneously by recording absorbance at 350 nm using a SpectraMax iD5 plate reader (Molecular Devices) at 23 °C, with a 3 s plate shaking step immediately preceding the measurement. Under these conditions, phLCD and “aged” non-phLCD samples had been incubated for ∼1 hour, while the “fresh” non-phLCD sample was measured immediately after dilution. Blank absorbance values from buffer-only wells were subtracted from the corresponding sample wells.

All measurements were performed with three biological replicates (independent protein preparations) and three technical replicates per biological replicate. Reported values represent the mean ± standard deviation across biological replicates.

### 4.7. Fluorescence imaging of TDP-43 LCD phase separation

To directly visualize the phase separation behavior of unphosphorylated and phosphorylated TDP-43 LCD, time-lapse fluorescence imaging was performed on a Yokogawa spinning disk confocal microscope.

#### 4.7.1 Sample chamber passivation

To minimize protein adsorption to the glass surface and to suppress surface-induced nucleation of phase separation, the wells of an ibidi µ-Slide 15 Well 3D Glass Bottom (formerly µ-Slide Angiogenesis Glass Bottom; ibidi GmbH, Munich, Germany; Cat. No. 81507; #1.5H glass coverslip bottom) were passivated prior to sample loading. Each well was filled to capacity with 10 % (w/v) Pluronic F-127 (Sigma-Aldrich) and incubated for 1 h at room temperature, allowing self-assembly of a poly(ethylene oxide)-based hydrophilic coating on the glass surface. The Pluronic solution was then aspirated, and each well was washed seven times with the corresponding experimental buffer to remove non-adsorbed Pluronic prior to sample loading. Wells were used immediately after passivation.

### 4.7.2 Sample preparation

For fluorescence visualization, LD655-labeled TDP-43 LCD A366C (prepared as described in Section 4.2) and unlabeled wild-type TDP-43 LCD were co-diluted from their respective stocks directly into the assay buffer to reach final concentrations of 10 nM labeled A366C and 50 µM unlabeled LCD (labeled-to-unlabeled molar ratio ≈ 1:5,000). All samples were prepared freshly immediately before imaging.

Unphosphorylated samples. Labeled and unlabeled protein were co-diluted into assay buffer containing 50 mM HEPES pH 7.5, 150 / 300 / 500 mM NaCl (as indicated), 10 mM MgCl_2_, and 1 mM DTT (no kinase, no ATP). These samples also served as the no-kinase/no-ATP control for the phosphorylated condition.

Phosphorylated samples. Labeled and unlabeled protein were co-diluted into assay buffer containing 50 mM HEPES pH 7.5, 150 mM NaCl, 10 mM MgCl_2_, 0.75 mM ATP, and 1 mM DTT, supplemented with recombinant CK1δ at a 1:10 molar ratio (kinase:LCD) to initiate phosphorylation. The reaction mixture was immediately transferred to a passivated well of the µ-Slide 15 Well 3D and incubated for 1 h at room temperature in the dark on the microscope stage (under a light-shielding cover, laser off), during which the CK1δ-catalysed phosphorylation reaction proceeded to near-completion. Image acquisition was then started (t = 0 = end of phosphorylation incubation) to capture the assembly behavior of the fully phosphorylated state.

All incubation and imaging were performed at room temperature.

#### 4.7.3 Image acquisition

Imaging was performed on a Yokogawa CSU-W1 SoRa spinning disk confocal scanner unit (Yokogawa Electric Corporation; 50 µm pinhole disk) mounted on an inverted Nikon Eclipse Ti2 microscope frame, equipped with a Nikon CFI Plan Apochromat Lambda S 60× silicone oil immersion objective (NA 1.30; Nikon part number MRD73600) and a Hamamatsu ORCA-Flash 4.0 sCMOS camera (2,048 × 2,048 pixels; 6.5 µm physical pixel pitch). Samples were excited with a 660 nm laser line at 40 % nominal output power. The unit was operated in standard W1 mode without SoRa pixel reassignment engaged. Single-plane fluorescence images were acquired at the bottom of the ibidi chamber with an exposure time of 175 ms per frame. The calibrated pixel size at the specimen level was 0.108 µm/pixel, providing a field of view of 222 × 222 µm. For each condition, time-lapse acquisitions consisting of 61 frames were recorded at 60 s intervals over 60 min. Images were saved as 16-bit OME-TIFF files using NIS-Elements software (Nikon).

#### 4.7.4 Image processing

Acquired time-lapse stacks (61 frames per condition) were processed in Fiji (ImageJ v1.53, NIH) for visualization. Grayscale 16-bit image stacks were re-coloured by applying the “Red” lookup table (LUT), and brightness and contrast were adjusted linearly for display purposes (identical settings applied across all frames of a given stack). A calibrated scale bar (using the recorded pixel size of 0.108 µm/pixel) and a time stamp (in min format) were overlaid onto each frame using the built-in Fiji tools (Analyze → Tools → Scale Bar; Image → Stacks → Time Stamper). For all conditions, t = 0 of the time stamp corresponds to the first acquired frame (defined per condition in Section 4.7.2). Processed stacks were exported as AVI movies for presentation. No quantitative image analysis was performed; the imaging served as direct optical confirmation of TDP-43 LCD phase separation and coalescence kinetics in the unphosphorylated state, as well as direct evidence for the suppression of phase separation in the phosphorylated state.

### 4.8. Native Mass Spectrometry

Native mass spectrometry was performed on a Micromass Q-ToF Ultima mass spectrometer (Waters Corporation, Milford, USA) modified for transmission of high masses (MS Vision, Almere, Netherland).[69] For this, the buffer of 50 µL protein solution was exchanged against 200 mM ammonium acetate using 10 kDa MWCO Amicon ultra centrifugal filters (Merck Millipore, Billerica, USA) according to the manufacturer’s instructions. The protein was then diluted with 200 mM ammonium acetate to a final concentration of 8 to 10 µM. For each measurement, 4 µL of the protein solution were loaded into gold-coated glass capillaries prepared in-house[70] and the sample was directly introduced into the mass spectrometer. The following parameters were used for data acquisition: capillary voltage 1.7 kV, sample cone voltage 80 V, RF lens voltage 80 V, collision voltage 40 V. Mass spectra were externally calibrated using 100 mg/mL cesium iodide solution and processed using MassLynx 4.1 (Waters Corporation, Milford, USA). Mass spectra were analyzed using Massign software (version 11/14/2014).[71]

### 4.9. Dynamic Light Scattering (DLS)

DLS measurements were performed on a Malvern Zetasizer Nano ZS instrument at 25 °C using backscattering geometry (detection angle 173°). Protein samples (100 μL, concentration range 0.1-70 μM) were centrifuged at 20,000 × *g* for 10 min to remove possible large aggregates or dust particles, then loaded into disposable UV-grade plastic micro cuvettes (BRAND GMBH, Cat# 759200, 70 µL, 8.5 mm centre height). Samples were equilibrated for 5 min prior to measurement. For each measurement, the instrument performed up to 15 acquisitions that were automatically averaged by the Zetasizer software with automatic attenuation and optimal cell position selection.

Size distributions were obtained from intensity-weighted analysis using the General Purpose algorithm of the Malvern Zetasizer software. A lower size limit of 1 nm was applied to exclude sub-nanometre size contributions arising from instrumental noise at the shortest lag times (∼0.1-5 µs) of the autocorrelation function. In this region, the correlation coefficient plateau displayed low-amplitude fluctuations characteristic of statistical and electronic noise rather than scattering from physical particles; these fluctuations were otherwise interpreted by the analysis software as a spurious sub-nanometre size population. The 1 nm threshold removes this artefact while preserving all real scattering signals from protein assemblies. Given the inherent polydispersity of the samples, mean hydrodynamic diameters were determined from the peak maximum corresponding to the micellar population in the intensity-weighted distribution (typically 10-100 nm range). The identity of this peak as the protein micellar species was independently confirmed by TEM and Cryo-EM imaging. Larger species (>100 nm), occasionally present in the distributions, were attributed to residual aggregates or dust contributions and were excluded from the analysis. Concentration- and ionic-strength-resolved DLS size distributions for phLCD, 12D and 12DD across the full tested range are provided in **Figures S16 - S18.**

Statistical analysis was performed at two levels of replication. For each biological replicate, 3-5 technical replicates (independent measurements of the same protein preparation in separate cuvettes) were averaged to obtain a single mean hydrodynamic diameter. A biological replicate represents an independent protein preparation including a new protein purification batch, fresh ATP stock, and freshly prepared buffer. Reported values represent the mean ± standard deviation (SD) calculated across n = 12 biological replicates per condition.

### 4.10. Transmission Electron Microscopy (TEM)

Protein samples were applied to carbon-coated copper grids (Quantifoil® Carbon Support Films, Cu 300 mesh, Cat# Q38936; Quantifoil Micro Tools GmbH, Großlöbichau, Germany). Prior to sample application, grids were glow-discharged using an Emitech K100X glow discharge at 20 mA for 60 s to render the carbon film hydrophilic. 5 µL of sample was applied to the carbon-coated side of the grid and incubated for 30 s. Excess liquid was removed by blotting with filter paper. The grid was then stained with 5 µL of 2 % (w/v) uranyl acetate for 30 s, blotted dry, and air-dried before imaging.

For TEM sample preparation in Sections 2.2-2.4, samples were centrifuged at 20,000 × g for 10 min at 22 °C prior to grid preparation to remove large aggregates. For Section 2.5 (Tween-20 experiments), samples were applied directly without centrifugation to preserve the full assembly population.

Grids were examined on a FEI Tecnai 12 BioTwin transmission electron microscope operated at 120 kV acceleration voltage, equipped with an EMSIS Makiya CMOS camera (2,448 × 2,048 pixels; EMSIS GmbH). Images were recorded at magnifications of 10,000-50,000×. Particle diameters were measured manually in Fiji (ImageJ v1.53, National Institutes of Health) on >200 particles per condition.

### 4.11. Cryo-Electron Microscopy (Cryo-EM)

Cryo-EM samples of phLCD, 12D, and 12DD were prepared at a final protein concentration of 20 µM following the same protocol used for all biophysical measurements in this study (Section 4.4). Protein was diluted from 4 M Gdn-HCl stock into the respective experimental buffer (phLCD: 50 mM HEPES pH 7.5, 150 mM NaCl, 10 mM MgCl_2_, 0.75 mM ATP, 1 mM DTT; 12D and 12DD: 20 mM Tris pH 7.5, 150 mM NaCl) and incubated overnight at room temperature to allow assembly equilibration. Immediately prior to grid preparation, samples were centrifuged at 20,000 × *g* for 10 min at 22 °C to remove large aggregates, and the supernatant was used directly for vitrification.

C-flateTM holey carbon grids (CF-2/2-2C, 2 µm hole diameter with 2 µm spacing, 200 mesh Cu; Protochips) were glow-discharged in air immediately before use using an Emitech K100X glow discharge unit at 20 mA for 60 s to render the carbon film hydrophilic. A 3.5 µL aliquot of sample was applied to the glow-discharged side of the grid mounted in a Thermo Fisher Scientific Vitrobot Mark IV operated at 20 °C and 100 % relative humidity. Grids were blotted with Whatman No. 1 filter paper using an asymmetric blotting configuration (a single filter paper on the back side of the grid and two stacked filter papers on the sample side; blot force 1, blot time 3 s, wait time 5 s, drain time 0 s, single blot per grid), and immediately plunge-frozen in liquid ethane cooled by clean (ice-free) liquid nitrogen. Vitrified grids were stored under liquid nitrogen until imaging.

Cryo-EM imaging was performed on the same FEI Tecnai 12 BioTwin transmission electron microscope (Thermo Fisher Scientific) operated at 120 kV and equipped with an EMSIS Makiya CMOS camera (2,464 × 2,056 pixels; EMSIS GmbH) as used for negative-stain TEM (Section 4.10). Data were acquired in low-dose mode using a three-step scheme: grid screening in search mode at 1,900× nominal magnification (spot size 1, intensity 48.70); focusing on an adjacent area in focus mode at 150,000× nominal magnification (spot size 3, intensity 36.88, focus distance 3.32 µm, focus angle 16.0°); and image acquisition in exposure mode at 68,000× nominal magnification (spot size 1, intensity 40.07) with an exposure time of 1 s. The beam diameter at the fluorescent screen was 165 mm. Imaging was performed under low-dose conditions to minimize beam-induced damage to the vitrified samples. Micrographs were inspected and analyzed in Fiji (ImageJ v1.53, NIH).

### 4.12. Fluorescence Assays for Critical Micelle Concentration (CMC)

#### 4.12.1 Pyrene Assay

CMC determination by pyrene fluorescence was carried out on a Duetta dual fluorescence/absorbance spectrometer (Horiba Scientific). A 3 mM pyrene stock solution (Sigma-Aldrich, ≥ 99 %) was prepared in acetone at 3 mM stock concentration. Working samples covering the desired protein concentration range were prepared in an assay buffer with 2 µM pyrene ^[49, 50, 51, 52]^ the final acetone content was kept below 0.5 % (v/v). Samples were equilibrated for 15 min at room temperature before measurement.

Spectra were recorded at λ_ex_= 335 nm with emission scanned from 350 to 600 nm (excitation bandpass 5 nm, emission increment 0.5 nm, integration time 0.05 s, detector accumulations 1). The I_1_/I_3_ ratio was obtained from the maxima at 373 ± 1 nm (I_1_) and 384 ± 1 nm (I_3_). Buffer blank spectra recorded under identical conditions were subtracted prior to analysis. Raw Duetta ASCII files were first collated in Microsoft Excel and then analysed with custom Python 3.9 notebooks in Google. Scripts (pandas, numpy) extracted intensities, computed I_1_/I_3_, performed linear fits (numpy.polyfit), calculated the intersection analytically, and generated publication-ready plots (matplotlib).

The CMC was estimated by the tangent-intersection method: linear regressions were fitted to (i) the gradually descending region at low protein concentrations and (ii) the lower plateau at high concentration; their intersection on the abscissa was taken as the apparent CMC. Linear fits were performed using numpy.polyfit, and the intersection point was calculated analytically as x_CMC_ = (b_2_ – b_1_)/(k_1_ – k_2_), where k_1_, b_1_ and k_2_,b_2_ are the slopes and intercepts of the two regression lines, respectively. A sigmoidal (logistic) fit is shown in figures solely for visual guidance. Because the protein is a true amphiphile with a hydrophobic core and charged termini, the absence of a flat pre-micellar baseline is attributed to partial hydrophobic exposure of monomeric and oligomeric species below the CMC, consistent with the intrinsically disordered nature of the LCD. Protein-based amphiphiles display a gradual I_1_/I_3_ transition-reflecting heterogeneous hydrophobic-patch exposure along flexible chains-such that the value obtained represents the onset of cooperative micellization rather than a sharp critical point. The elevated ionic strength of the physiological buffer (20 mM Tris, 150 mM NaCl, pH 7.5) further compresses the observable dynamic range of the I_1_/I_3_ transition by lowering the aqueous baseline value from ∼1.87 (pure water) to ∼1.25. The pyrene fluorescence data therefore serve as corroborating evidence for the CMC, which was independently established by DLS and TEM (Sections 2.2 and 2.3)

For construct 12D at 300 mM NaCl and 500 NaCl the I_1_/I_3_ curve lacked a clear inflection, precluding reliable CMC extraction; in this case the Nile Red assay (Section 2.4) was used instead.

#### 4.12.2 Nile Red Assay

CMC determination by Nile Red fluorescence was carried out on a Duetta dual fluorescence/absorbance spectrometer (Horiba Scientific). A 1 mM Nile Red stock solution (Sigma-Aldrich) was prepared in ethanol. Working samples covering the desired protein concentration range were prepared in an assay buffer with 2 µM Nile Red^[72, 73]^; the final ethanol content was kept below 0.5 % (v/v). Samples were equilibrated for 30 min at room temperature before measurement. Spectra were recorded at λ_ex_ = 550 nm with emission scanned from 560 to 700 nm (excitation bandpass 5 nm, emission increment 0.5 nm, integration time 0.05 s, detector accumulations 1). Buffer blank spectra recorded under identical conditions were subtracted prior to analysis.

Nile Red is a solvatochromic dye that is weakly fluorescent in aqueous solution but undergoes a marked increase in emission intensity and a blue shift of the emission maximum (∼15-25 nm) upon incorporation into a hydrophobic micellar core.^[73, 74]^ The CMC was determined by the three-tangent intersection method applied to the fluorescence intensity vs. protein concentration profile: linear regressions were fitted independently to (i) the low-concentration baseline region, (ii) the steeply rising transitional region, and (iii) the upper plateau at high protein concentrations. The intersections of adjacent regression lines defined CMC_1_ (onset of micelle formation) and CMC_2_ (completion of the micellar transition).[75]

Raw spectra were processed using the same custom Python 3.9 scripts in Google Colaboratory as described for the pyrene assay (Section 4.12.1); peak intensities were extracted at the emission maximum, linear fits were performed using numpy.polyfit, and intersection points were calculated analytically as x_CMC_ = (b_2_ – b_1_)/(k_1_ – k_2_), where k and b denote the slope and intercept of each regression line. All analysis scripts are available upon request.

### 4.13. Simulations

All of our simulations were carried out with the HOOMD-blue package, version 3.11.0.[76] We used gsd 2.9.0 to record trajectories to binary files. Non-bonded interactions were modeled with the HPS force field [39] using a Debye length of κ = 1.0 nm^-1^ (∼ 100 mM NaCl), except in the simulations where we tested the effect of salt where we selected KAPPA using the relation 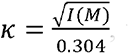, where I(M) represents the salt concentration in moles/L.[44] We utilized the corresponding bonded parameters of 3.8 Å for bond length and 10[kJ/Å² for the spring constant. Polymer chains were initialized in a 3D “snake” configuration on a cubic lattice, with consecutive monomers separated by one bond length. As the chain zigzags along the lattice, each directional change is accompanied by a perpendicular displacement of 2 monomers instead of one to ensure no overlap happens.

Simulations were carried out at 300 K, with the temperature being kept constant using the Langevin thermostat, with a drag coefficient of 10.0 m_u_/ps (m_u_ = atomic mass unit). Simulation boxes were cubic, with dimensions chosen to keep a fixed volume_fraction. The timestep was set to 10[fs and trajectories were output at intervals of 1[ns.

For the equilibration step we start by thermalizing all particles at temperature 150 K and the timestep to 10^-4^ fs. We create a spherical wall to confine chains to the center of the box and induce assembly. The wall is centered at the origin and its radius is chosen so as to give the confined region a volume fraction of 0.01. We perform a short 500 fs step where non-bonded interactions are progressively turned-on. This is followed by a 740 fs step where temperature is progressively increased to 300 K. We then progressively bring the timestep up to 10 fs in a longer step that runs for 1.82 x 10^7^ timesteps. We then run the simulation for an extra 280 ns to further relax chains. We then delete the spherical walls and run the simulation for 10 ns. The simulation is then run for 7.78 μs, we use the last 7 μs as the production run. Convergence to a steady state was verified by tracking the size of the largest cluster over the course of the simulation, and the production window was defined only after this quantity had stabilised around a mean value (**Figure S19**).

#### 4.13.1. Molecular dynamics simulations

We performed preliminary molecular dynamics simulations for the 4 sequences included in this work (TDP-43 LCD, 12D, 12DD and 12pS). Simulations contained 150 chains each, with a fixed volume fraction of 0.005. We performed 3 replicas for each sequence. Analysis of MD trajectories are shown in **Figures S20 - S24**, they are obtained by combining trajectories from all 3 replicas for each sequence.

#### 4.13.2. Semi-grand canonical monte carlo simulations

Most simulations used in this paper were carried out using an in-house semi-grand canonical monte carlo algorithm, developed as a plugin for HOOMD-blue. We used it in combination with molecular dynamics in order to obtain more accurate results. In this method, protein chains can be inserted or deleted from the simulation every few MD steps. Since adding/removing particles from the box can cause memory issues our simulations are composed of a mix of interacting and non-interacting chains. Non-interacting are composed of ideal beads with no non-bonded interactions. An insertion move consists of “turning on” non-bonded interactions, whereas a deletion move consists of turning them “off”. Trial configurations are biased and selected by the Rosenbluth algorithm.[77] This method was developed for chain molecules and creates trial configurations monomer by monomer, selecting orientations with higher Boltzmann weights, which severely punishes unphysical conformations. In order to respect detailed balance we modified our acceptance criteria.

This approach is more well suited for the study of finite-sized assemblies like micelles. It minimizes finite-sized effects that arise from the small number of chains and yields more accurate micelle sizes and dilute phase concentrations. It also has the added advantage of exhibiting long-range diffusion, which allows the simulation to reach a steady state much faster despite its low overall density. Our implementation has a finite-sized reservoir of chains, which corresponds to a semi-grand canonical ensemble instead of a true grand canonical one.

It is important to highlight a subtle but significant difference between the semi-grand and grand canonical ensemble simulations. In grand canonical simulations, one should not see phase separated regions - only one phase or the other. The reason why we see droplets coexisting with dilute chains in systems that would otherwise phase separate is that the maximum number of chains is constrained in the semi-grand canonical ensemble.

We attempt a monte carlo move every 316 timesteps (3.16 ps). We chose this value because each monte carlo move takes around the same time to complete as 10^2.5^ ≈ 316 MD timesteps to complete. We therefore spend around the same amount of time on MD as we do on MC. If a move is accepted we conserve the individual velocities of each particle.

Simulations were initialized with 30 interacting chains in proximity, confined by a spherical wall. The non-interacting chains are initialized near the edges of the box, outside of the region confined by the spherical walls. The equilibration procedure is the same as the one described for the MD simulations.

To confirm the validity of our method we compared our results to equivalent MD simulations (**Figure S21**). We obtained nearly identical size distribution histograms (**Figure S21, left**) for the two micellizing sequences. The histograms do not overlap perfectly since they were not normalized by the overall number of chains, as that is not a fixed number in SGCMC. However, our fit yields a nearly identical size distribution for the micelles. The hydrodynamic diameters obtained in both methods are nearly indistinguishable (**Figure S21, right**)

#### 4.13.3. Analysis

We conducted the post analysis of our simulations using python. For semi-grand canonical monte carlo simulations we ignored non-interacting chains. Some of our analysis requires convergence of the simulation to a stationary state. This is the case for the size distribution histograms. To ensure that convergence had been reached in the distribution of cluster sizes we analyzed the size of the largest cluster over time (**Figure S19**). We consider that convergence has been reached once the size of the largest cluster has stabilised around a mean value.

##### 4.13.3.1. Clustering

All of our analyses rely in some way or another on identifying clusters of protein chains. We performed this step by using the clustering algorithm from the freud python package (v 2.13.2).[78] We consider 2 chains to be in the same cluster when at least one pair of amino acids from each is separated by a distance of 7.0 Å or less.

#### 4.13.3.2. Size distribution

To compute size distribution histograms we utilized the clustering data from each frame. We computed the cluster size histograms for each frame and averaged them over all of them. That histogram is then multiplied term-wise by the vector [1, 2, … , N-1, N]. The distribution now reflects the probability of observing a chain in state M, which represents a chain located in a cluster of size M. In systems exhibiting micelle formation this quantity is known to follow the distribution:

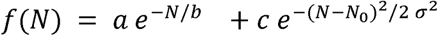

Where the first term represents a dilute polydisperse solution and the second one represents the distribution of micelle sizes of mean aggregation number N_o_ and standard deviation a. By fitting our data with this function we are able to extract the mean number of chains in the simulated micelles shown in **Figure 2d** and which are used to define the micelle size interval in other analyses.

We chose to discard the first 2μs of the simulation for the calculations of cluster sizes to ensure convergence had been reached. We thus end up with 6μs per replica to compute size distribution histograms.

##### 4.13.3.3. Hydrodynamic diameter

The hydrodynamic radius (*R_H_*) of a polymer or polymer assembly can be computed by the relation:

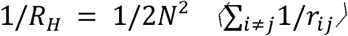

Where N represents the number of monomers in the polymer/cluster and *r_ij_* the distance between monomers *i* and *j*. Computing *r_ij_* for every single pair of monomers across every single frame in the simulation is computationally prohibitive. To speed-up calculations we restrict ourselves to computing a fraction s of all amino acid pairs for each frame. We chose *s* = 10^-s^. For reference, in a cluster containing 20 chains of TDP-43 LCD 12pS that means computing *r_ij_* for 86 out of the ∼ 8.6 million pairs of residues. In each frame pairs are picked at random, making it so that by performing the same calculations throughout multiple frames assures a representative sample or inter residue distances within a cluster and yields a representative mean value of the real hydrodynamic radius. The obtained value is thus multiplied by 2 to obtain the hydrodynamic diameter.

##### 4.13.3.4. Per-block density profiles

To compute our per-block density profiles we used an approach used in one of our group’s previous works.[56] We start by taking the center of mass (COM) of each cluster which is provided by Freud’s clustering algorithm. We compute each of the constituent particles’ distance from the COM. We group these values by sequence position (0 to N-1), which gives us the ensemble of distances from COM for each of the residues in the sequence.

We defined each of the 3 blocks by dividing the sequence into 3 - nearly - identically long segments. The constituent residues for each block in each sequence is summarized in the table below. Block 3 contains all the phosphoserines or equivalent phosphomimetic substitutions (located between 107-144). Block 3 is longer in the 12DD sequence due to this sequence being slightly longer than the others, while blocks 1 and 2 are identical through all sequences.

##### 4.13.3.5. Contact maps

Contact maps were computed for each cluster in each simulation frame. We chose to focus on only interchain contacts since they show a clearer picture of contacts inside IDP assemblies without the very prominent near-diagonal terms that come from intrachain contacts. We start by using freud’s locality module to build a neighbor list for each amino acid. The list consists of all other amino acids located at a distance 1.5a from each other, where a is the mean value of the Ashbaugh-Hatch parameter a of the two. We then compute the mean number of contacts for each pair, which are plotted in the upper diagonal contact maps.

The lower diagonal part of the contact map represents interchain contacts between pairs of blocks. They are easily computed from by averaging per-residue pair contacts computed previously. We chose to multiply this value by the number of amino acids to have the mean number of interchain contacts that each amino acid in block X forms with amino acids in block Y. For blocks of different sizes we multiplied the value by *n* = 1/2 * (*n_x_* + *n_y_*).

#### 4.14 Shape descriptors

The parameters characterizing the shape of clusters were obtained from the corresponding gyration tensors. For each individual cluster obtained from our clustering algorithm, we computed the gyration tensor using the freud python package (v 2.13.2).[78] Its principal moments (the diagonal terms of the diagonalized tensor) are represented as 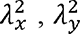 and 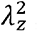, where 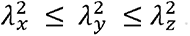. The parameters can be computed from the relations below:

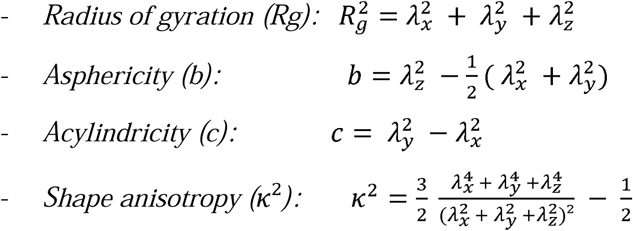

#### 4.15. Sequence property profiles

The plots in **Figure 2a** were made with python. Hydrophobicity values were taken from the λ parameter of the HPS force field.[39] We used a sliding window of size 25 to smooth the values across neighboring residues. We defined the x coordinate as the central position of the sliding window and omitted the first 12 and last 13 residues.

### 4.16. Statistical analysis

#### 4.16.1 Sample sizes and replication

All experimental measurements were performed using independent biological replicates, defined as independent protein preparations (separate purification batches, freshly prepared buffers, and, for phosphorylated samples, fresh ATP stocks and independent kinase reactions). Dynamic light scattering (DLS) measurements were performed with n = 12 biological replicates per condition. For each biological replicate, 3-5 technical replicates (independent measurements of the same protein preparation in separate cuvettes) were averaged to obtain a single mean hydrodynamic diameter; each individual measurement comprised up to 15 instrument runs, automatically averaged by the Zetasizer software to yield a single autocorrelation function per acquisition. Turbidity and Thioflavin-T (ThT) fluorescence assays were each performed with n = 3 biological replicates, with 3 technical replicates per biological replicate. For negative-stain TEM and cryo-EM, particle sizes were measured for >200 particles per condition (TEM) and >100 particles per condition (cryo-EM), pooled across multiple grid regions and independent imaging sessions performed on separate days, comprising both independent protein preparations and separate grids prepared from shared protein stocks.

#### 4.16.2 Data processing and presentation

For turbidity and ThT measurements, buffer/blank values were subtracted from all readings prior to analysis. For DLS, technical replicates were averaged to yield a single mean hydrodynamic diameter per biological replicate, and reported values represent the mean ± standard deviation (SD) across the n = 12 biological replicates; mean diameters were extracted from the peak maximum of the intensity-weighted size distribution (General Purpose algorithm). For electron microscopy, particle sizes were measured manually in Fiji (ImageJ v1.53, National Institutes of Health) following spatial calibration from the microscope pixel-size metadata; for spherical micelles the particle diameter was measured, whereas for worm-like micelles the cross-sectional diameter (width) was measured perpendicular to the long axis at several positions along each structure and averaged. Critical micelle concentrations were determined from logistic fits and the tangent-intersection method, with the coefficient of determination (R^2^) reported for each fit. Particle-size dispersity is reported as the coefficient of variation (CV = SD/mean). Sub-1 nm DLS signals attributable to instrumental noise and >100 nm signals attributable to residual aggregates or dust were excluded as described in the DLS Methods section.

#### 4.16.3 Software

Data were processed and plotted in Python (NumPy, SciPy, Matplotlib), with DLS size distributions computed using the Malvern Zetasizer software. Quantitative results are expressed as mean ± SD with the indicated number of replicates; no formal null-hypothesis significance testing was applied, as the study characterises self-assembly behaviour through direct physical measurements rather than statistical group comparisons.

## Acknowledgements

We thank the laboratories of AG Lemke and AG Dormann, in particular Joana Caria, and Saskia Hutten, for helpful advice and laboratory support. We also thank Tobias Simone for technical support and assistance with TEM measurements and instrument settings, as well as the Tecnai 12 TEM project. We acknowledge funding by the Deutsche Forschungsgemeinschaft (DFG, German Research Foundation) within the Collaborative Research Center SFB1551 “Polymer Concepts in Cellular Function” (Project No. 464588647), including support from its internal service projects for protein production at the IMB Protein Production Core Facility and protein analytics at the MPI Core Facility. We also acknowledge the Z02 service project for access to TEM infrastructure. E.A.L. acknowledges funding from the European Research Council (ERC Advanced Grant “MultiOrganelleDesign”). D.D. acknowledges funding from the DFG Heisenberg Programme (Project ID 442698351). This work was supported by the Max Planck Computing and Data Facilities (MPCDF).

## Author Contributions

**R.F.D**., **A.L**., **H.R.** contributed equally to this work and share first authorship. **R.F.D**.: Conceptualization, Methodology, Software, Formal analysis, Data curation, Visualization, Writing - original draft, Writing - review and editing. **A.L.:** Conceptualization, Methodology, Validation, Formal analysis, Investigation, Data curation, Visualization, Writing - original draft, Writing - review and editing. **H.R.:** Methodology, Formal analysis, Writing - review and editing. **T.S.:** Methodology, Writing - review and editing. **Si.Mo**.: Methodology, Visualization, Writing - review and editing. **E.P.:** Methodology, Writing - review and editing. **J.B.:** Methodology, Formal analysis, Visualization**. F.S.-D.:** Methodology, Resources, Writing - review and editing. **K.L.:** Methodology, Resources, Writing - review and editing. **C.S.:** Methodology, Resources, Writing - review and editing. **M.M.M.:** Methodology, Resources, Writing - review and editing. **Sv.Mor**.: Methodology, Writing - review and editing. **F.S.:** Methodology, Writing - review and editing. **D.D.:** Methodology, Resources, Writing - review and editing. **L.S.:** Conceptualization, Methodology, Writing - review and editing. **M.G.:** Conceptualization, Methodology, Software, Resources, Supervision, Project administration, Funding acquisition, Writing - review and editing. **E.A.L.:** Conceptualization, Methodology, Resources, Supervision, Project administration, Funding acquisition, Writing - review and editing.

## Data Availability Statement

The experimental data that support the findings of this study are available from the corresponding authors upon reasonable request. The molecular dynamics simulation trajectory files are deposited in the Edmond repository of the Max Planck Society (https://doi.org/10.17617/3.A4AM5H), and the associated simulation and analysis code is available at https://gitlab.mpcdf.mpg.de/dillenburgr/tdp43_lcd_micelles/.

---

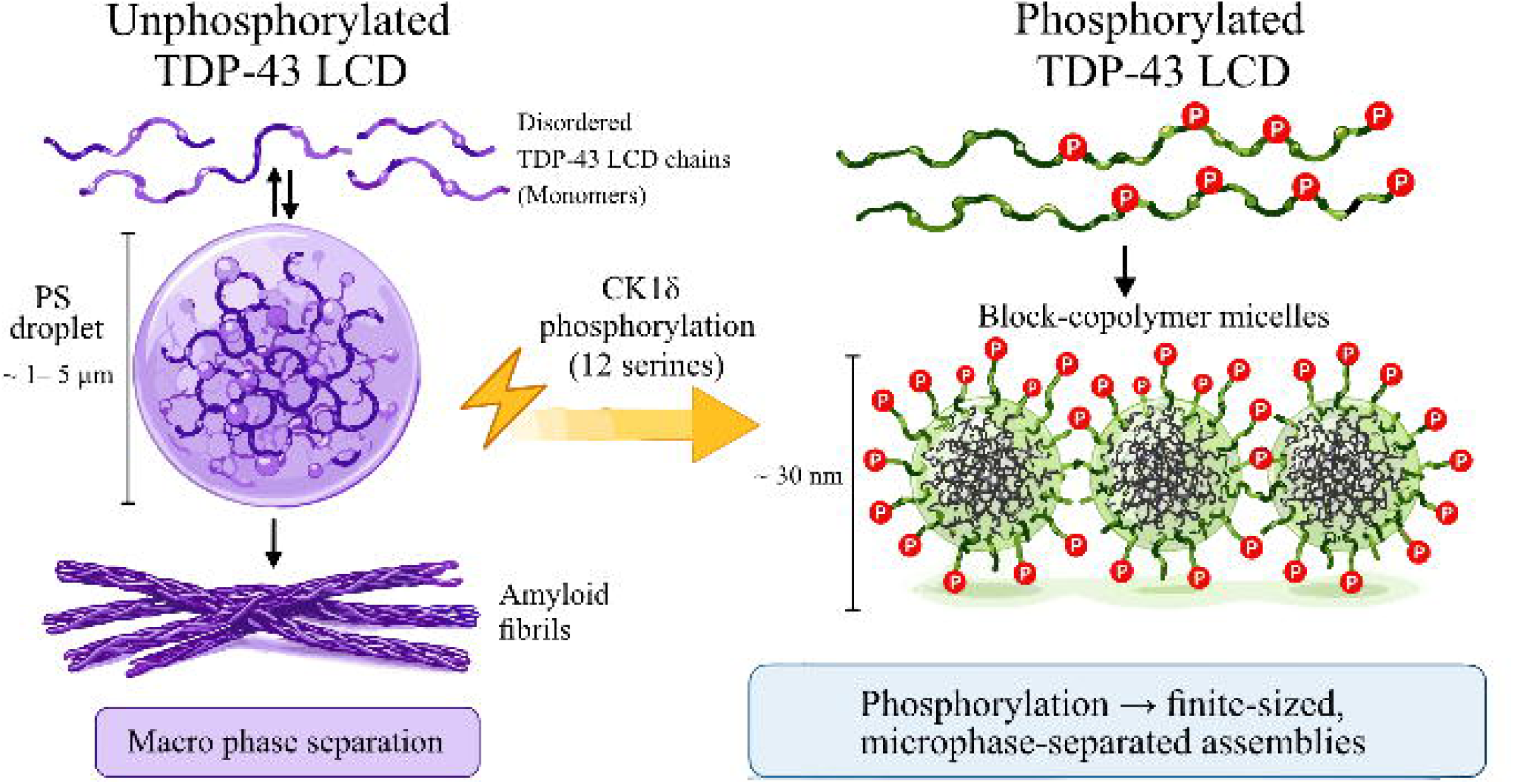
Phosphorylation of TDP-43’s low-complexity domain by CK1δ acts as a molecular switch, redirecting its self-assembly from macroscopic phase separation toward finite-sized, spherical block-copolymer micelles of ~30 nm. The resulting core-corona architecture, with hydrophobic interior and charged phospho-corona, reveals a previously unrecognised regulatory regime for this aggregation-prone protein.

