## Supplementary Information for "“A Phosphorylation-Induced Micellization switch in the low complexity domain of TDP-43”"

A. Lopatina, H. Ruan, T. Scheidt, S. Mosna, E. Pekbilir
Biocentre II, Johannes Gutenberg University, Mainz, Germany

Rodrigo F. Dillenburg

Max Planck Institute for Polymer Research, Mainz, Germany

Biocentre II, Johannes Gutenberg University, Mainz, Germany

Department of Physics, Johannes Gutenberg University Mainz, Germany

Julia Bieber, Carla Schmidt

Biocentre II, Johannes Gutenberg University, Mainz, Germany

Department of Chemistry-Biochemistry

D. Dormann, L. Stelzl, E. A. Lemke
Biocentre II, Johannes Gutenberg University, Mainz, Germany

Institute for Molecular Biology, Mainz, Germany

Frank Schäfer-Depoix

Biocentre I, Johannes Gutenberg University, Mainz, Germany

Electron Microscopy Core Facility (EMCF)

Katharina Landfester*,* Martin Girard*,* Svenja Morsbach

Max Planck Institute for Polymer Research, Mainz, Germany

Friederike Schmid

Department of Physics, Johannes Gutenberg University Mainz, Germany

Martin M. Möckel

Protein Production Core Facility, Institute of Molecular Biology (IMB), Mainz, Germany

Funding: Deutsche Forschungsgemeinschaft (DFG, German Research Foundation)-SFB1551-“Polymer concepts in Cellular Function” Project No. 464588647

Keywords: TDP-43, low-complexity domain, intrinsically disordered proteins, protein phosphorylation, micro phase separation, micellization, worm-like micelles

*#* These authors contributed equally to this work and are co-first authors*: Rodrigo F. Dillenburg, Anastasia Lopatina, Hao Ruan*

| Sequences | (B = phosphoSerine) |
| --- | --- |
| TDP-43 LCD | NRQLERSGRFGGNPGGFGNQGGFGNSRGGGAGLGNNQGSNMGGGMNFGAFSINPAMMAAAQAALQSSWGMMGMLASQQNQSGPSGNNQNQGNMQREPNQAFGSGNNSYSGSNSGAAIGWGSASNAGSGSGFNGGFGSSMSDKSSGWGM |
| TDP-43 LCD 12pS | NRQLERSGRFGGNPGGFGNQGGFGNSRGGGAGLGNNQGSNMGGGMNFGAFSINPAMMAAAQAALQSSWGMMGMLASQQNQSGPSGNNQNQGNMQREPNQAFGSGNN**B**Y**B**GSN**B**GAAIGWG**B**A**B**NAG**B**G**B**GFNGGFG**BB**M**B**DK**BB**GWGM |
| TDP-43 LCD 12D | NRQLERSGRFGGNPGGFGNQGGFGNSRGGGAGLGNNQGSNMGGGMNFGAFSINPAMMAAAQAALQSSWGMMGMLASQQNQSGPSGNNQNQGNMQREPNQAFGSGNN**D**Y**D**GSN**D**GAAIGWG**D**A**D**NAG**D**G**D**GFNGGFG**DD**M**D**DK**DD**GWGM |
| TDP-43 LCD 12DD | NRQLERSGRFGGNPGGFGNQGGFGNSRGGGAGLGNNQGSNMGGGMNFGAFSINPAMMAAAQAALQSSWGMMGMLASQQNQSGPSGNNQNQGNMQREPNQAFGSGNN**DD**Y**DD**GSN**DD**GAAIGWG**DD**A**DD**NAG**DD**G**DD**GFNGGFG**DDDD**M**DD**DK**DDDD**GWGM |
| TDP-43 LCD 12DA | NRQLERSGRFGGNPGGFGNQGGFGNSRGGGAGLGNNQGSNMGGGMNFGAFSINPAMMAAAQAALQSSWGMMGMLASQQNQSGPSGNNQNQGNMQREPNQAFGSGNN**DA**Y**DA**GSN**DA**GAAIGWG**DA**A**DA**NAG**DA**G**DA**GFNGGFG**DADA**MD**DA**K**DADA**GWGM |
| TDP-43 LCD 12DD-148aa | NRQLERSGRFGGNPGGFGNQGGFGNSRGGGAGLGNNQGSNMGGGMNFGAFSINPAMMAAAQAALQSSWGMMGMLASQQNQSGPSGNNQNQGNMQREPNQAFGSGN**DD**Y**DD**SN**DD**AAIGW**DD**A**DD**A**DD**G**DD**FNGGF**DDDDDDDDDD**WGM |

**Supplementary Table 1.** **Sequences used in simulations and experiments in the manuscript.** Phosphoserines are represented by the letter B. Phosphorylation sites and phosphomimetic substitutions are marked in bold.

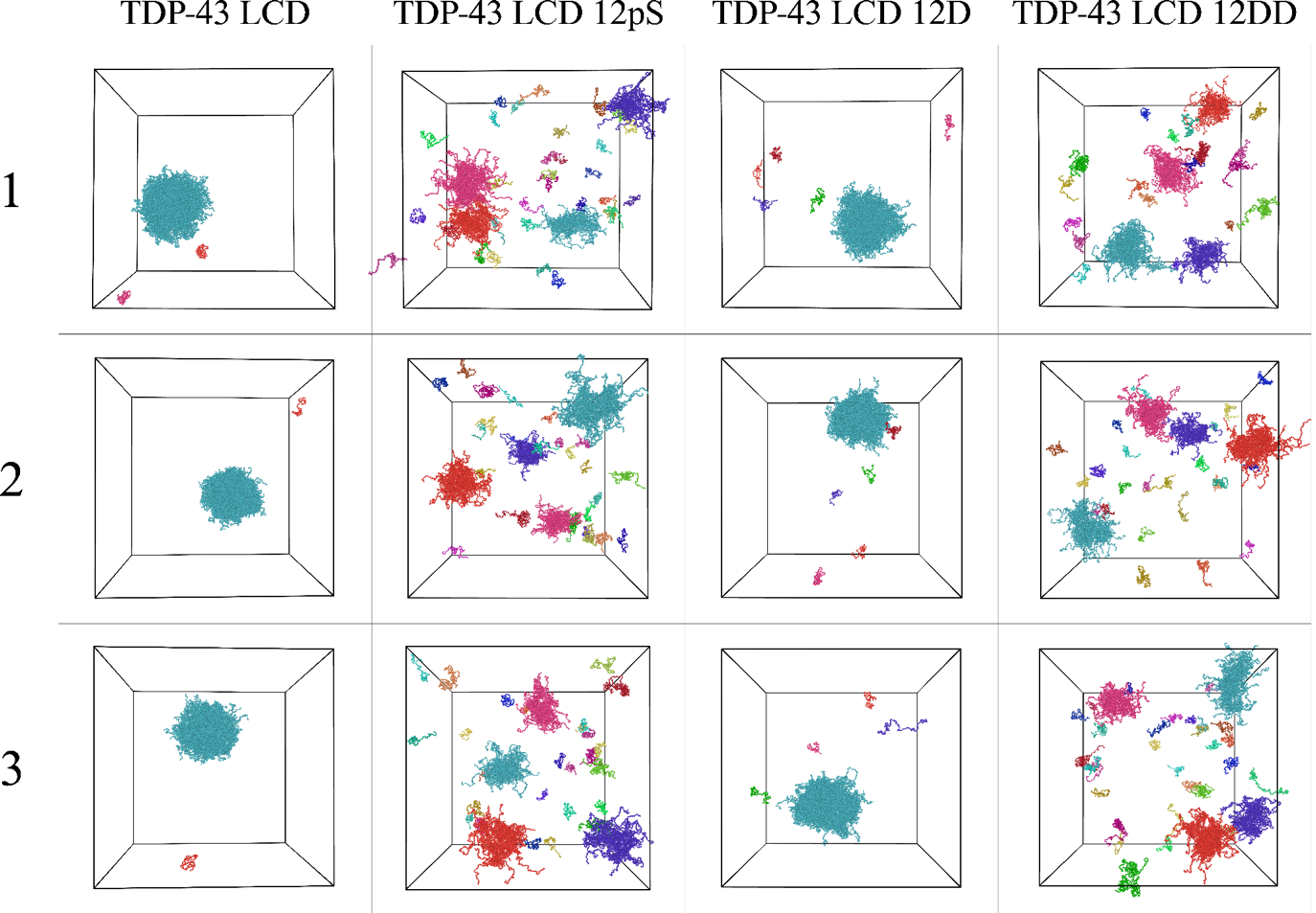

**Supplementary Figure 1. MD simulations yield identical phenotypes as SGCMC.** Snapshots of MD simulations for the 4 simulated sequences. We present one snapshot for each of the 3 replicas, with replica numbers shown on the left. Simulation boxes contain 150 chains each. Images made with OVITO.

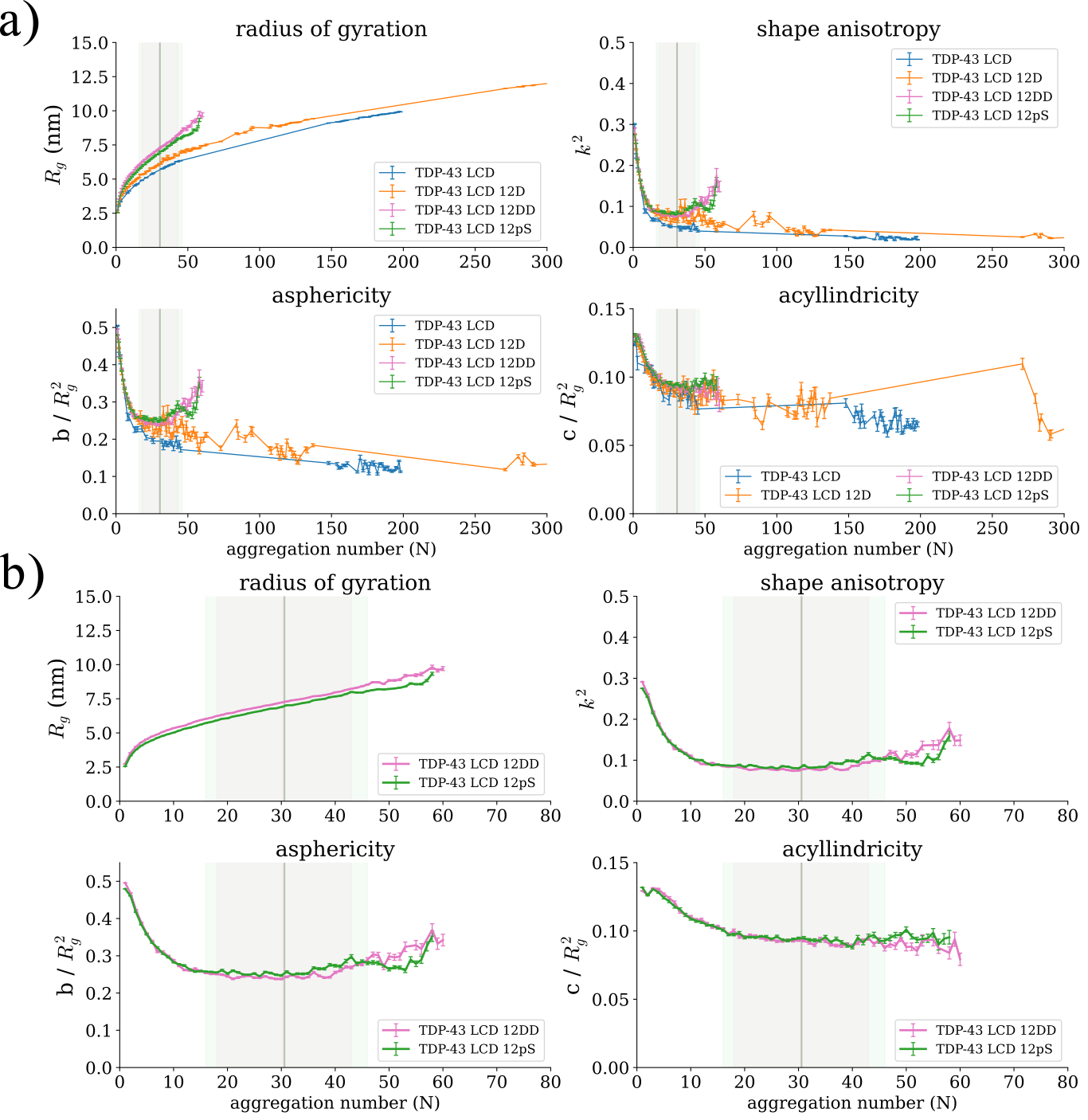

**Supplementary Figure 2. Shape descriptors are consistent with micellization.** Plots of the shape descriptors obtained from the gyration tensor for clusters of different aggregation numbers. Values for **a)** all 4 sequences and **b)** the 2 micellizing sequences (12 DD and 12 pS) zoomed-in. Values are shown as mean $\pm$ the standard error on the mean. The vertical lines represent the mean aggregation number obtained for the fits of micelle size distributions, the shaded region around it represents one standard deviation.

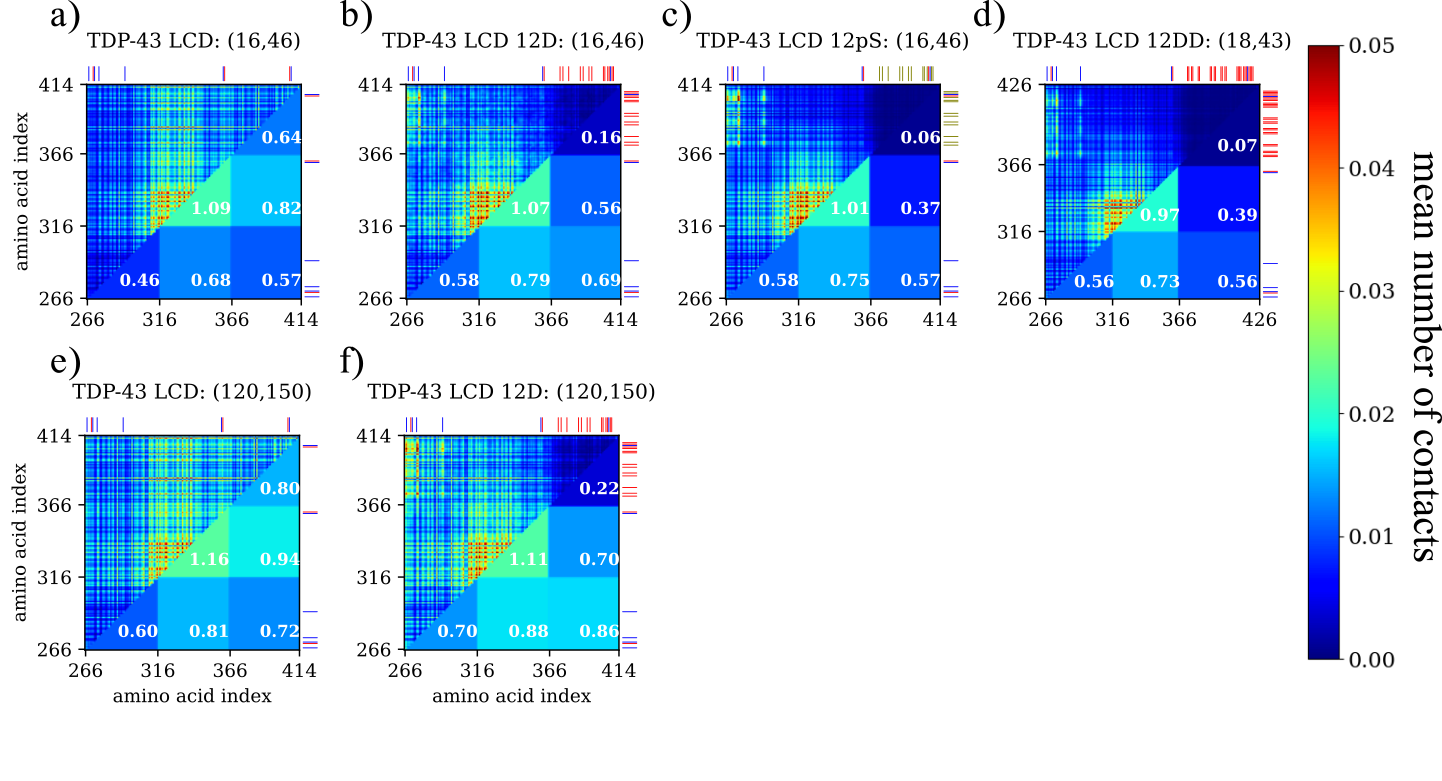

**Supplementary Figure 3. Interchain contact maps highlight the blockiness of phosphorylated TDP-43.** Contact maps for **a)** TDP-43 LCD, **b)**12D, **c)**12pS and **d)**12 DD for micelle-sized clusters and at the most common aggregation numbers for **e)**TDP-43 LCD and **f)**12D. The upper diagonal plot represents the mean number of contacts between each amino acid pair. Lower diagonal terms represent the mean number of contacts per amino acid between 2 blocks. Aggregation number interval is indicated at the top of each contact map. All simulations at 100 mM salt concentration.

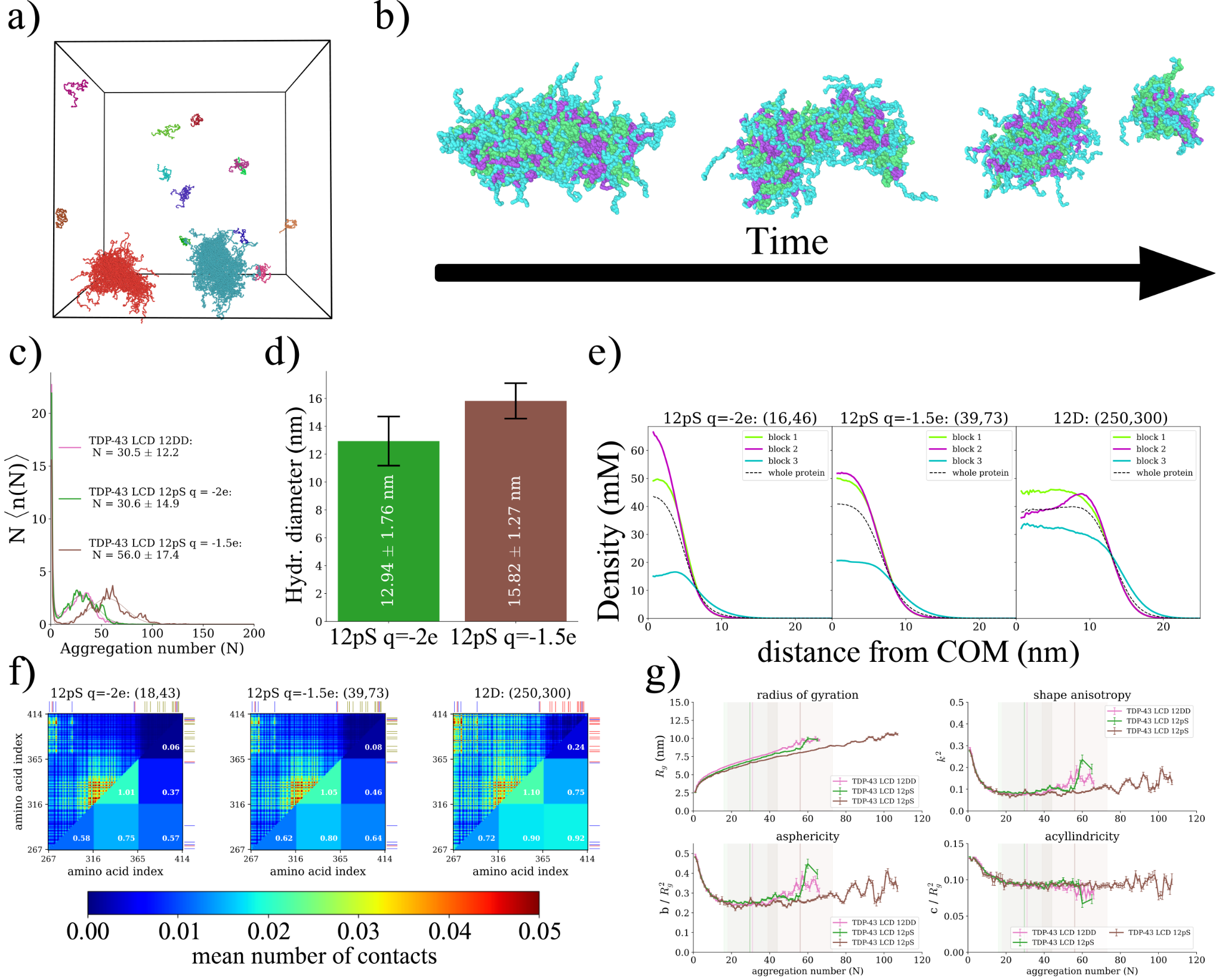

**Supplementary Figure 4. Simulations with phosphoserines with charge -1.5e. a)** Snapshot of simulation box. **b)** Time-lapse of the formation and disassociation of an oblong-shaped micelle resembling a worm-like chain. Image made with OVITO.^[47]^ **c)** Size distribution histograms (full lines) and corresponding fitting curves (dashed lines) for 12 pS at charge -1.5e (brown), 12pS at charge -2.0e (green) and 12 DD (pink). Mean aggregation number and standard deviation are shown in legend. **d)** Mean hydrodynamic radius for 12 pS at charge -1.5e (brown) and 12pS at charge -2.0e (green). Results were obtained from a weighted average of the hydrodynamic radii values of clusters of sizes within one standard deviation of the mean aggregation number of the micelles obtained from the fit to the size distributions. Error bars represent the standard deviation of the distribution of hydrodynamic diameters. **e)** Per-block density profiles at most common aggregation numbers for 12 pS at charge -1.5e, 12pS at charge -2.0e and 12 D mutant. Aggregation number interval is shown on the top between parenthesis. **f)** Interchain contact maps for 12 pS at charge -2.0e, 12pS at charge -1.5e and 12 D. The upper diagonal plot represents the mean number of contacts between each amino acid pair. Lower diagonal terms represent the mean number of contacts per amino acid between 2 blocks. Aggregation number interval is indicated at the top of each contact map. **g)** Shape descriptors obtained from the gyration tensor of individual assemblies as a function of aggregation number for 12 pS at charge -1.5e (brown), 12pS at charge -2.0e (green) and 12 DD (pink). Values are shown as mean $\pm$ the standard error on the mean. The vertical lines represent the mean aggregation number obtained for the fits of micelle size distributions, the shaded region around it represents one standard deviation.

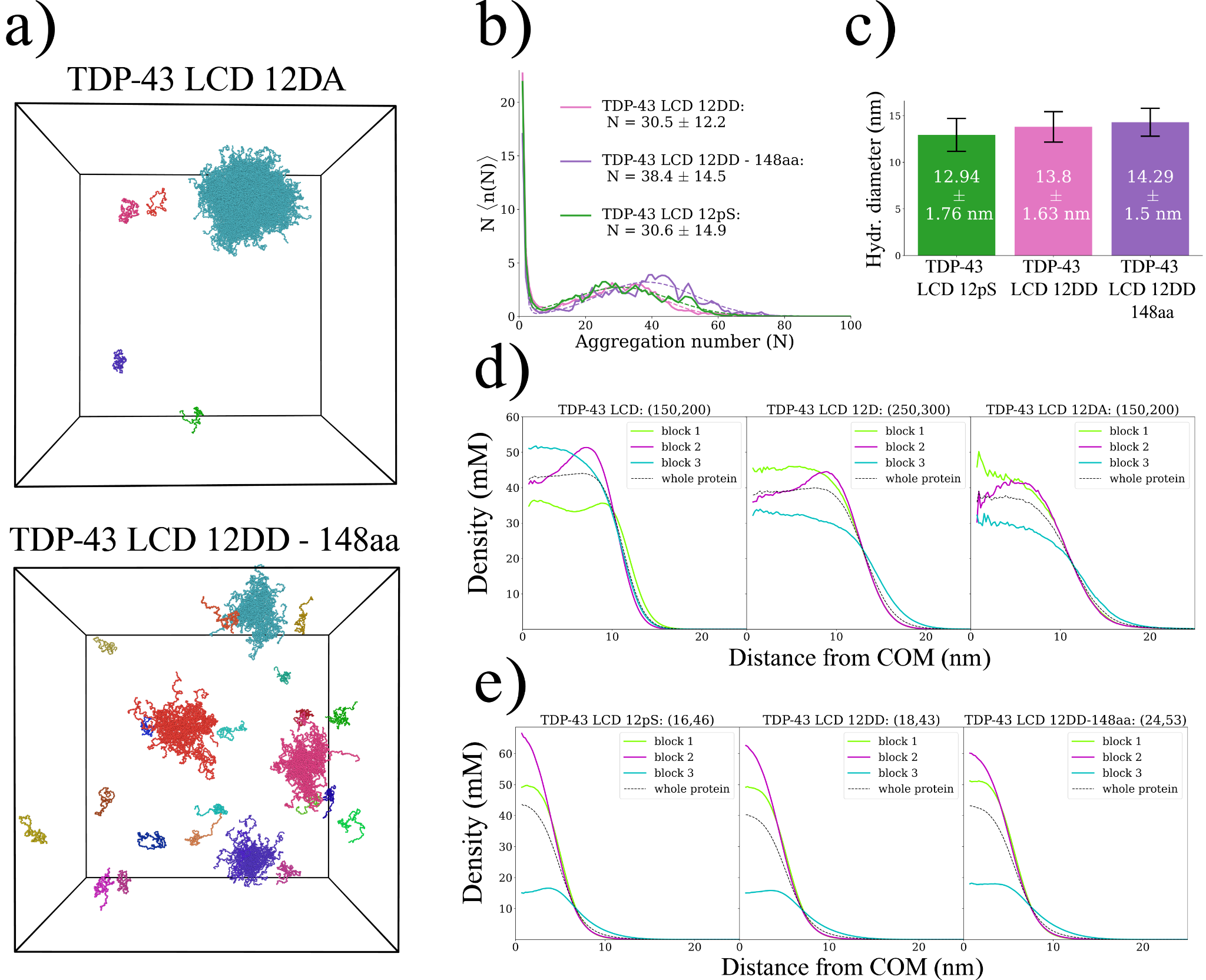

**Supplementary Figure 5: Effect of chain length. a)** Snapshots of simulations for the 12DA and 12DD-148 aa mutants. Made with OVITO. **b)** Size distribution (full lines) and corresponding fitting curves (dashed lines) for the micellizing sequences (TDP-43 LCD 12pS, 12DD and 12DD-148aa). Mean aggregation number and standard deviation obtained from the fit are shown in legend. **c)** Mean hydrodynamic diameter for micellizing sequences. Results were obtained from a weighted average of the hydrodynamic radii values of clusters of sizes within one standard deviation of the mean aggregation number of the micelles obtained from the fit to the size distributions. Error bars represent the standard deviation of the distribution of hydrodynamic diameters. **d)** and **e)** Monomer density profiles per block, measured as a function of the distance from the micelle’s (or cluster’s) centre of mass. Numbers inside parentheses represent the aggregation number interval considered when computing the density profiles. All simulations at 100 mM salt concentration.

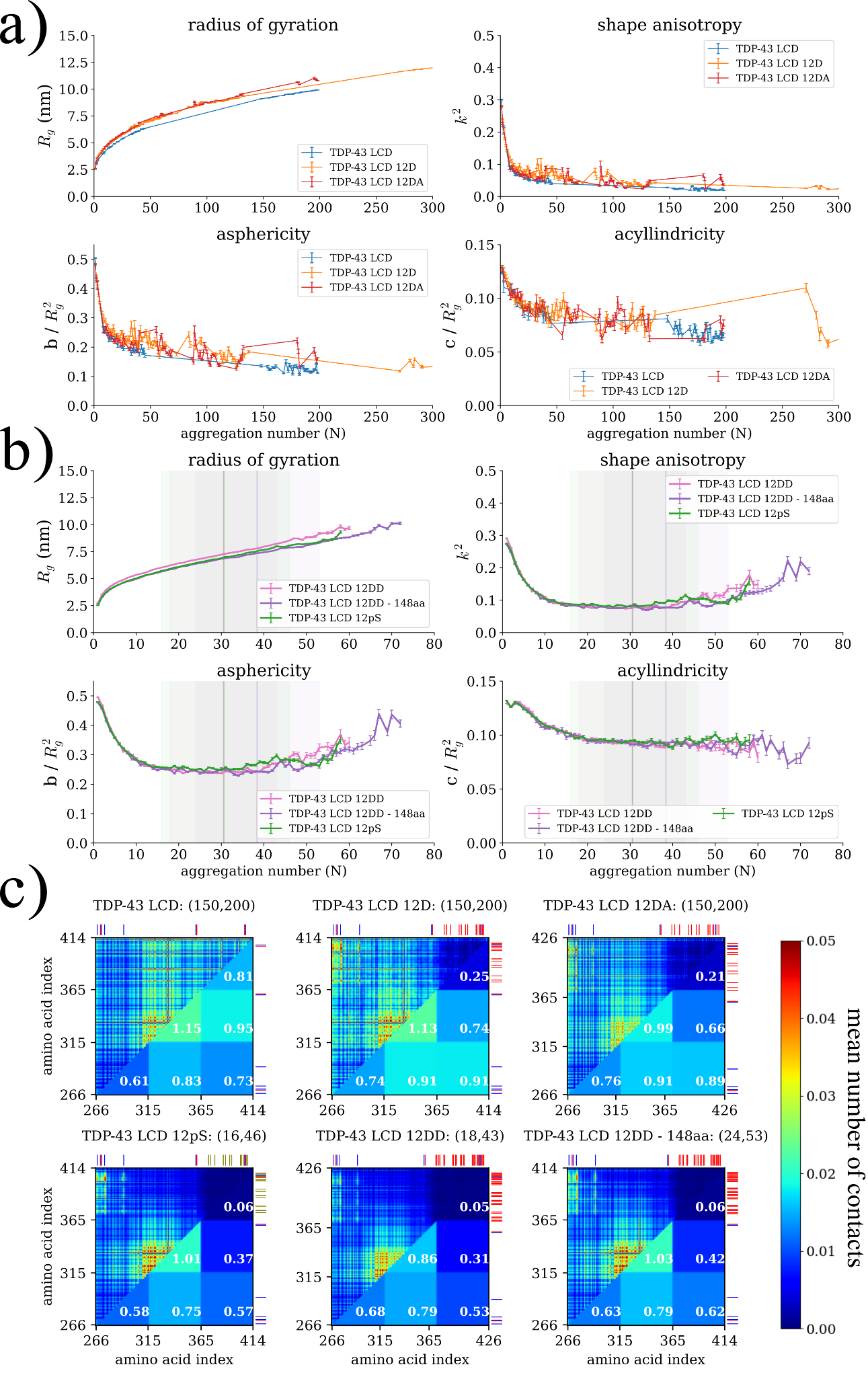

**Supplementary Figure 6. Effect of chain length.** Shape descriptors obtained from the gyration tensor for clusters of different aggregation numbers for the **a)** phase separating and **b)** micellizing sequences. Values are shown as the mean $\pm$ the standard error on the mean. The vertical lines represent the mean aggregation number obtained for the fits of micelle size distributions, the shaded region around it represents one standard deviation. **c)** Interchain contact maps. The upper diagonal plot represents the mean number of contacts between each amino acid pair. Lower diagonal terms represent the mean number of contacts per amino acid between 2 blocks. Aggregation number interval is indicated at the top of each contact map. All simulations at 100 mM salt concentration.

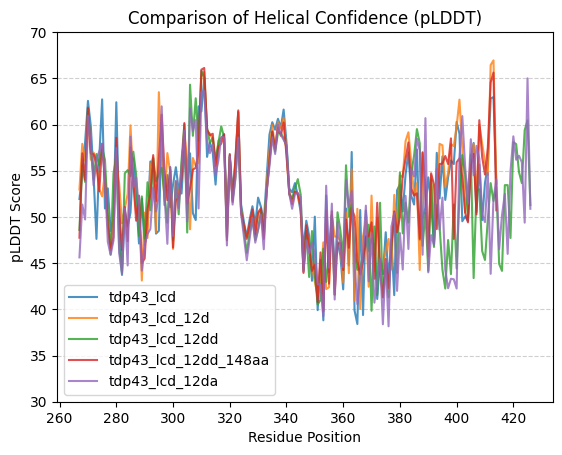

**Supplementary Figure 7. Alpha-fold predicts disordered conformations for all phosphomimetics.** pLDDT scores obtained from alpha-fold predictions indicate that the phosphomimetic substitutions do not induce the formation of secondary or tertiary structure in TDP-43 LCD.

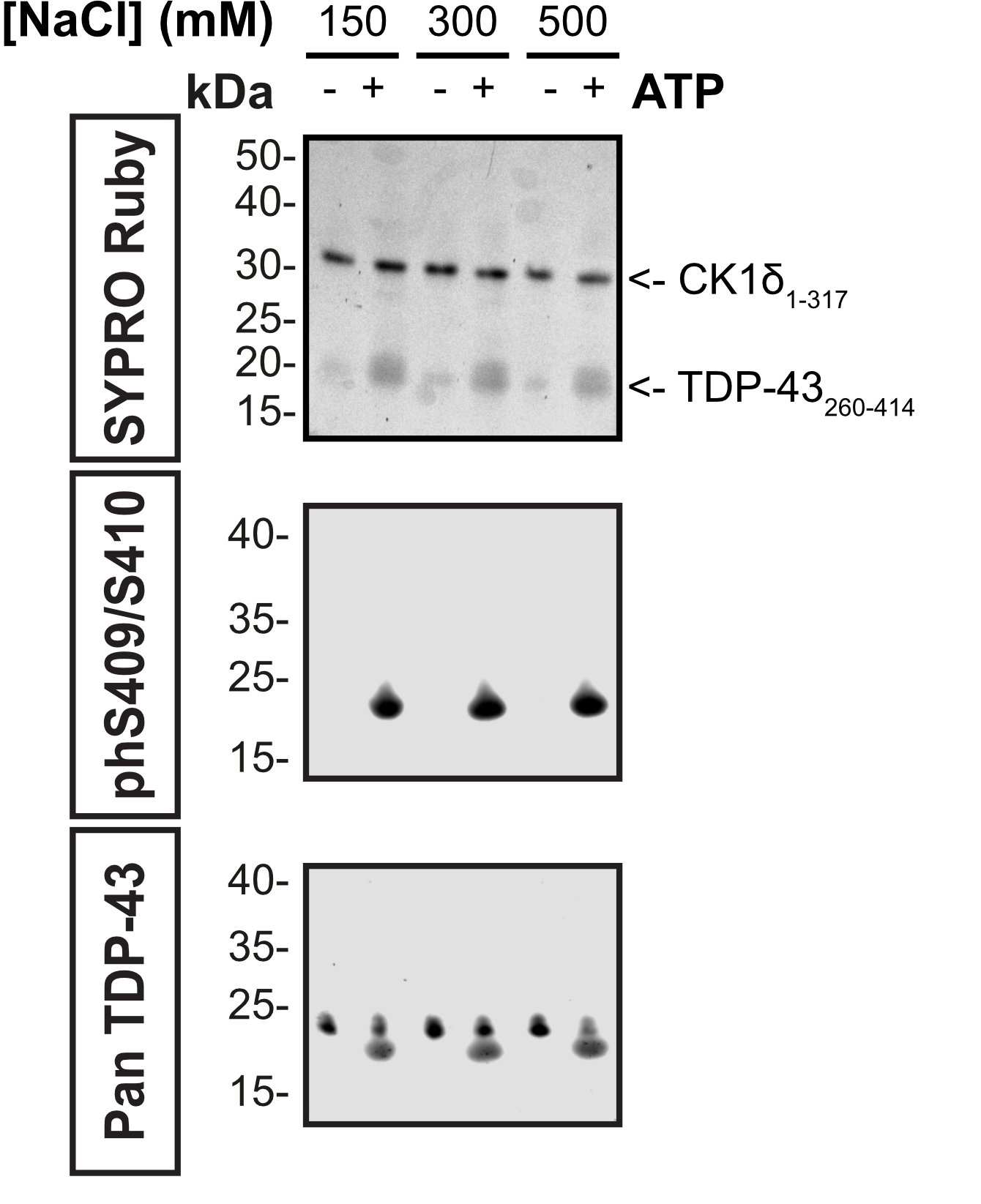

**Supplementary Figure 8. CK1δ-mediated phosphorylation of TDP-43 LCD is maintained across ionic strength conditions. TDP-43 LCD (267-414) was incubated with CK1δ (1-317) in the presence or absence of ATP at 150, 300, and 500 mM NaCl.** Top: SYPRO Ruby total protein stain showing CK1δ (~30 kDa) and TDP-43 LCD (~20 kDa) in all lanes. Middle: Anti-phS409/S410 immunoblot detecting phosphorylated TDP-43 LCD exclusively in ATP-containing reactions (+) at all three ionic strengths, confirming that CK1δ activity is not impaired by elevated salt concentrations. Bottom: Pan TDP-43 antibody confirming equal loading of TDP-43 LCD across all conditions. Together, these data demonstrate that the phosphorylation state of the LCD is equivalent at 150, 300, and 500 mM NaCl, and that differences in assembly behaviour observed across ionic strengths are attributable to electrostatic screening rather than to changes in the phosphorylation pattern.

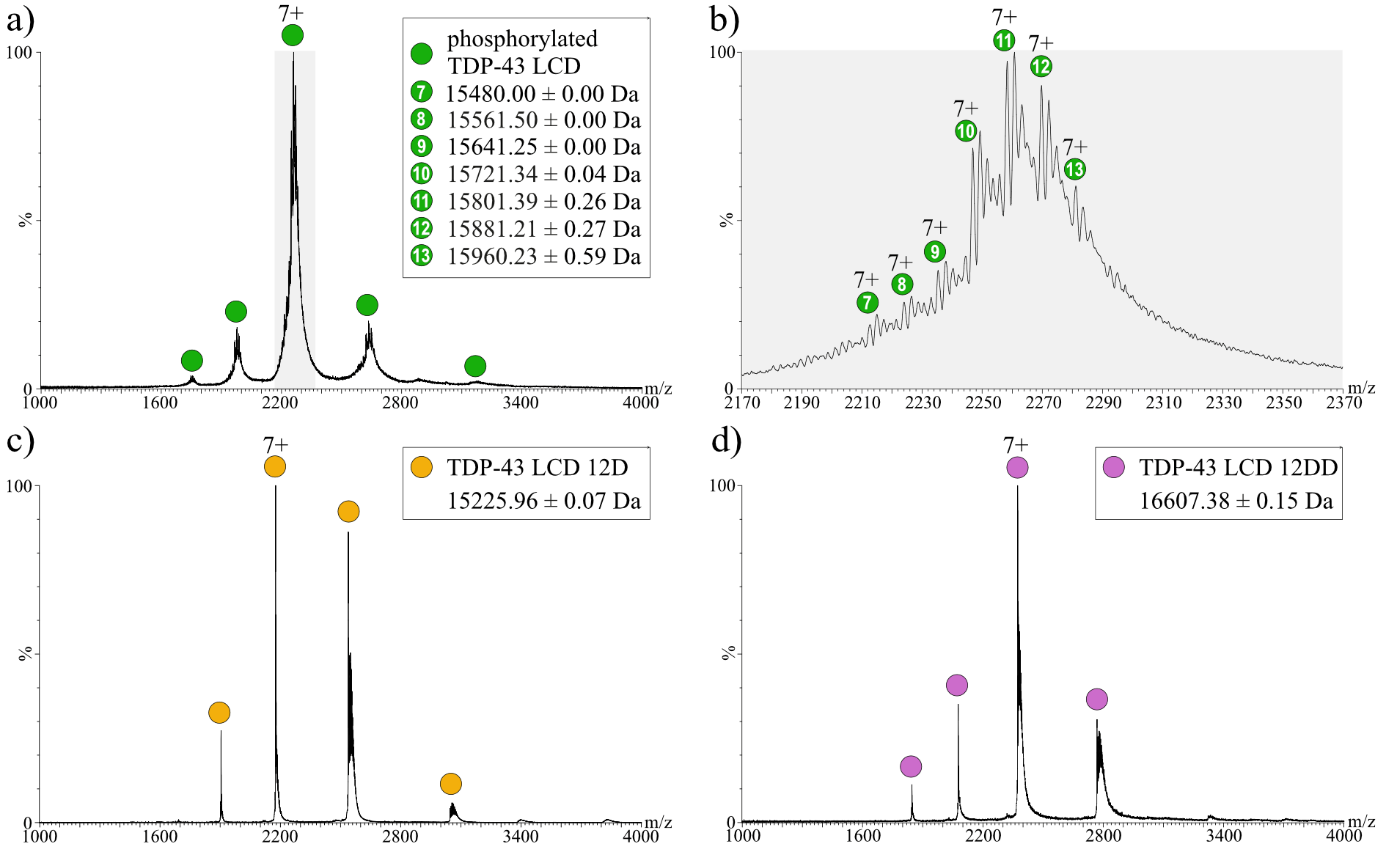

**Supplementary Figure 9. Native mass spectrometry of phosphorylated TDP-43 LCD and TDP-43 LCD phosphomimetic mutants. a)** Mass spectrum of in vitro phosphorylated TDP-43 LCD. Charge state distributions corresponding to a range of TDP-43 LCD populations differing in the phosphorylation stoichiometry were observed. The masses obtained from these charge state distributions agree with seven to thirteen phosphorylated serine residues. **b)** Magnification of the 7+ charge states of phosphorylated TDP-43 LCD (highlighted in grey in panel a). Several populations of phosphorylated TDP-43 LCD varying in the phosphorylation stoichiometry are assigned. Numbers in circles correspond to the number of phosphorylation sites of the specific population. Additional signals of each population indicate sodium adducts. **c)** Mass spectrum of the 12D TDP-43 LCD phosphomimetic mutant. The corresponding charge state distribution is assigned. The obtained molecular mass agrees with the theoretical mass of the mutant (15226.06 Da). Sodium adducts are observed for all charge states. **d)** Mass spectrum of the 12DD TDP-43 LCD phosphomimetic mutant. The corresponding charge state distribution is assigned. The obtained molecular mass agrees with the theoretical mass of the mutant (16607.12 Da). Sodium adducts are observed for all charge states.

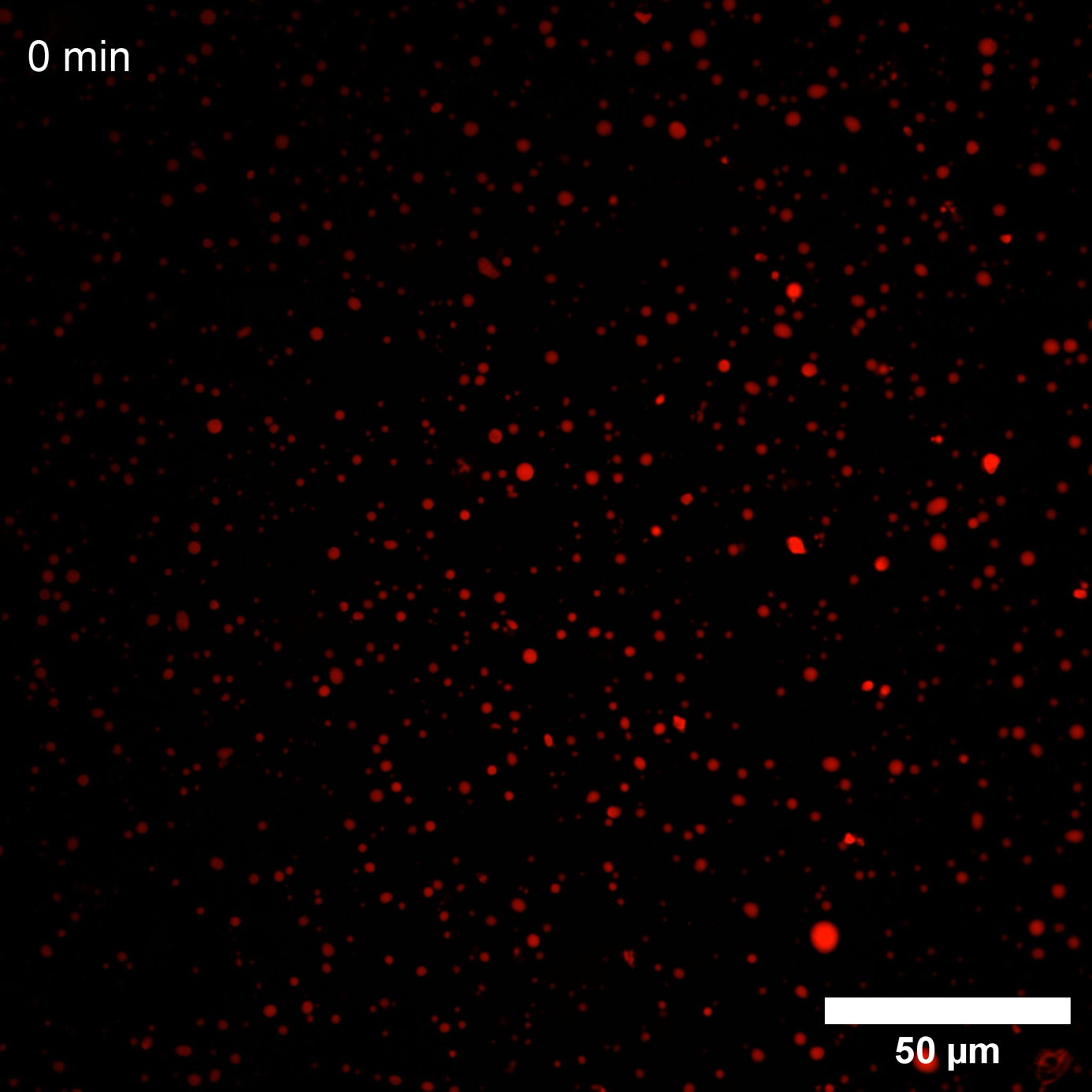

**Supplementary Movie S1.** Time-lapse fluorescence microscopy of unphosphorylated TDP-43 LCD (50 μM total protein doped with 10 nM LD655-labeled A366C TDP-43 LCD) at 150 mM NaCl in a Pluronic-passivated ibidi µ-Slide chamber, acquired at 60-s intervals over 60 min. The droplet population progressively coalesces into fewer, larger droplets over the observation window. Scale bar, 50 µm. Playback speed, 5 fps.

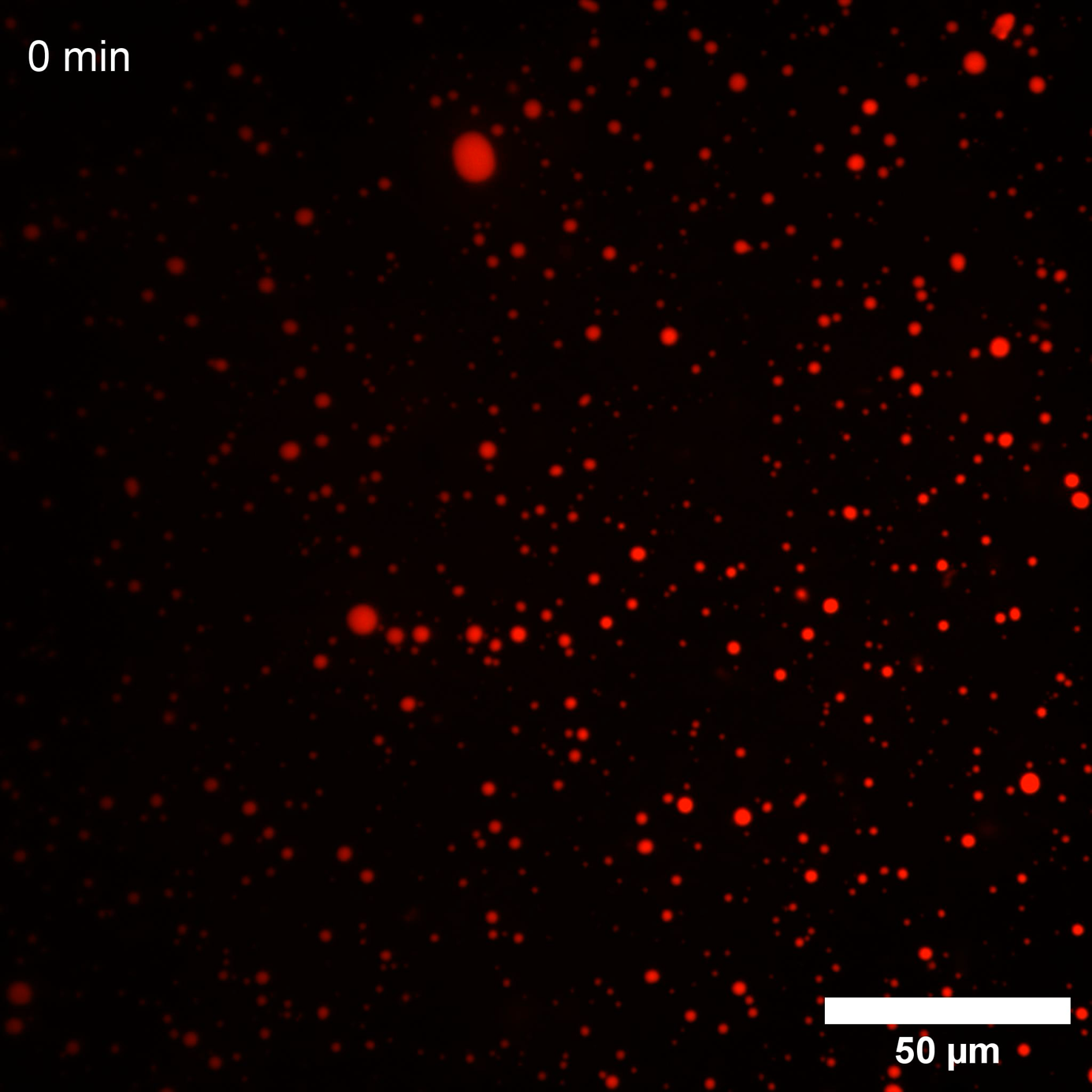
**Supplementary Movie S2.** Time-lapse fluorescence microscopy of unphosphorylated TDP-43 LCD (50 μM total protein doped with 10 nM LD655-labeled A366C TDP-43 LCD) at 300 mM NaCl in a Pluronic-passivated ibidi µ-Slide chamber, acquired at 60-s intervals over 60 min. The droplet population progressively coalesces into fewer, larger droplets over the observation window. Scale bar, 50 µm. Playback speed, 5 fps.

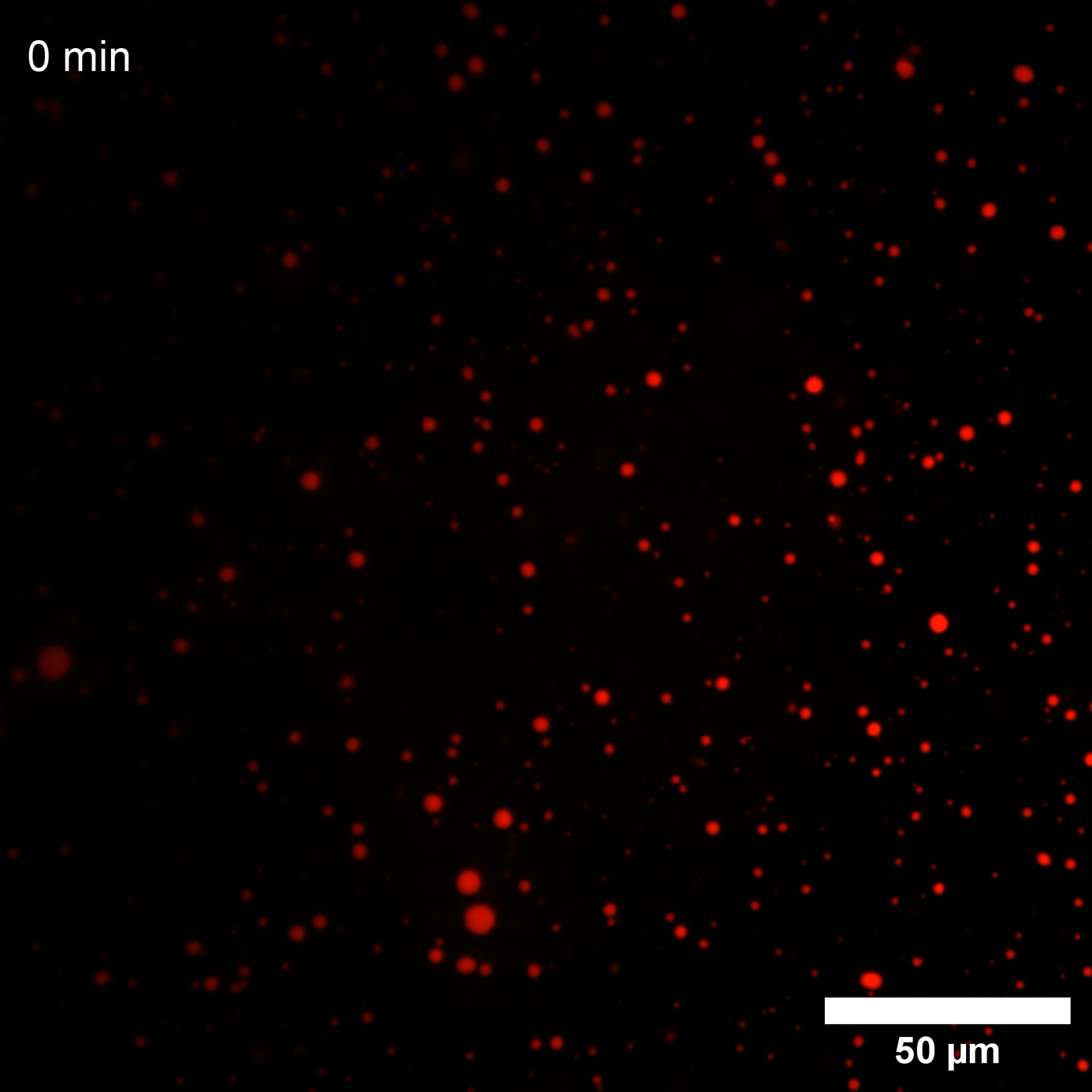
**Supplementary Movie S3.** Time-lapse fluorescence microscopy of unphosphorylated TDP-43 LCD (50 μM total protein doped with 10 nM LD655-labeled A366C TDP-43 LCD) at 500 mM NaCl in a Pluronic-passivated ibidi µ-Slide chamber, acquired at 60-s intervals over 60 min. The droplet population progressively coalesces into fewer, larger droplets over the observation window. Scale bar, 50 µm. Playback speed, 5 fps.

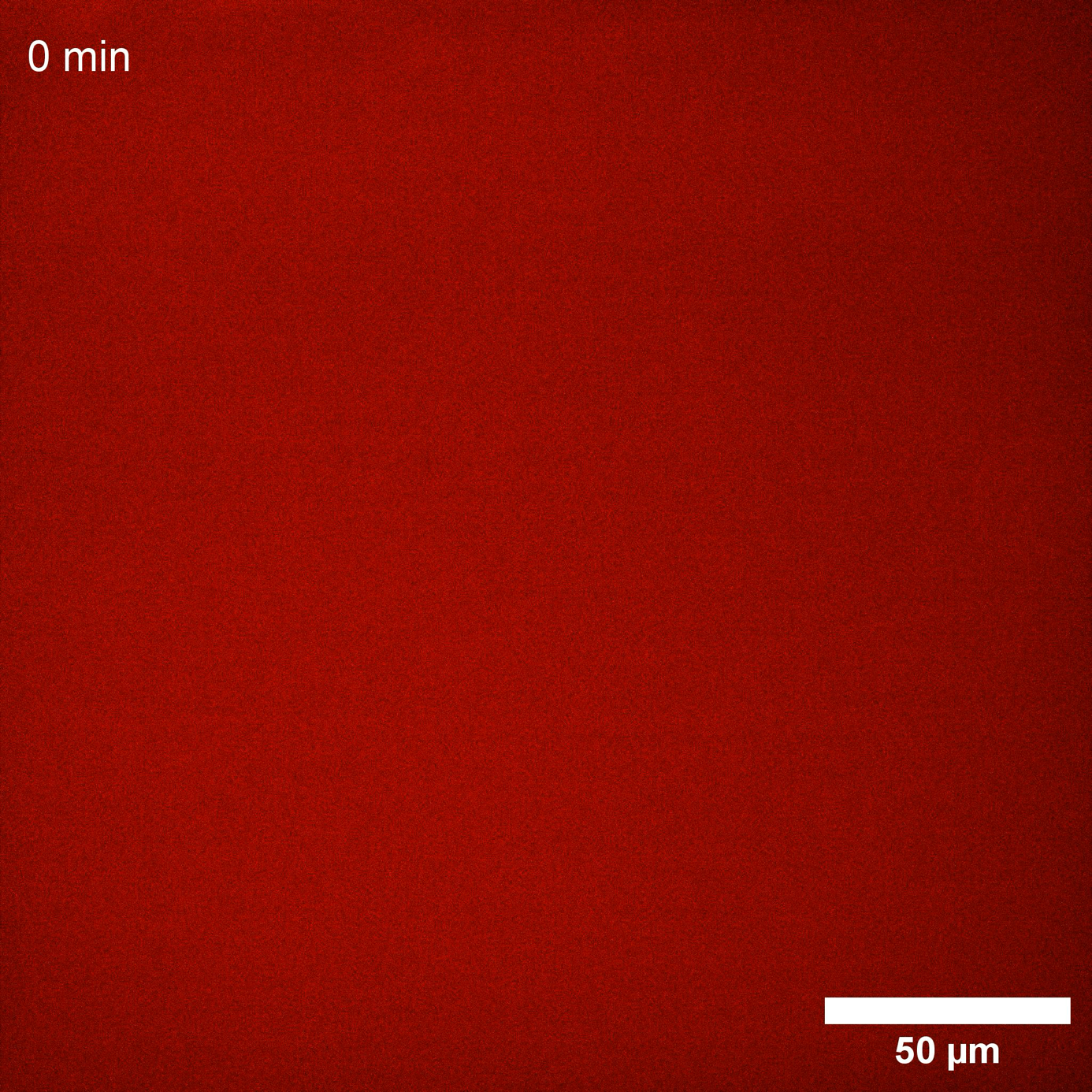
**Supplementary Movie S4.** Time-lapse fluorescence microscopy of CK1δ-phosphorylated TDP-43 LCD (50 μM total protein doped with 10 nM LD655-labeled A366C TDP-43 LCD) at 150 mM NaCl in a Pluronic-passivated ibidi µ-Slide chamber, acquired at 60-s intervals over 60 min following a 1-h in-well incubation. No droplets are detected throughout the observation window. Scale bar, 50 µm. Playback speed, 5 fps.

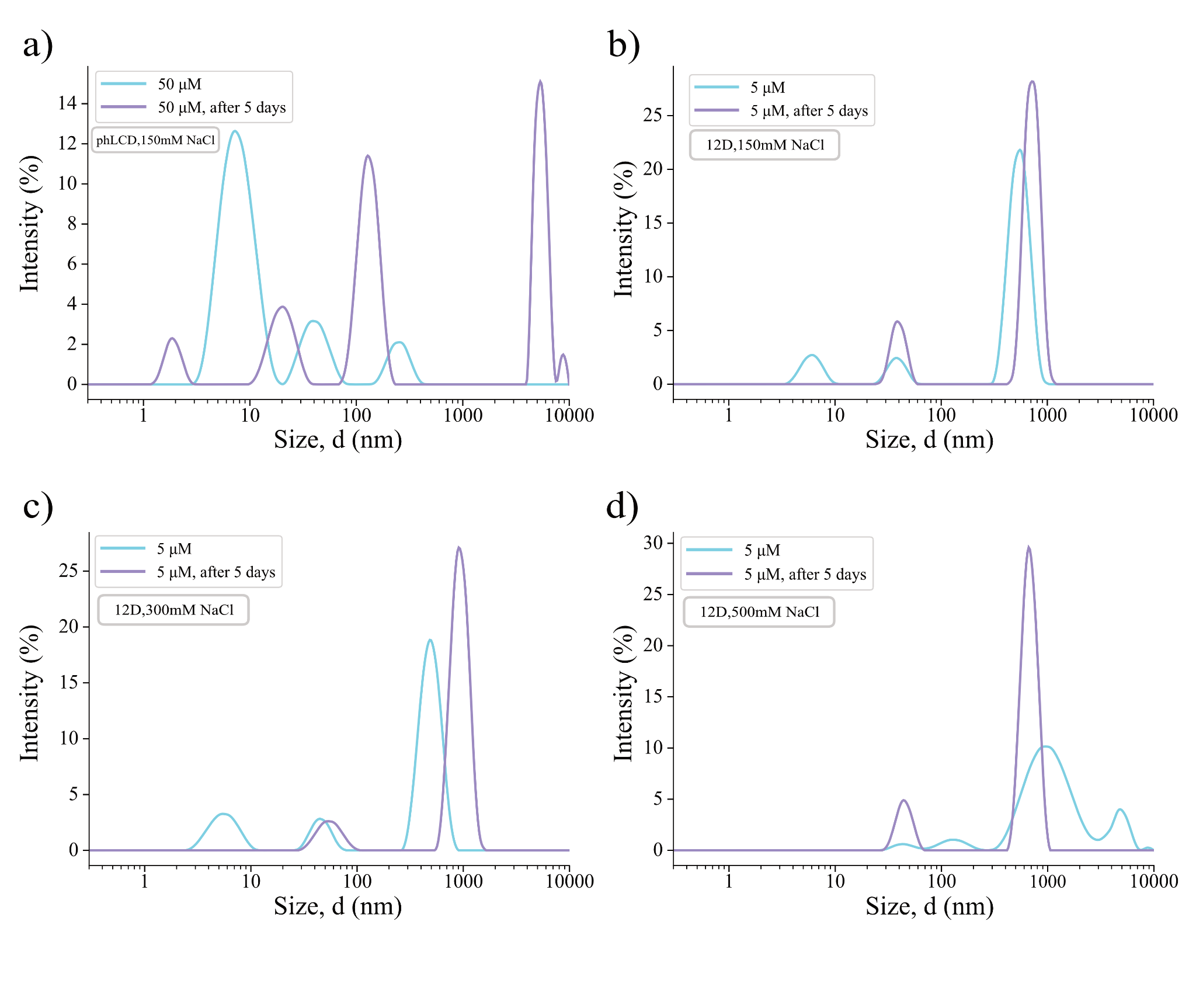

**Supplementary Figure 10. Time-dependent DLS size distributions of phosphorylated TDP-43 LCD and 12D at increasing ionic strength. a)** Intensity-weighted hydrodynamic diameter distributions of 50 µM phLCD in 150 mM NaCl buffer, measured immediately after sample preparation (blue) and again after 5 days of incubation at room temperature (purple). The ~30-60 nm micellar population present above the CMC remains the dominant species after 5 days, indicating that the spherical phLCD micelles are temporally stable on multi-day timescales. Large species >100 nm detected in both protein samples and buffer controls are attributed to background scattering and excluded from analysis. **b-d)** Intensity-weighted hydrodynamic diameter distributions of 5 µM 12D phosphomimetic at increasing ionic strength - **b)** 150 mM NaCl, **c)** 300 mM NaCl, and **d)** 500 mM NaCl - measured after sample preparation (blue) and after 5 days of incubation (purple). At all three salt concentrations, the size distributions remain qualitatively similar after 5 days, confirming that the 12D assemblies are temporally stable across the full ionic-strength range tested and demonstrating that the morphological transitions observed at elevated salt (Section 2.4) reflect equilibrium states.

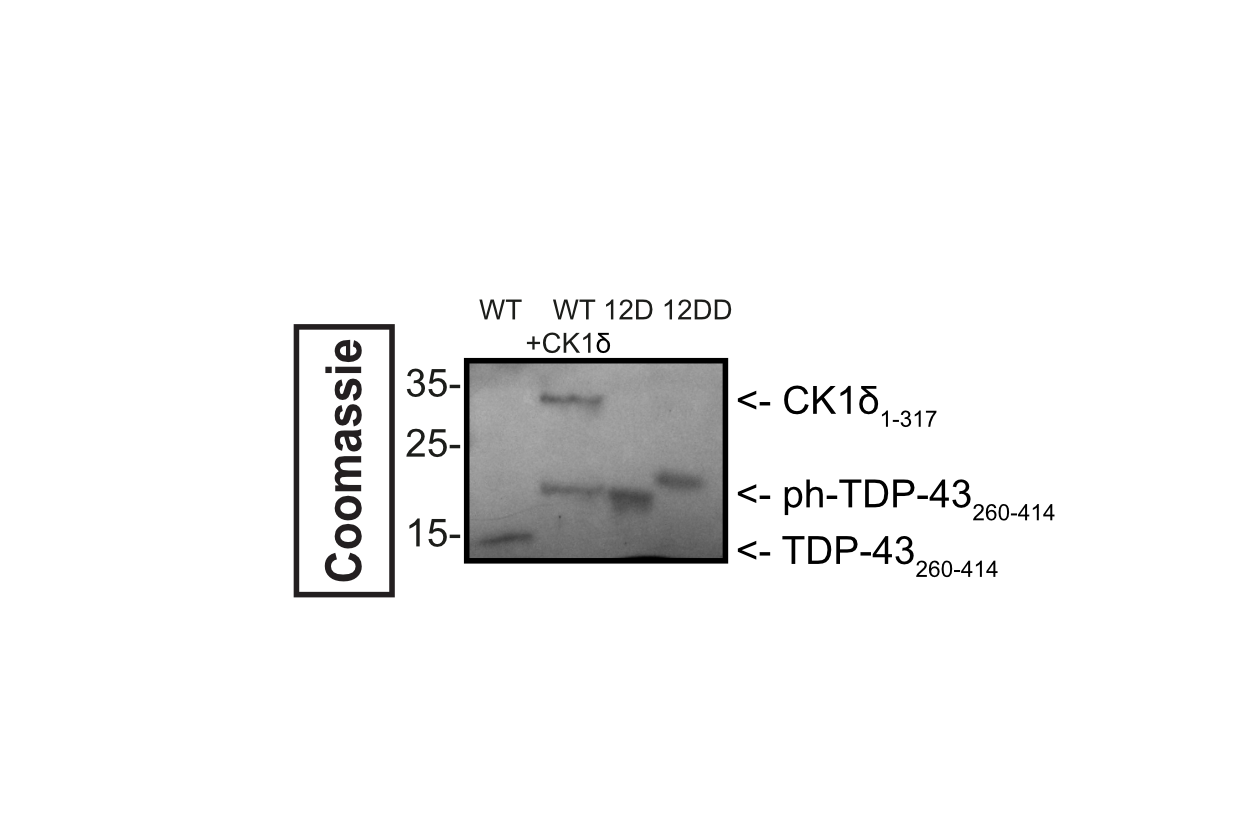
**Supplementary Figure 11. SDS-PAGE resolves CK1δ-phosphorylated TDP-43 LCD and the 12D and 12DD phosphomimetic variants as electrophoretically distinct species.** Coomassie-stained SDS-PAGE of recombinant TDP-43 LCD (residues 267-414): unphosphorylated (WT), CK1δ-treated (WT + CK1δ; 1:10 kinase-to-protein, 0.75 mM ATP), and the phosphomimetic variants 12D and 12DD, resolved on a 4-12% Bis-Tris gel in 1×MES buffer. Molecular-weight markers (kDa) are indicated on the left. WT migrates as a sharp band close to 15 kDa, consistent with its formula mass (~14.8 kDa); upon CK1δ treatment, it shifts to ~16-18 kDa (CK1δ_1-317_ visible at ~27-30 kDa). The phLCD band migrates between the 12D (~16 kDa) and 12DD (~19 kDa) bands. Under these denaturing conditions, electrophoretic mobility is dominated by bound SDS and, for the 12-residue-longer 12DD construct, additionally by chain length; the gel therefore serves only to illustrate that phLCD, 12D and 12DD are electrophoretically distinct species and is not used to quantify the phosphorylation-induced charge, which is established by native mass spectrometry (Figure S9).

| **Construct** | **NaCl** | **Method** | **CMC_1_ (µM)** | **CMC_2_ (µM)** | **Figure** | **Notes** |
| --- | --- | --- | --- | --- | --- | --- |
| phLCD | 150 mM | Pyrene | 1.38 | __ | 4c | R^2^ = 0.93 |
| phLCD | 300 mM | Pyrene | 1.60 | __ | 8a | R^2^ = 0.85 |
| phLCD | 500 mM | Pyrene | 1.45 | __ | 8b | R^2^ = 0.91 |
| 12D | 150 mM | Pyrene | 1.50 | __ | 7a | R^2^ = 0.99 |
| 12D | 300 mM | Nile Red | 1.32 | 5.67 | 9a (top) | Two-step |
| 12D | 500 mM | Nile Red | 0.48 | 3.28 | 9a (bot) | Two-step |
| 12DD | 150 mM | Pyrene | 1.93 | __ | 7b | R^2^ = 0.93 |
| 12DD | 300 mM | Pyrene | 0.52 | __ | 10a | R^2^ = 0.82 |
| 12DD | 500 mM | Pyrene | 0.89 | __ | 10b | R^2^ = 0.89 |

**Supplementary Table 2.** **Summary of critical micelle concentrations (CMCs) for TDP-43 LCD constructs under all tested conditions.** CMCs were determined by fluorescence-based assays as described in Methods Sections 4.12.1 (pyrene) and 4.12.2 (Nile Red). For the 12D construct at elevated ionic strength, two CMC values are reported (CMC_1_ and CMC_2_), corresponding to the onset of spherical micelle formation and the transition to worm-like micelles, respectively. Buffer compositions: phLCD - 50 mM HEPES pH 7.5, 10 mM MgCl_2_, 0.75 mM ATP, 1 mM DTT, (150 mM/300 mM/500 mM NaCl); 12D and 12DD - 20 mM Tris pH 7.5, (150 mM/300 mM/500 mM NaCl). All measurements performed at 25 °C. R^2^ values correspond to the logistic fit shown in the respective figure (provided for visual guidance only; CMC values were extracted exclusively by the tangent-intersection method).

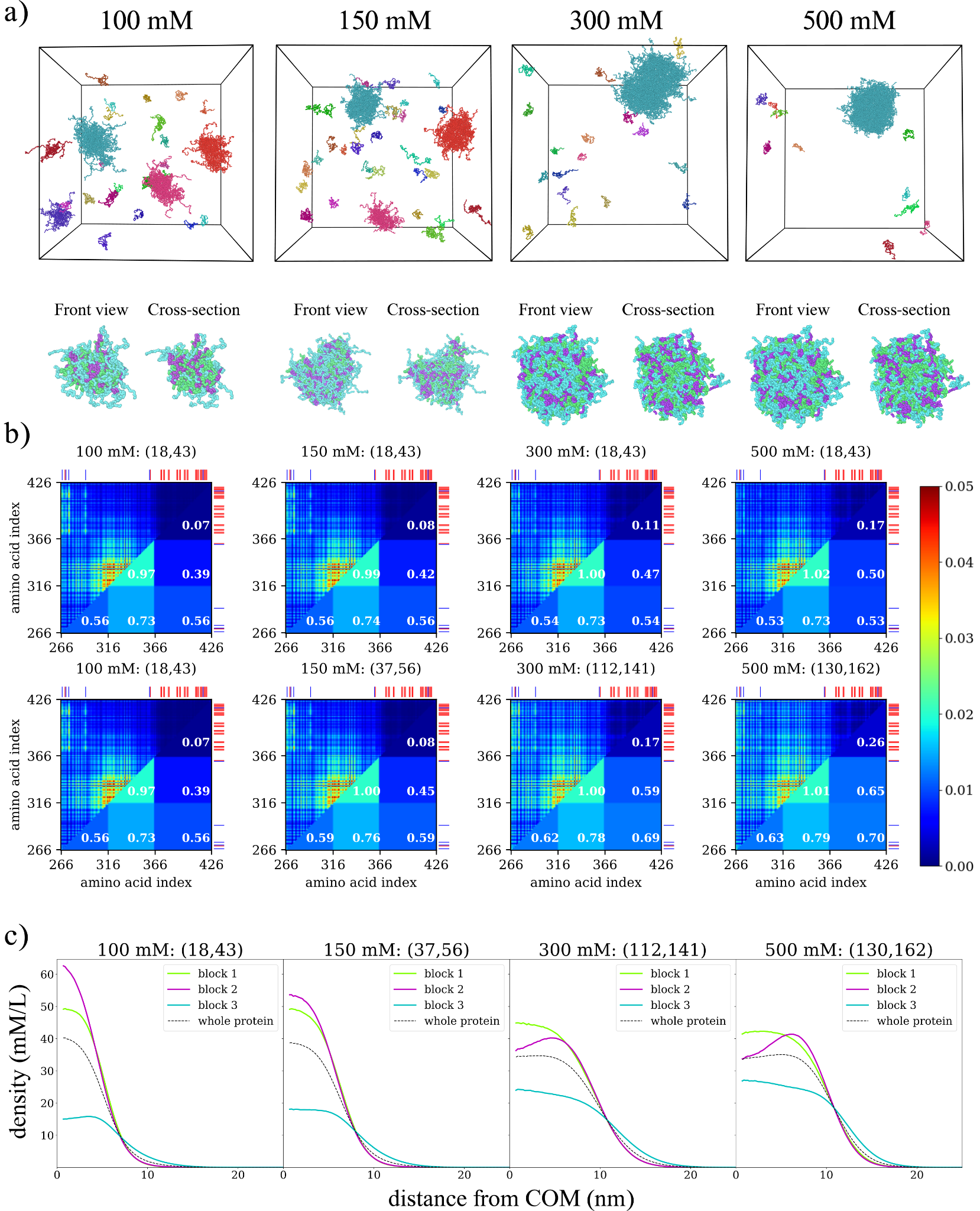

**Supplementary Figure 12. Effect of salt concentration on micellization of the 12 DD mutant. a)** (top) snapshot of simulation box for the 4 salt concentrations, (bottom) snapshot of one micelle as well as a cross-section view of it revealing its internal architecture. Different colors represent the 3 blocks of TDP-43 LCD. Images made with OVITO. **b)** Interchain contact maps at (top) characteristic micelle aggregation number interval for micelles at 100 mM salt concentration and (bottom) most common values of aggregation number observed. The upper diagonal plot represents the mean number of contacts between each amino acid pair. Lower diagonal terms represent the mean number of contacts per amino acid between 2 blocks. Aggregation number interval is indicated at the top of each contact map. **c)** Monomer density profiles per block, measured as a function of the distance from the micelle’s (or cluster’s) centre of mass. Numbers inside parentheses represent the aggregation number interval considered when computing the density profiles.

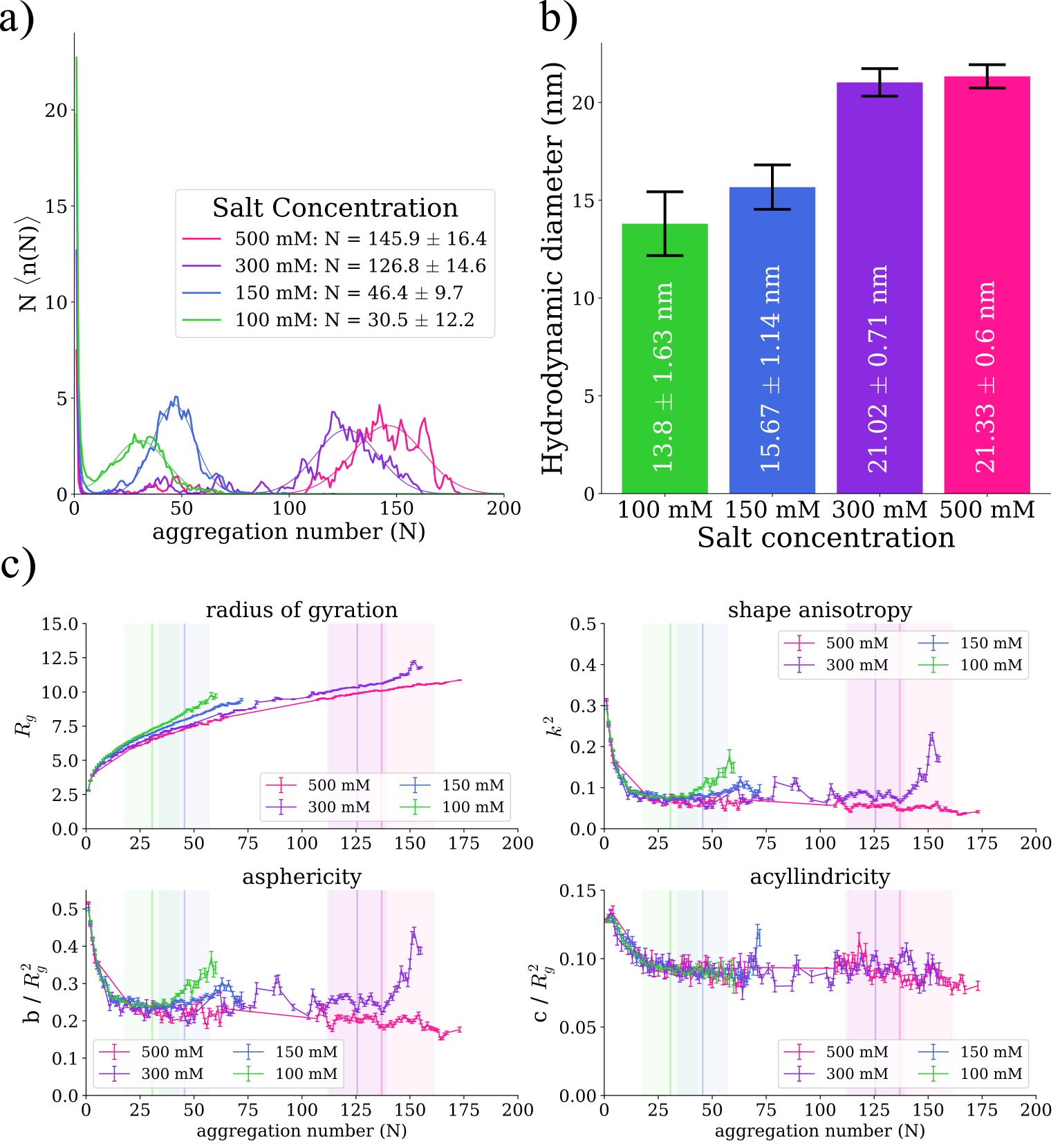

**Supplementary Figure 13. Effect of salt concentration on micellization of the 12 DD mutant. a)** Size distribution (full lines) and corresponding fitting curves (dashed lines) for different salt concentrations. Mean aggregation number and standard deviation obtained from the fit are shown in legend. **b)** Mean hydrodynamic diameter at different salt concentrations. Results were obtained from a weighted average of the hydrodynamic radii values of clusters of sizes within one standard deviation of the mean aggregation number of the micelles obtained from the fit to the size distributions. Error bars represent the standard deviation of the distribution of hydrodynamic diameters. **c)** Shape descriptors obtained from the gyration tensor of individual assemblies as a function of aggregation number at different salt concentrations. Values are shown as the mean $\pm$ the standard error on the mean. The vertical lines represent the mean aggregation number obtained for the fits of micelle size distributions, the shaded region around it represents one standard deviation.

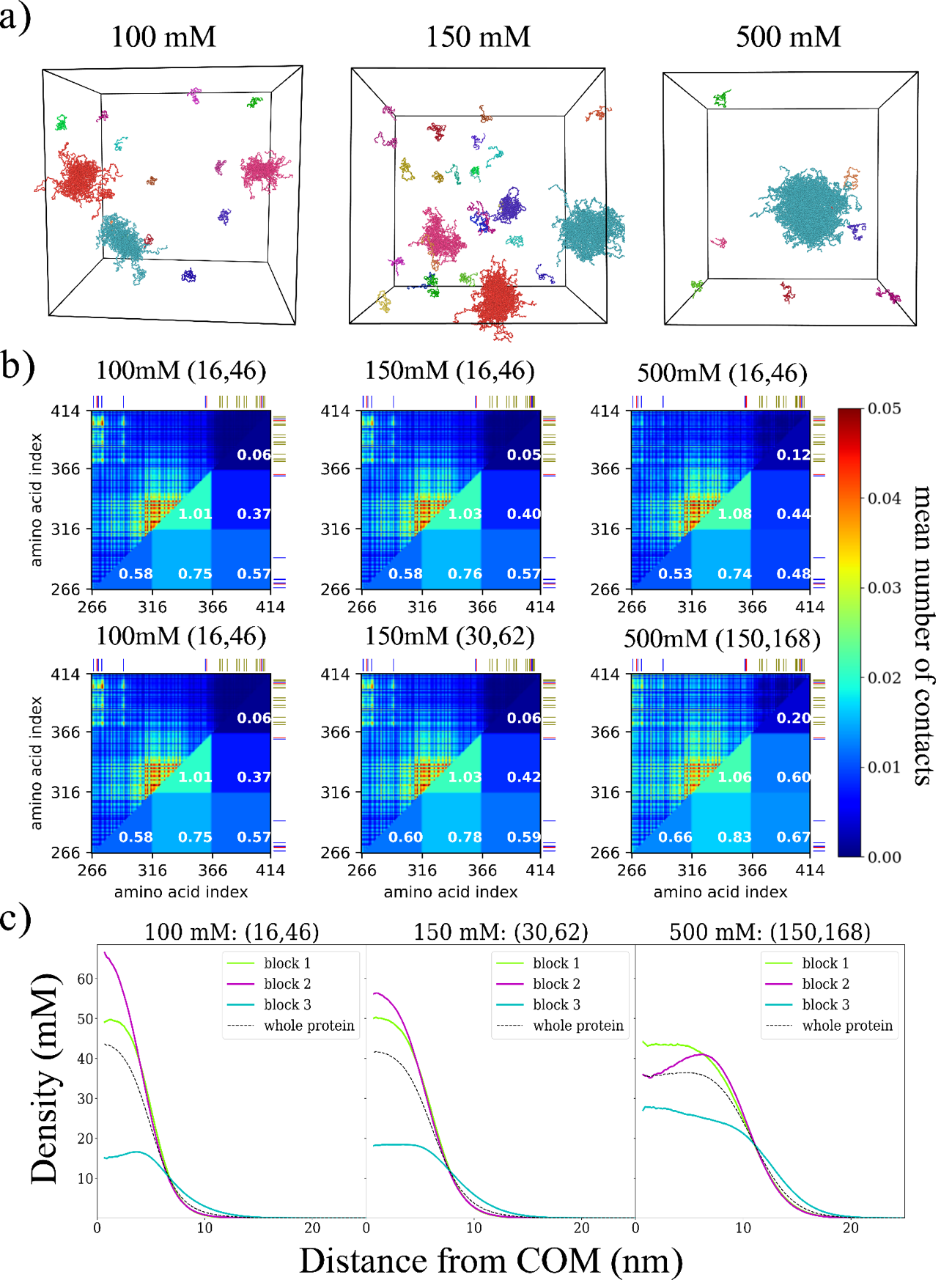

**Supplementary Figure 14. Effect of salt concentration on micellization of TDP-43 LCD 12 pS. a)** snapshot of simulation box for the 3 salt concentrations. Images made with OVITO. **b)** Interchain contact maps at (top) characteristic micelle aggregation number interval for micelles at 100 mM salt concentration and (bottom) most common values of aggregation number observed. The upper diagonal plot represents the mean number of contacts between each amino acid pair. Lower diagonal terms represent the mean number of contacts per amino acid between 2 blocks. Aggregation number interval is indicated at the top of each contact map. **c)** Monomer density profiles per block, measured as a function of the distance from the micelle’s (or cluster’s) centre of mass. Numbers inside parentheses represent the aggregation number interval considered when computing the density profiles.

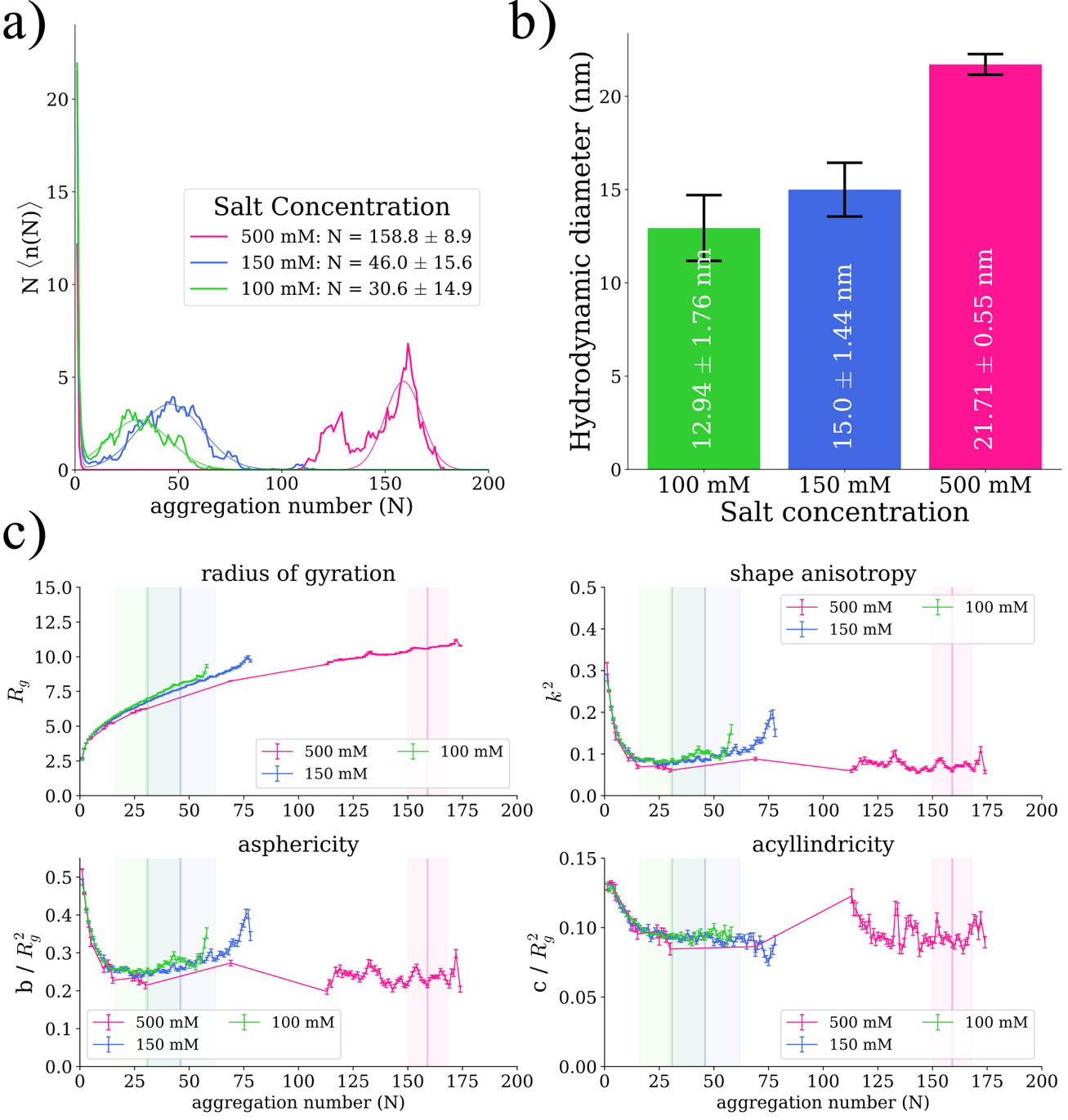

**Supplementary Figure 15. Effect of salt concentration on micellization of TDP-43 LCD 12 pS. a)** Size distribution (full lines) and corresponding fitting curves (dashed lines) for different salt concentrations. Mean aggregation number and standard deviation obtained from the fit are shown in legend. **b)** Mean hydrodynamic diameter at different salt concentrations. Results were obtained from a weighted average of the hydrodynamic radii values of clusters of sizes within one standard deviation of the mean aggregation number of the micelles obtained from the fit to the size distributions. Error bars represent the standard deviation of the distribution of hydrodynamic diameters. **c)** Shape descriptors obtained from the gyration tensor of individual assemblies as a function of aggregation number at different salt concentrations. Values are shown as the mean the standard error on the mean. The vertical lines represent the mean aggregation number obtained for the fits of micelle size distributions, the shaded region around it represents one standard deviation.

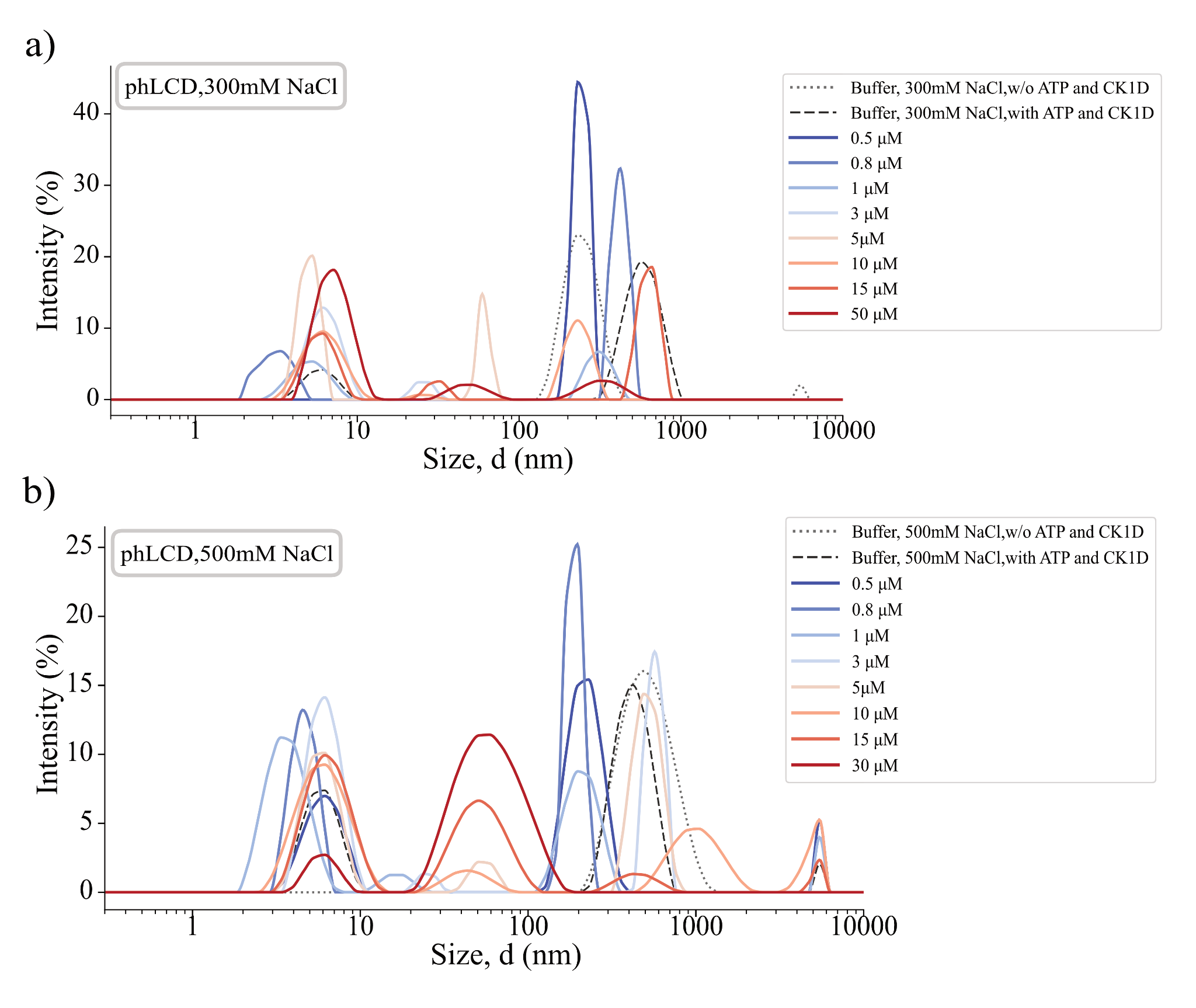

**Supplementary Figure 16. Concentration- and time-dependent DLS size distributions of phosphorylated TDP-43 LCD at increasing ionic strength.** Intensity-weighted hydrodynamic diameter distributions of phLCD at **a)** 300 mM NaCl and **b)** 500 mM NaCl, across a range of protein concentrations (0.5-50 µM for 300 mM; 0.5-30 µM for 500 mM). Buffer-only controls - with and without CK1δ and ATP - are shown as dashed and dotted lines, respectively. Large species >100 nm detected in both protein samples and buffer controls are attributed to background scattering and excluded from analysis. The temporal stability of the phLCD micellar population over multi-day timescales is independently confirmed in **Supplementary Figure 10**.

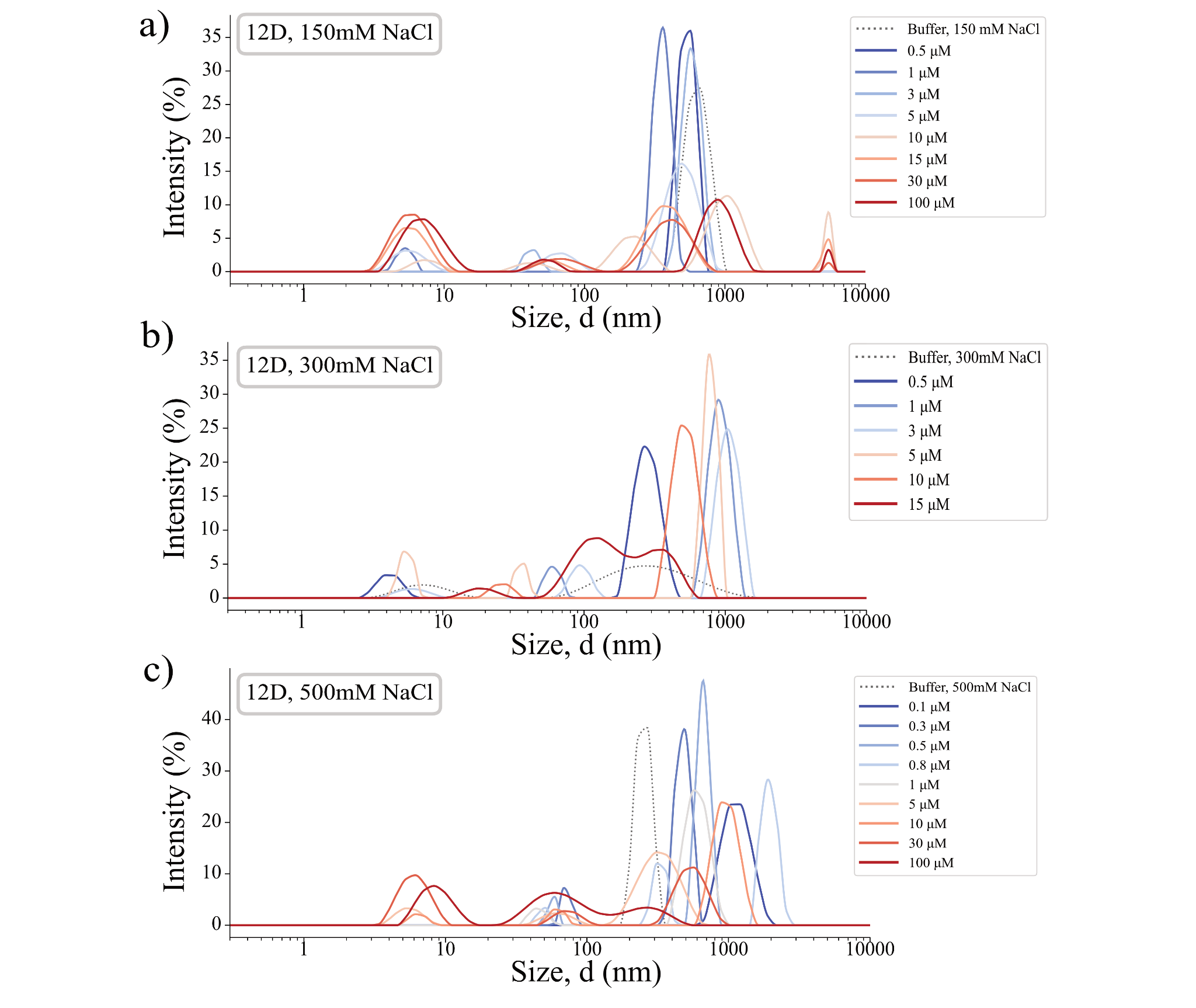
**Supplementary Figure 17. Concentration-dependent DLS size distributions of phosphomimetic 12D TDP-43 LCD at increasing ionic strength.** Intensity-weighted hydrodynamic diameter distributions of 12D LCD at **a**) 150 mM, **b**) 300 mM, and **c**) 500 mM NaCl across a range of protein concentrations. Buffer-only controls are shown as dotted lines.

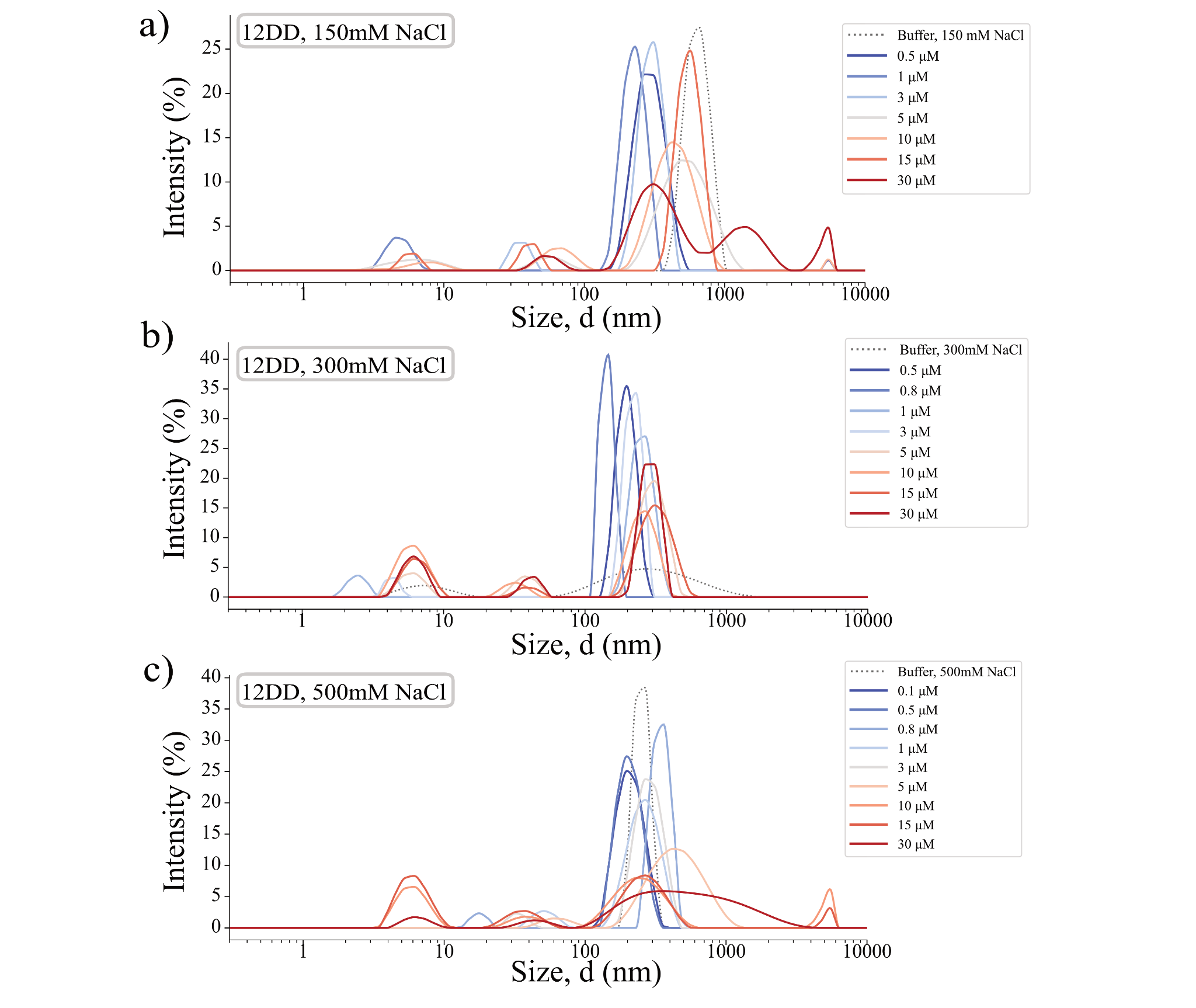

**Supplementary Figure 18. Concentration-dependent DLS size distributions of phosphomimetic 12DD TDP-43 LCD at increasing ionic strength.** Intensity-weighted hydrodynamic diameter distributions of 12DD LCD at **a**) 150 mM, **b**) 300 mM, and **c**) 500 mM NaCl across a range of protein concentrations. Buffer-only controls are shown as dotted lines.

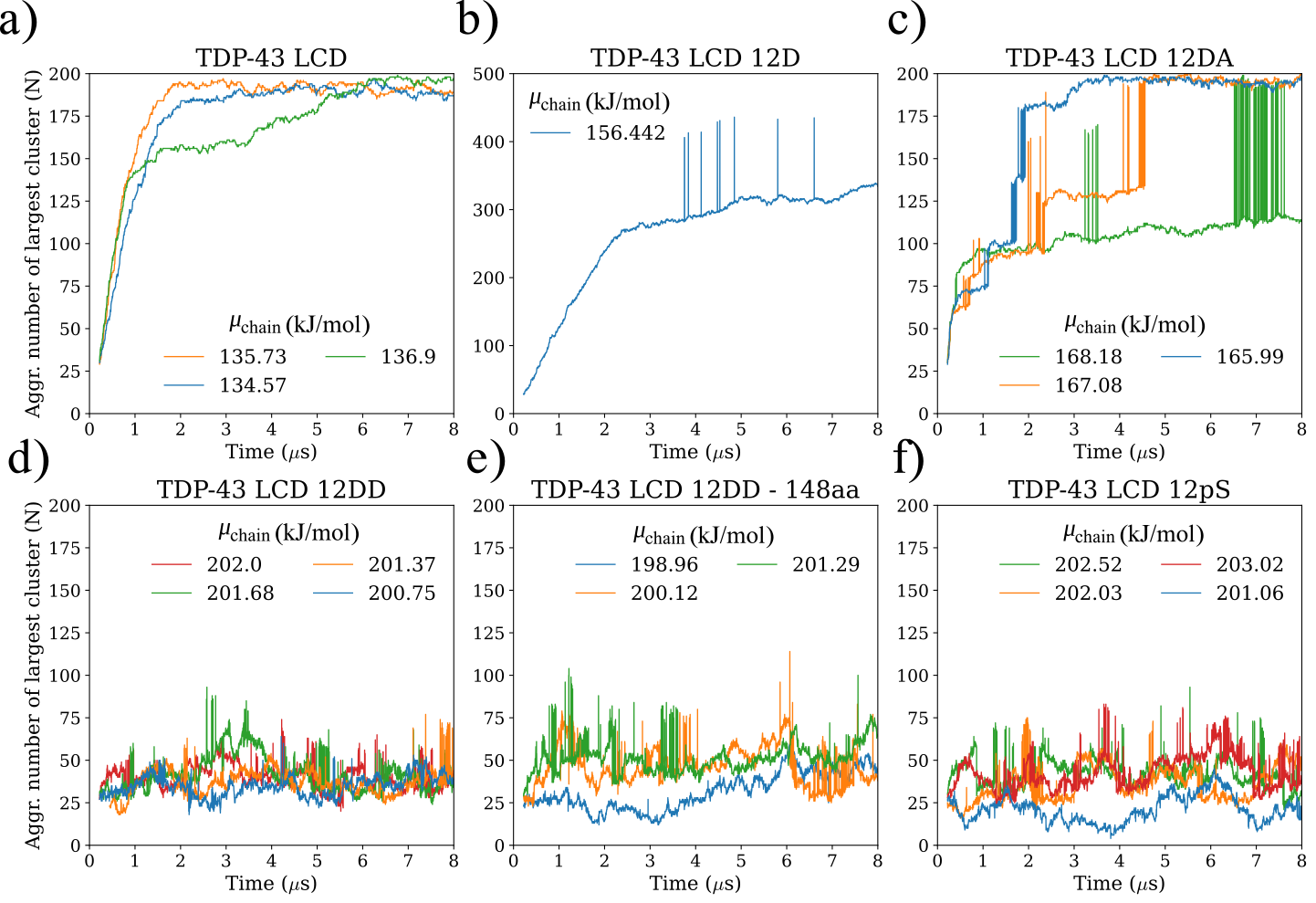

**Supplementary Figure 19. Aggregation number of the largest cluster in the simulation box as a function of time.** The size of the largest cluster grows from the initial seed and stabilizes around a mean value. The jumps that are sometimes observed are artifacts from the clustering algorithm. All data is extracted from simulations at 100 mM salt concentration. The sequence name is given on top of each subplot. Values in the legend indicate the chemical potential value per chain used in each SGCMC run.

*Validation of the Semi-Grand Canonical Monte Carlo simulations*

We validated our semi-grand canonical Monte-Carlo by comparing with equivalent MD simulations. We did so for all the 4 main sequences used across the manuscript.

***Phenotypes***

We confirmed that we observed the same phenotype (micellization or phase separation) in both methods for each sequence. Snapshots of the simulations are shown below and show identical phenotypes to those in Figure 2b.
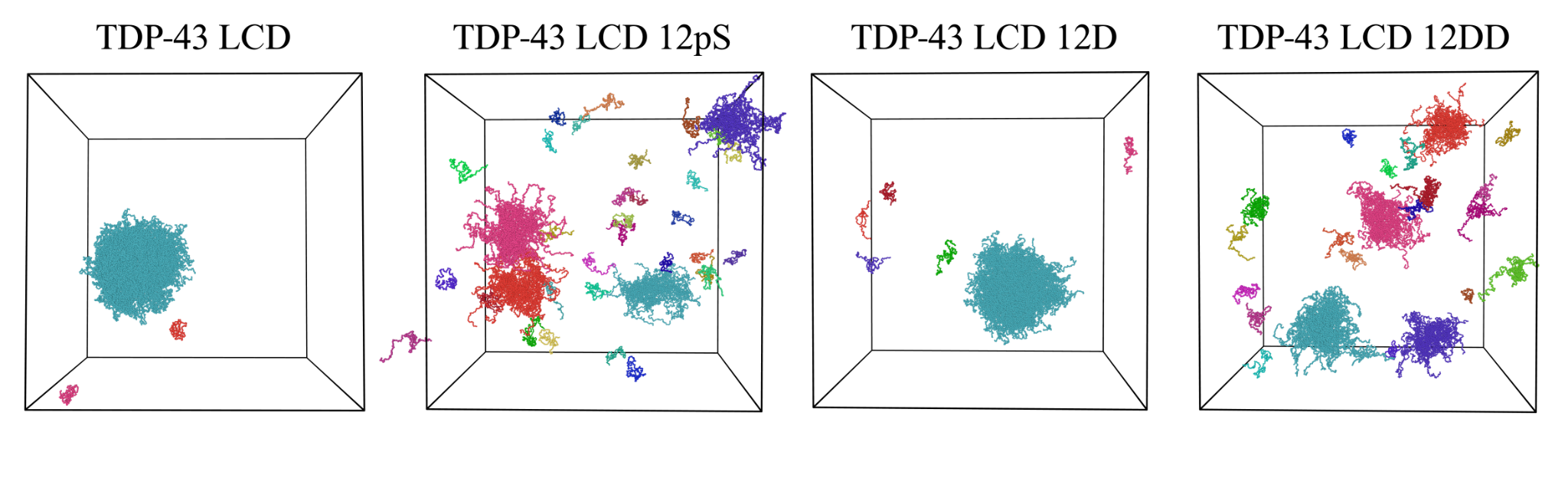

**Supplementary Figure 20. MD simulations yield identical phenotypes as SGC.** Snapshots of MD simulations for the 4 simulated sequences. Images made with OVITO.

***Size Distributions & Hydrodynamic Diameters***

We computed the size distribution histograms for the micellizing sequences and plotted them against each other. We observe nearly identical size distributions for both methods. We then compute the hydrodynamic diameters for micelles within the aggregation number interval given by the fit. They are nearly identical for SGCMC and MD simulations.

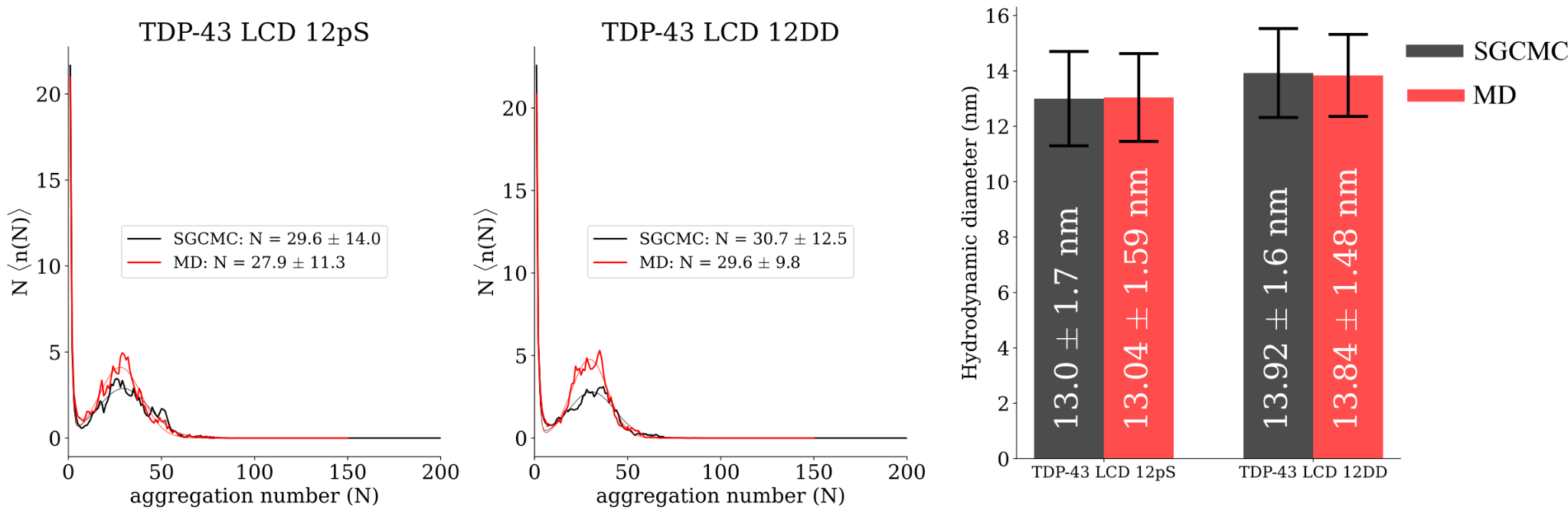

**Supplementary Figure 21. SGCMC simulations yield similar size distributions and hydrodynamic diameters as MD for micellizing sequences.** (Right) Size distribution histograms for both micellizing sequences. Black and red lines represent results from SGCMC and MD simulations. Full lines represent the histograms and dashed lines represent the fitted curves. (Left) Hydrodynamic diameters for both micellizing sequences. Error bars represent the standard deviation. Black and red bars represent results from SGCMC and MD simulations.

***Density Profiles***

We plotted per-block and whole-protein density profiles for both methods. We considered the same cluster size interval and we observe that they overlap significantly well. To quantify the degree of quantitative agreement we plot the densities of both methods at the same x coordinate on an xy plot, with results for SGCMC simulations in the x axis and those for MD in the y-axis. We see that for all blocks and for the whole-protein density profiles the points fall consistently on the diagonal line. Small deviations are observed at high densities for the phase-separating sequences, which is expected, since they represent regions close to the COM of the cluster, which have small volumes and are more prone to fluctuations in the number of particles. Further deviations from the diagonal in the phase-separating sequences seem to come from uneven sampling of cluster sizes. As can be seen in the size distribution plots, MD simulations explore a much more narrow interval of cluster sizes than SGCMC, which overrepresents larger cluster sizes which are more energetically favorable and shift the density profiles slightly to the right. For the micellizing sequence we observe near perfect quantitative agreement for density profiles. This is explained by the very good match between the size distribution histograms from both methods.

We quantified how well the points in the xy plot of densities fall on the diagonal line through the coefficient of determination ${R^{2}}$, defined by the formula:

$$R^{2} = 1 - \frac{\sum_{i} (y_{i}-x_{i})^{2}}{{\sum_{i} \left( y_{i}-\underline{y} \right)}^{2}}$$

where $x_{i}$ and $y_{i}$ are the paired data points, $\underline{y}$ is the mean of the $y_{i}$ values, the numerator represents the sum of squared deviations from the identity line y=x and the denominator is the total variance in the y values. Thus, $R^{2}$ quantifies how closely the data follow the diagonal parity line, with $R^{2}$ = 100% indicating perfect agreement. Despite the small deviations in the phase-separating systems we still observe $R^{2}$ > 99% for every single density profile and approaching 100% for the micellizing sequences. This indicates that SGCMC captures the internal architecture of the clusters observed in MD simulations.

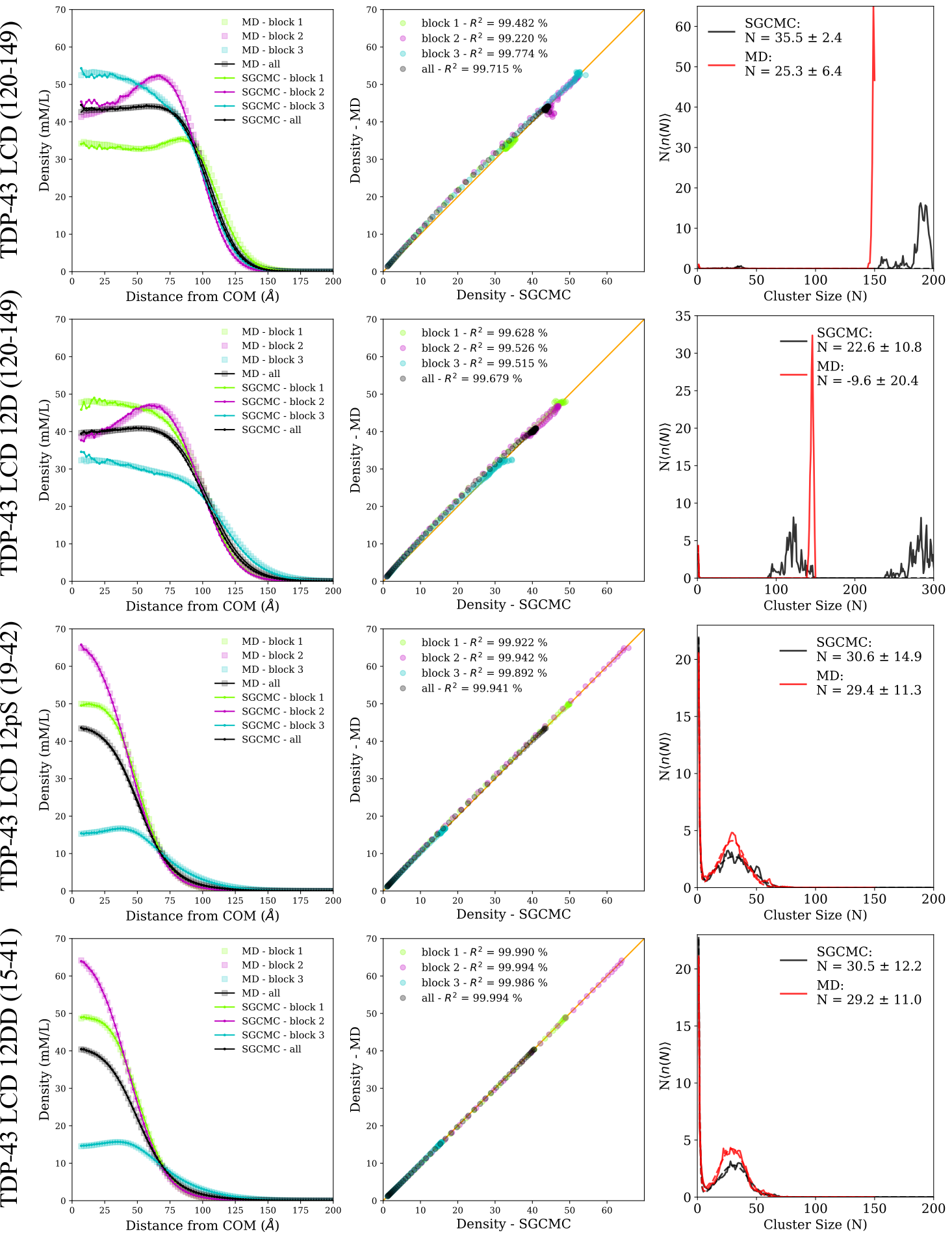

**Supplementary Figure 22. Density profiles obtained from MD and SGCMC simulations are almost identical.** Each line represents one of the 4 sequences. The aggregation number interval considered for each plot is given in parentheses on the y axis. (Left column) Density profiled for each of the 3 blocks and the whole sequence. Results obtained from MD are shown as square markers and results from SGCMC are shown as full lines with dot markers. (Middle column) Density profiles from MD and SGCMC plotted against each other in an xy plot. Full orange line represents the diagonal (y=x). Computed coefficient of determination ($R^{2}$) is given in the legend. (Right column) Size distribution histograms (full lines) and the corresponding fit (dashed lines) for SGCMC (black) and MD (red).

***Contact Maps:***

We compared the interchain contact maps obtained from both methods at the same cluster size intervals used in the validation for density profiles. We also observe very good agreement, with contact maps appearing indistinguishable to the eye in both methods. We observe tiny deviations in the mean number of contacts per residues within each block (lower diagonal part of the plot). We subtract the contact maps to compute $\Delta_{ij}= C_{ij}^{MD} - {C_{ij}^{SGCMC}}$. These results are plotted in the bottom row and we observe very small deviations, in order to make the plot visible we had to set a colorbar in the range (-0.005-0.005), whereas $C_{ij}$ ranges from 0 to 0.05, suggesting deviations are at least one order of magnitude smaller than the actual value of the number of contacts between each pair of residues. This indicates that the choice of simulation method also does not significantly affect the contact landscape between proteins.

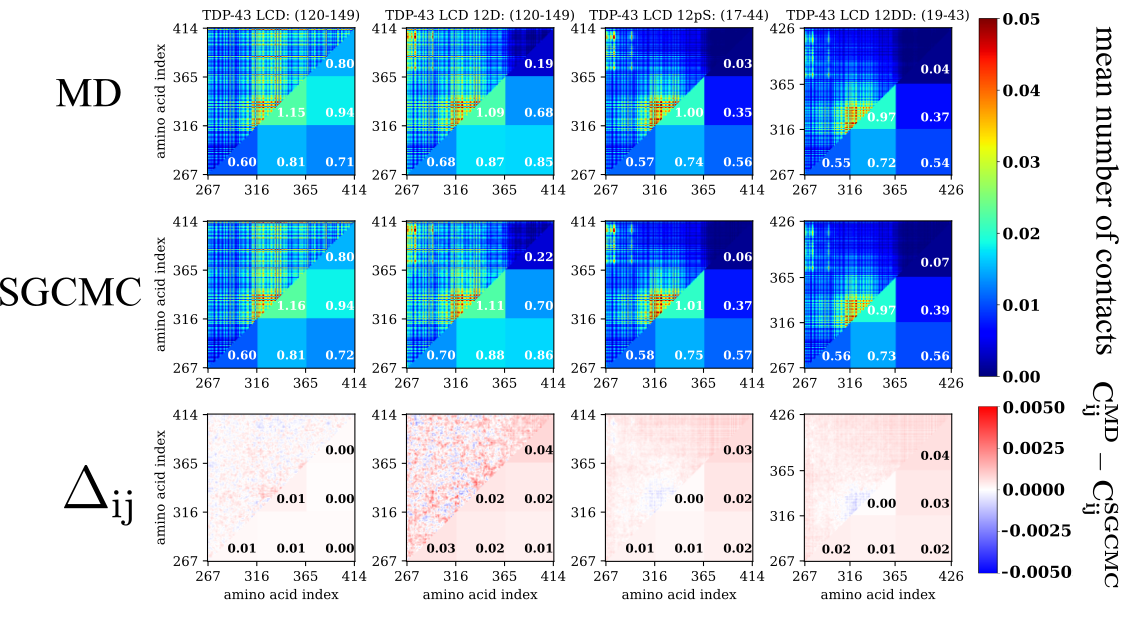

**Supplementary Figure 23. Interchains contact maps obtained from MD and SGCMC simulations are nearly identical.** Interchain contact maps from MD (top) and SGCMC (middle) simulations. The upper diagonal plot represents the mean number of contacts between each amino acid pair. Lower diagonal terms represent the mean number of contacts per amino acid between 2 blocks. Aggregation number interval is indicated at the top of each contact map. All simulations at 100 mM salt concentration. In the bottom row we show the subtraction between the contact maps from MD and SGCMC.

***Shape descriptors***

Finally, we compared the shape descriptors (radius of gyration, shape anisotropy, asphericity and acyllindricity) as a function of cluster size for each simulation method. As can be seen in the plots below, we observe qualitatively identical curves in the micellizing sequences, with deviations at cluster sizes significantly larger than the equilibrium aggregation number. In the phase-separating sequences we observe very good agreement for the cluster sizes which were observed in both methods. As discussed previously, MD simulations explore a narrower range of cluster sizes due to how they are initialized. These results show that both methods yield clusters with identical shape properties, indicating they do not affect bulk interactions and surface effects that determine the shape of the clusters.

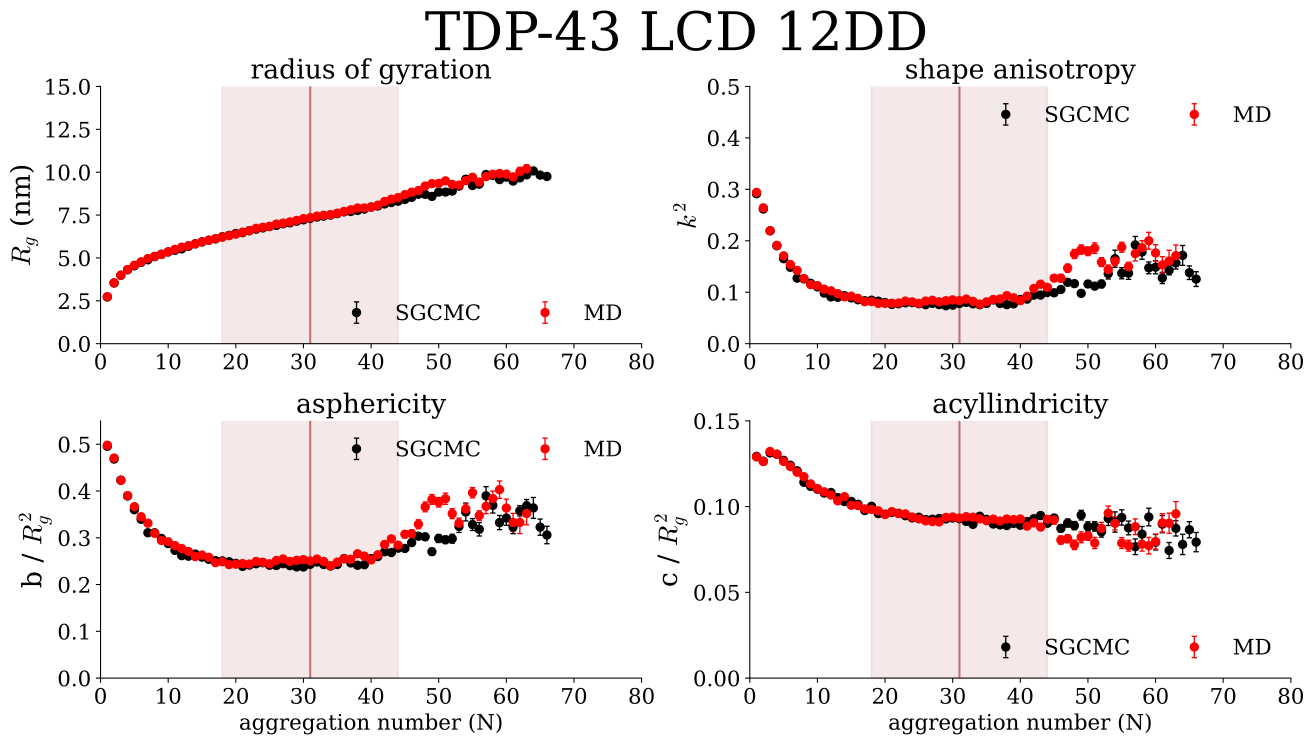

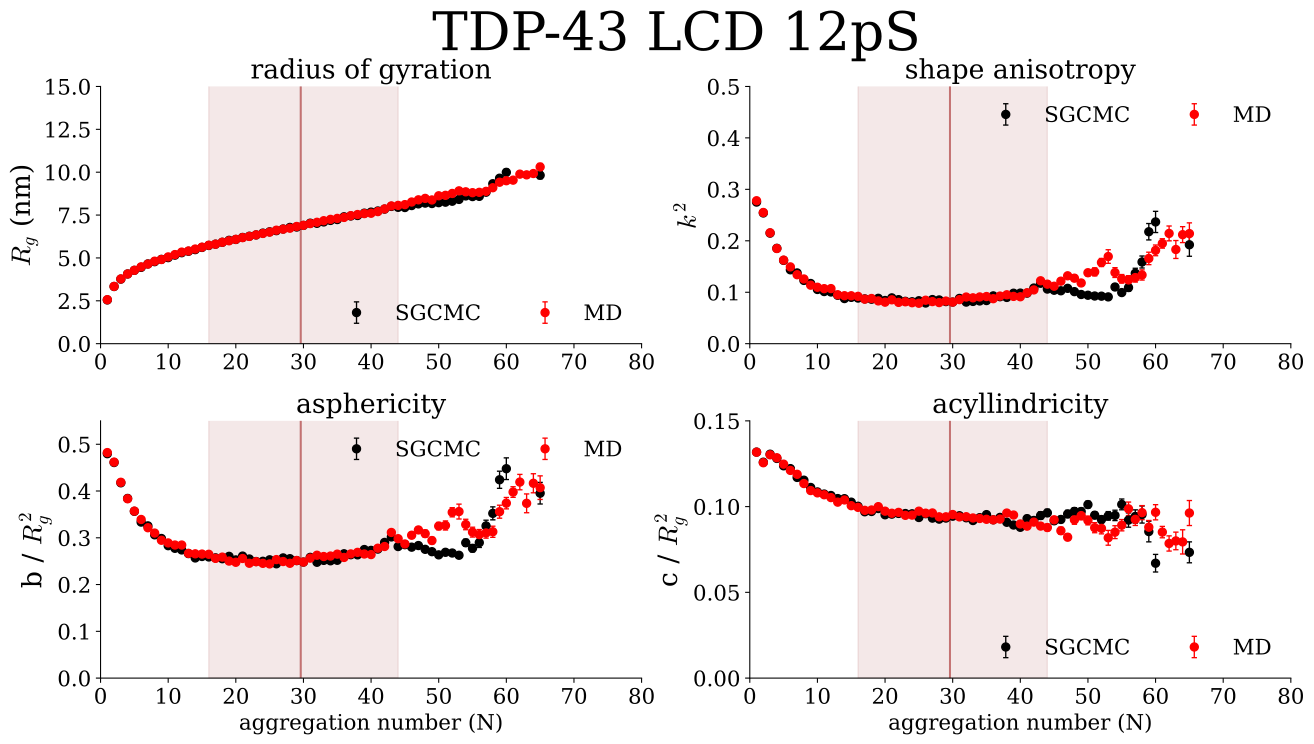

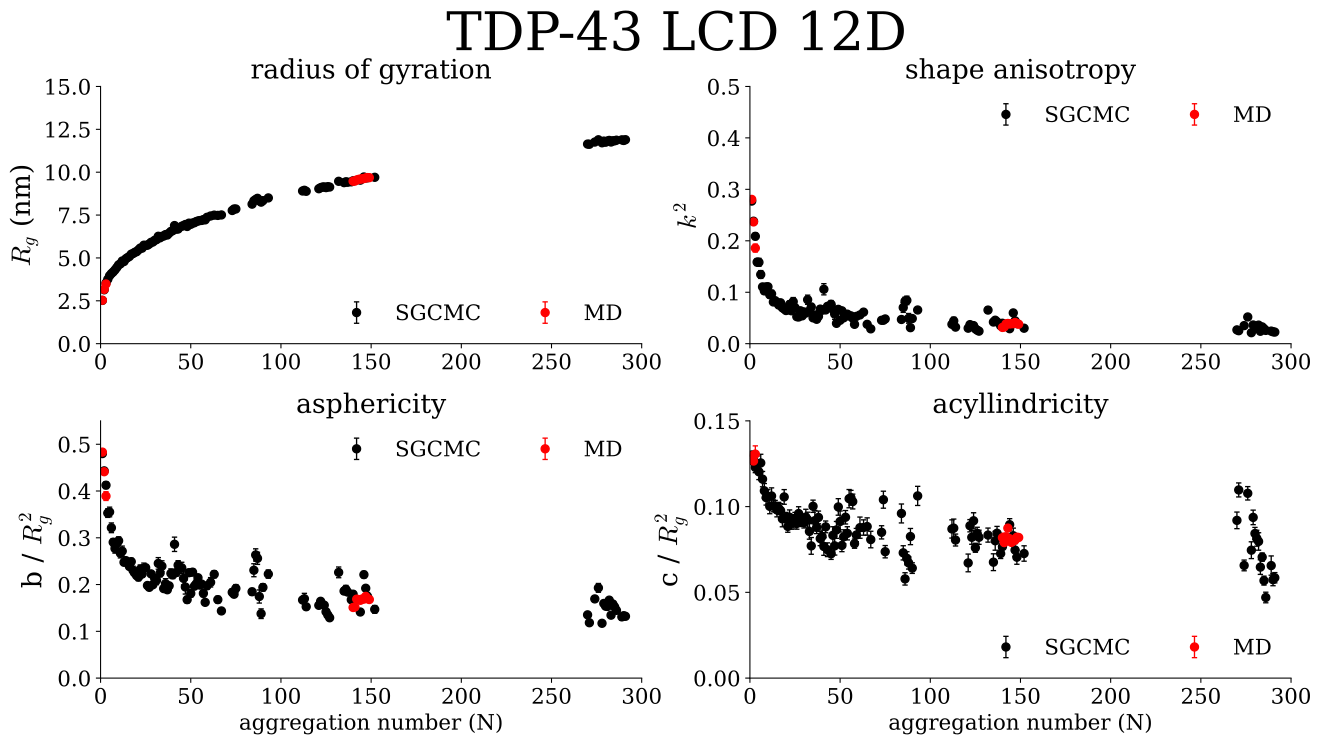

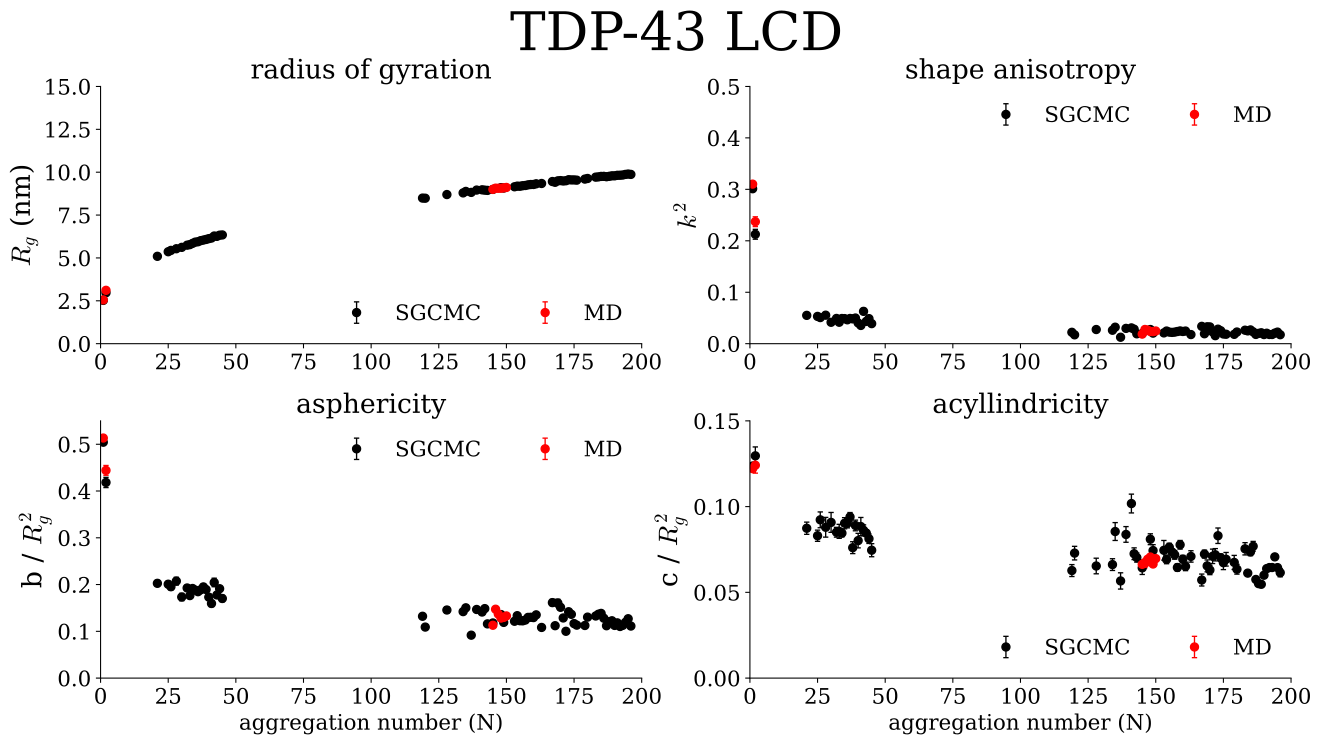

**Supplementary Figure 24. Shape descriptors obtained from MD and SGCMC are nearly identical.** Plots of the shape descriptors obtained from the gyration tensor for clusters of different aggregation numbers. Values are shown as mean $\pm$ the standard error on the mean. The vertical lines represent the mean aggregation number obtained for the fits of micelle size distributions, the shaded region around it represents one standard deviation.

We conclude that the results provided are more than enough to prove the validity of our SGCMC implementation. All relevant quantities used to describe the mechanisms behind phase separation and micellization in the 4 relevant sequences are nearly identical in MD and SGCMC simulations.
